# Immunotherapy of glioblastoma explants induces interferon-γ responses and spatial immune cell rearrangements in tumor center, but not periphery

**DOI:** 10.1101/2022.01.19.474897

**Authors:** Tala Shekarian, Carl P. Zinner, Ewelina M. Bartoszek, Wandrille Duchemin, Anna T. Wachnowicz, Sabrina Hogan, Manina M. Etter, Julia Flammer, Chiara Paganetti, Tomás A. Martins, Philip Schmassmann, Steven Zanganeh, Francois Le Goff, Manuele G. Muraro, Marie-Françoise Ritz, Darci Phillips, Salil S. Bhate, Graham L. Barlow, Garry P. Nolan, Christian M. Schürch, Gregor Hutter

## Abstract

Recent therapeutic strategies for glioblastoma (GBM) aim at targeting immune tumor microenvironment (iTME) components to induce antitumoral immunity. A patient-tailored, *ex vivo* drug testing and response analysis platform for GBM would facilitate personalized therapy planning, provide insights into treatment-induced immune mechanisms in the iTME, and enable the discovery of biomarkers of therapy response and resistance.

We cultured 47 GBM explants from tumor center and periphery from 7 patients in perfusion bioreactors to assess iTME responses to immunotherapy. Explants were exposed to antibodies blocking the immune checkpoints CD47, PD-1 or or their combination, and were analyzed by highly multiplexed microscopy (CODEX, <u>co</u>-detection by in<u>dex</u>ing) using an immune-focused 55-marker panel. Culture media were examined for changes of soluble factors including cytokines, chemokines and metabolites. CODEX enabled the spatially resolved identification and quantification of >850,000 single cells in explants, which were classified into 10 cell types by clustering. Explants from center and periphery differed significantly in their cell type composition, their levels of soluble factors, and their responses to immunotherapy. In a subset of explants, culture media displayed increased interferon-γ levels, which correlated with shifts in immune cell composition within specific tissue compartments, including the enrichment of CD4^+^ and CD8^+^ T cells within an adaptive immune compartment. Furthermore, significant differences in the expression levels of functional molecules in innate and adaptive immune cell types were found between explants responding or not to immunotherapy. In non-responder explants, T cells showed higher expression of PD-1, LAG-3, TIM-3 and VISTA, whereas in responders, macrophages and microglia showed higher cathepsin D levels. Our study demonstrates that *ex vivo* immunotherapy of GBM explants enables an active antitumoral immune response within the tumor center in a subset of patients, and provides a framework for multidimensional personalized assessment of tumor response to immunotherapy.

## Introduction

Glioblastoma (GBM) is a fatal brain tumor without effective treatment options. Current standard of care consists of gross total surgical resection followed by chemo-radiation, resulting in a mean overall survival of 14 months [35]. Recently, strategies harnessing the non-neoplastic immune tumor microenvironment (iTME) have evolved. A major problem in GBM therapy lies in the inherent immunosuppression exerted by the cell types residing in the iTME. The GBM iTME consists of myeloid-derived macrophages and yolk sac-derived microglia (collectively termed glioma-associated macrophages/microglia, or GAMs) as well as other myeloid cell types and lymphocytes. GAMs infiltrate into GBM tumors and, by interaction with tumor cells, change their functional state towards an immunosuppressive and regenerative phenotype [1]. The composition of GAMs within GBM, their origin and phenotypic evolution during tumorigenesis are currently under intense research [5, 25]. Importantly, GBM is a heterogeneous and widely invasive malignancy, and human data on the composition of the iTME and its interaction with neoplastic cells in different tumor regions (contrast medium-enhancing tumor center, peripheral infiltration zone) are scarce [9]. Recently, major efforts have been undertaken to describe the GBM iTME on the single-cell level, highlighting the predominance of myeloid cells [11, 20].

Immune checkpoint inhibitor therapies with anti-CTLA-4 and/or anti-PD-1/PD-L1 antibodies have revolutionized the treatment of many solid tumors. However, so far, results from clinical trials of systemic T cell checkpoint blockade in GBM were disappointing [26]. Other studies suggest that PD-1 blockade as a neoadjuvant treatment in combination with adjuvant maintenance therapy could increase survival compared to adjuvant PD-1 blockade alone [7]. High PD-L1 expression levels are associated with decreased survival in GBM patients [7]. The expression of PD-L1 and indoleamine 2,3-dioxygenase (IDO-1), as well as the accumulation of regulatory T cells (Tregs) in response to the presence of CD8^+^ T cells are known mechanisms of adaptive resistance [24] and might counteract immune responses. Combinatorial immunotherapies addressing both innate and adaptive immune cell types of the iTME may therefore circumvent these resistance mechanisms [32].

Previously, we focused on the disruption of the CD47-Sirpα axis to regain antitumor activity of GAMs against malignant brain tumors. We and others showed that Sirpα itself is a potent modulator of macrophage-mediated phagocytosis [23, 36]. Blocking the CD47-Sirpα axis suppresses a “don’t eat me” signal to macrophages and allows for efficient phagocytosis of most cancers tested so far [39, 30, 31]. Preclinical analysis of a humanized anti-CD47 antibody demonstrated potent *in vitro* and *in vivo* tumor killing ability against GBM [12, 42]. Anti-CD47 treatment induced a macrophage polarization change of GAMs *in vivo*. Furthermore, both M1 and M2-polarized macrophages displayed a higher GBM cell phagocytosis rate under anti-CD47 treatment, indicating that even M2 macrophages can be rendered phagocytic [42]. Moreover, anti-CD47 treatment induced exclusive microglia-mediated GBM cell phagocytosis in a xenograft mouse model, when macrophage influx was impeded [17].

The culture of intact tumor tissues is an attractive strategy to assess the effects of cancer immunotherapies on the iTME [38]. To dissect the composition of the human GBM iTME and its response to innate and adaptive immune checkpoint inhibitor therapy *ex vivo*, we used 3D tissue perfusion bioreactors, CO-Detection by indEXing (CODEX) highly multiplexed microscopy, soluble protein arrays and mass spectrometry. Here, we present an in-depth analysis of the GBM iTME of intra-operatively annotated samples from tumor center and periphery at baseline and after 7 days of *ex vivo* immunotherapy targeting CD47 and/or PD-1. We included the peripheral invasion zone since most tumor recurrences originate from this region. Hence, targeting the periphery after surgical tumor control by immunotherapy would be an important pillar in adjuvant GBM treatment. Our approach mimics a localized application of immunotherapy, which poses a clinically feasible modality with less systemic toxicity. We found that a subset of explants from the tumor center was responsive to the tested immunotherapies. Moreover, we provide evidence of intratumoral re-activation of T cells and relief of immunosuppression using immunotherapies targeting both innate and adaptive immune cell types in a subset of responding tumors. This approach provides a multidimensional proof-of-concept strategy to identify patients who might be amenable to local immunotherapeutic approaches.

## Materials and Methods

### GBM tissue collection and patient characteristics

Biopsy samples from contrast medium-enhancing tumor center and non-enhancing tumor periphery were collected during surgery via intraoperative navigation. From each patient, 5-aminolevulinic acid (5-ALA) positive, vital tumor center biopsies as well as biopsies from the 5-ALA low/negative tumor periphery/infiltration zone were obtained. Tissue acquisition was documented by intraoperative imaging (taking into consideration 5-ALA positivity and neuronavigation, Figure S1). In total, 7 tumor samples from *isocitrate dehydrogenase 1/2* wild type primary GBM were included in this study. One patient (ID 588) was pretreated with temozolomide and radiation therapy before resection. Patient characteristics, survival data, molecular and histopathological data are summarized in **Figure 1B** and Table S1. MRI images from the localizations of patients’ tumor biopsies are shown in Figure S1. Written informed consent was obtained from all patients, and the study was approved by and conducted according to the guidelines of the local ethics committee (Ethikkommission Nordwestschweiz, #42/10).

**Figure 1.**
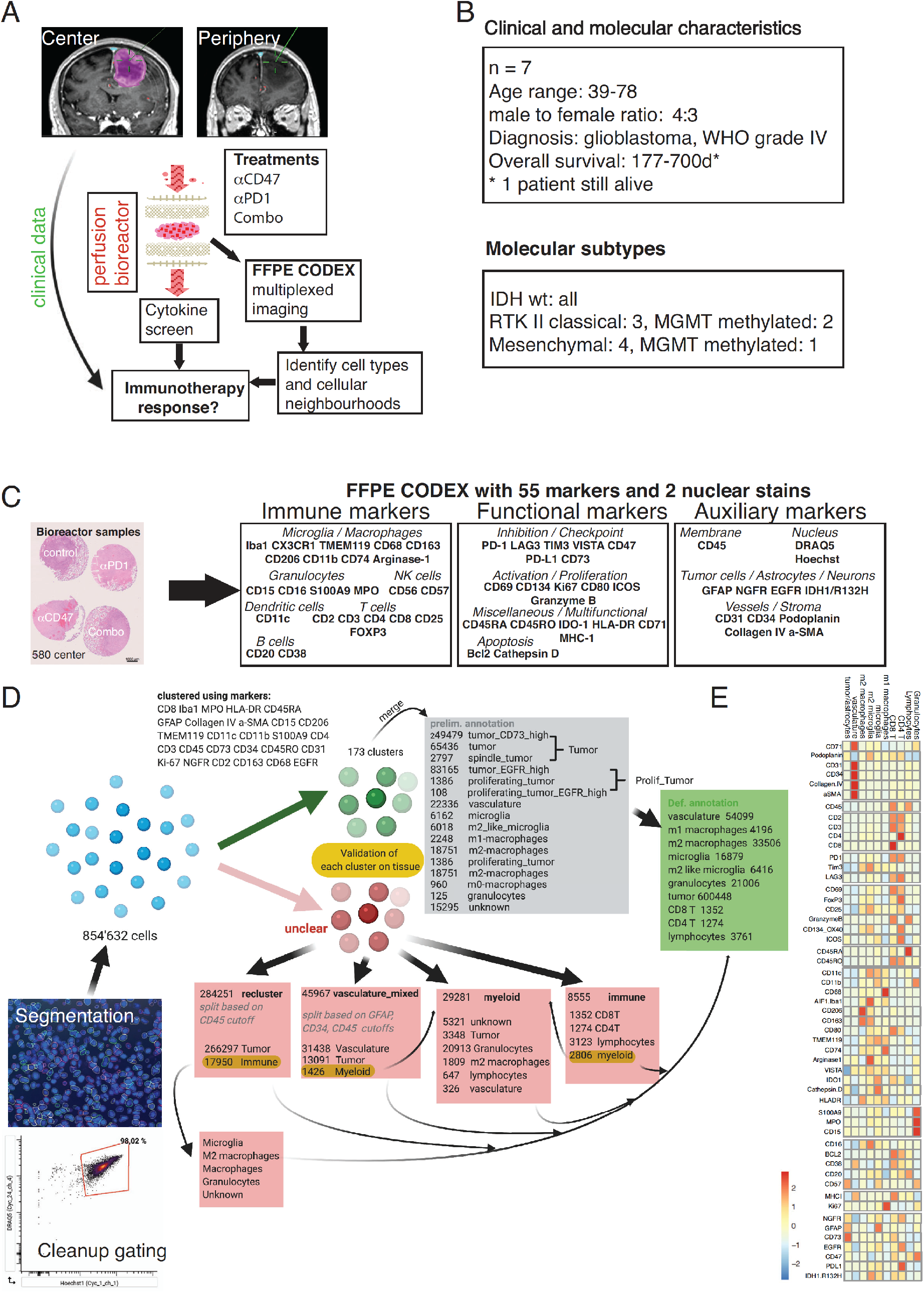
Experimental setup, clinical characteristics, and CODEX clustering pipeline. **(A)** Experimental setup of the proof-of-concept study: Fresh tumor biopsies were taken according to neuronavigation and directly transferred into 3D perfusion bioreactors. Immunomodulatory treatments consisted of innate (anti-CD47) and/or adaptive (anti-PD1) immune checkpoint inhibition for 7 days. Soluble proteins and metabolites from bioreactor media, and multidimensional CODEX microscopy data were integrated to assess immunotherapy response per patient/tumor region sample. **(B)** Clinical baseline and genetic data of the included patients (n=7). **(C)** Left panel: Representative arrangement of an FFPE processed, H&E-stained center biopsy consisting of explants cultured in the bioreactors and treated with different immunotherapies. Scale bar: 1000 μm. Right panel: FFPE CODEX antibody panel. **(D)** Clustering strategy for cell type annotation. Cells were segmented based on nuclear DRAQ5 staining, and DRAQ5/Hoechst double-positive cells were gated in CellEngine (lower left panel), resulting in 854’632 single cells across all samples. Vortex-Clustering at first using the markers specified led to 173 clusters, which were subsequently merged/simplified based on manual validation on the tissue. Unclear cells were reclustered, re-annotated, and re-validated on each tissue section using the cell finder package. Finally, 10 main cell types throughout all samples could be annotated with high confidence (cf. **Fig S3**). **(E)** Heatmap of mean fluorescence intensities of individual markers used in the CODEX pipeline per annotated cell type across all samples.

### *Ex vivo* perfusion bioreactor cultures

Bioreactor cultures under perfusion were performed according to the manufacturer’s instructions (Cellec Biotek AG, Basel, Switzerland). Briefly, fresh, intact GBM tissue explants were placed into ice-cold neurobasal-A medium (Life Technologies, #21103049) and immediately taken to the laboratory (less than 30 min). Tissue from the center and periphery of the tumor were cut into ∼20-30 mm^3^ fragments. This size was selected based on preliminary experiments, which showed improved survival with limited cutting-related damage. Tissue fragments were placed between two 10 mm diameter discs of microfibrillar collagen hemostat sheets (Ultrafoam, Avitene) pre-wet with PBS for 1 h at 37 °C, 5% CO_2_. Silicone adaptors and ETFE mesh grids were arranged on the top and bottom of the collagen scaffold and the tissues were placed into U-CUP perfusion chambers (Cellec Biotek AG). The perfusion media consisted of a 50:50 mix of Neurobasal-A medium (Life Technologies, #21103049) and DMEM/F12 medium (Gibco, #21331020) supplemented with non-essential amino acids (1x, Sigma, #M7145), 1 mM sodium pyruvate (Sigma, #S8636), 44 mM sodium bicarbonate (Gibco, #5080094), 25 mM HEPES (Gibco, #156301), 4 mM L-alanyl-L-glutamine (Corning, #25-015-CI), antibiotic-antimycotic (1x, Gibco, #LS15240062), 20 ng/mL human recombinant EGF (Gibco, #PHG0314), 20 ng/mL human recombinant FGF (Gibco, #AA10-155), 10 ng/mL heparin sulfate (Stemcell Technologies, #07980) and 5% human serum (Sigma-Aldrich, #H4522-100ML). The perfusion flow rate was set at 0.47 mL/min, resulting in a superficial flow velocity of 100 µm/s. Starting from day 0, bioreactor media were either left untreated or supplemented with the anti-PD-1 antibody nivolumab (Opdivo^TM^, Bristol Myers Squibb, 5 µg/mL), anti-CD47 antibody (BioXCell, µg/mL), alone or in combination. On day 7, culture media were frozen at −80°C and tissues were fixed in 10% neutral buffered formalin (Sigma, #HT501128) for 24 h and processed according to standard histopathology procedures for paraffin embedding [13].

### Generation of CODEX DNA-conjugated antibodies

Purified, carrier-free monoclonal and polyclonal antibodies (Table S2) were conjugated to short DNA oligonucleotides (Trilink Biotechnologies) as described before [4, 19, 29]. Briefly, antibodies were concentrated on 50 kDa molecular weight cut-off centrifugal filter columns (Amicon Ultra, EMD Millipore, #UFC505096), and sulfhydryl groups were activated using a mixture of 2.5 mM TCEP (Sigma, #C4706-10G) and 2.5 mM EDTA (Sigma, #93302) in PBS, pH 7.0, for 30 min at room temperature. Toluene-deprotected, lyophilized, maleimide-modified DNA oligonucleotides were then conjugated to the antibodies at a 2:1 weight/weight ratio for 2 h, with at least 100 μg of antibody per reaction. Conjugated antibodies were washed, eluted in PBS-based antibody stabilizer (Thermo Fisher, #NC0436689) containing 5 mM EDTA and 0.1% NaN_3_ (Sigma, #S8032), and stored at 4 °C.

### CODEX antibody validation, titration and staining

Validation and titration of the CODEX antibodies, as well as the stainings of formalin-fixed, paraffin-embedded (FFPE) GBM bioreactor tissues were performed as described before [4, 29]. Briefly, DNA-conjugated antibodies were first validated on tonsil tissues or a multi-tumor tissue microarray under the supervision of a board-certified surgical pathologist (C.M.S.), and staining patterns were confirmed with online databases (The Human Protein Atlas, www.proteinatlas.org [37]; Pathology Outlines, www.pathologyoutlines.com) and the published literature. GBM bioreactor tissues were punched out of their respective paraffin blocks using 4 mm diameter punches, assembled into arrays, sectioned at 4 μ (Electron Microscopy Sciences, #72204-01) pre-treated with Vectabond^Ⓡ^ (Vector Labs, #SP-1800) according to the manufacturer’s instructions. Coverslips were baked at 70 °C for 1 h, deparaffinized, and heat-induced epitope retrieval was performed in a LabVision^TM^ PT module (Thermo Fisher) using Dako target retrieval solution, pH 9 (Agilent, #S236784-2) at 97 °C for 10 min. After blocking for 1 h at room temperature, 100 μ coverslip and staining was performed in a sealed humidity chamber overnight at 4 °C on a shaker, followed by washing and fixation in 1.6 % paraformaldehyde (10 min, room temperature), 100% methanol (5 min, 4 °C), and BS3 (Thermo Fisher, #21580; 20 min, room temperature).

### CODEX multi-cycle reaction and image acquisition

CODEX multi-cycle reactions and image acquisition were performed as described before [4, 29]. Briefly, stained coverslips were mounted onto custom-made acrylic plates (Bayview Plastic Solutions) using coverslip mounting gaskets (Qintay, #TMG-22), and the tissue was stained with Hoechst 33342 (Thermo Fisher, #62249; 1:1000). Acrylic plates were mounted onto a custom-designed plate holder and secured onto the stage of a BZ-X710 inverted fluorescence microscope (Keyence). Fluorescent oligonucleotides (concentration: 400 nM) were aliquoted in black 96-well plates (Thermo Fisher, #07-200-762) in 250 μ nuclear stain (1:600) and 0.5 mg/ml sheared salmon sperm DNA (Thermo Fisher, #AM9680). DRAQ5 nuclear stain (Cell Signaling Technologies, #4084L) was added to the last well at a dilution of 1:100. For details on the order of fluorescent oligonucleotides and microscope light exposure times, see Table S2. Automated image acquisition and fluidics exchange were performed using a CODEX instrument and driver software (Akoya Biosciences) according to the manufacturer’s instructions, with slight modifications. Tissue overview images were acquired manually using a CFI Plan Apo λ 2x/0.10 objective (Nikon), and automated imaging was performed using a CFI Plan Apo λ 20x/0.75 objective (Nikon). After each multi-cycle reaction, hematoxylin and eosin (H&E)-stainings were performed according to standard pathology procedures, and tissues were reimaged in brightfield mode. Staining quality, marker expression and distribution was verified on each individual section according to [4, 29].

### Computational image processing

Raw TIFF image files were processed, deconvolved and background-subtracted using the CODEX Toolkit uploader and Microvolution software (Microvolution) as described before [4, 14, 29]. Antibody stainings were visually assessed for each channel and cycle in each spot using ImageJ software (Fiji, version 2.0.0). Final figures of cell passports and composites for main cell clusters were generated using OMERO.web app (https://www.openmicroscopy.org/omero/figure/).

### Cell segmentation

TIFF hyperstacks were segmented based on DRAQ5 nuclear stain, pixel intensities were quantified, and spatial fluorescence compensation was performed using the CODEX Toolkit segmenter as described previously [29], using the following settings: Radius: 7. Max. cutoff: 1.0. Min. cutoff: 0.07. Relative cutoff: 0.2. Cell size cutoff factor: 0.4. Nuclear stain channel: 4. Nuclear stain cycle: 23. Membrane stain channel: 1. Membrane stain cycle: −1 (i.e., not used). Use membrane: false. Inner ring size: 1.0. Delaunay graph: false. Anisotropic region growth: false. This generated comma-separated value (CSV) and flow cytometry standard (FCS) files for further downstream analysis.

### Cleanup gating, unsupervised hierarchical clustering and cluster validation

All background-subtracted FCS files were imported into CellEngine (https://cellengine.com). Gates were tailored for each file individually in a blinded manner by two experts in flow and mass cytometry (T.S. and C.M.S.). Nucleated cells were positively identified, and artifacts were removed by gating on Hoechst1/DRAQ5 double-positive cells, followed by gating on focused cells in the Z plane. After cleanup gating, FCS files were re-exported and subsequently imported into VorteX clustering software, where they were subjected to unsupervised hierarchical X-shift clustering using an angular distance algorithm [27]. The following data import settings were applied: Numerical transformation: none. Noise threshold: no. Feature rescaling: none. Normalization: none. Merge all files into one dataset: yes. Clustering was based on the following antibody markers: CD8, Iba1, MPO, HLA-DR, CD45RA, GFAP, Collagen IV, CD206, TMEM119, CD11c, CD11b, S100A9, CD4, CD3, CD45, CD73, CD34, CD45RO, CD31, Ki-67, NGFR, CD2, CD163, CD68, EGFR. The following settings were used for clustering: Distance measure: Angular distance. Clustering algorithm: Xshift (gradient assignment). Density estimate: N nearest neighbors (fast). Number of neighbors for density estimate (K): From 150 to 5, steps 30. Number of neighbors: determine automatically. The optimal cluster number was determined using the elbow point validation tool and was at K=40, resulting in 173 clusters. Clusters and corresponding data were exported as a CSV file and were manually verified and assigned to cell types by overlaying the single cells from each individual cluster onto the stitched bioreactor images in ImageJ, based on the unique cluster identifiers and cellular X/Y position, using custom-made ImageJ scripts (available at https://github.com/bmyury/CODEX-fiji-scripts). Clusters with similar morphological appearance in the tissue and similar marker expression profiles were merged, and artifacts were removed. Unclear clusters were reclustered based on the markers highlighted in **Figure 1D**, re-checked individually on the stitched tissue slides, and merged back to the already annotated cell types, resulting in 10 final clusters.

### CODEX marker expression analysis

Data normalization was achieved by log-transformed median normalization of each marker in each sample to their respective baseline control sample to focus on expression changes and allow comparisons between samples. Differences in median expression were assessed using the non-parametric Kruskal-Wallis H-test [21], using Benjamini-Hochberg procedure to control the false discovery rate.

### CytoMAP spatial analysis to identify tissue compartments (TCs)

CytoMAP [34] was written using MATLAB version 2018b (Mathworks). A detailed description of the workflow and functions built into CytoMAP is available in the online user manual (https://gitlab.com/gernerlab/cytomap/-/wikis/home). Below is a brief discussion of the analysis used for the datasets described in this manuscript. The annotated cell types for each dataset were loaded into CytoMAP by importing the corresponding CSV files. These collectively contain the cell type annotations derived from the final clustering, mean fluorescent pixel intensity values per cell for each marker, and spatial positions (centroids) for each cell object. Once imported, the following functions were used in CytoMAP to analyze the data.

### Raster Scan Neighborhoods

This function identifies the local composition of cell types within a circular area in the tissue, which we here refer to as “tissue compartment” (TC). This function calculates the number of cells and the mean fluorescent intensity of each channel summed over all cells in each TC. The positions of the TCs are evenly distributed throughout the tissue in a grid pattern with a distance between TC centers of half of the user defined radius (r=75 μ The TC information was used for further analysis (e.g., local cellular densities, cell-cell associations).

### Cell-Cell Correlation Analysis

The local cell density within individual TCs was used to correlate the location of different cell types, revealing which cell populations preferentially associate with one another, or conversely avoid one another. This function calculates the Pearson correlation coefficient of the number of cell or object types within the scanned TCs and graphs these on a heatmap plot. This correlation analysis can be performed across multiple samples, and can be done either over entire tissues or within specified tissue regions. This is important, as cells may have distinct associations with one another in different tissue compartments.

Analysis of TC frequencies and distributions among samples TC prevalence and cell type composition per sample were exported and analyzed per treatment condition or location.

### Multiplexed secreted protein analysis in explant culture media

Bioreactor media were centrifuged to remove cell debris, and 92 secreted proteins, including cytokines, chemokines and soluble cell membrane proteins were measured externally by proximity extension technology using the Olink immuno-oncology panel (http://www.olink.com/products/immuno-oncology). Data are presented as normalized protein expression values, Olink Proteomics’ arbitrary unit on a log_2_ scale. Missing data were associated with a lower median expression. They were imputed as either half the molecule detection threshold or such that the sum of all imputed values for a molecule is 0.1 of the sum of the molecule’s expressions (whichever is the smallest). The principal component analysis was performed on centered and scaled data.

### Determination of kynurenine in explant culture media by liquid chromatography-mass spectrometry (LC-MS)

Stock solutions of kynurenine (Sigma) and D6-kynurenine (Cambridge Isotope Laboratories Inc.) were prepared at 5 and 10 mM in water. A series of seven standard solutions for the calibration curve were prepared at concentrations from 5000 nM to 39 nM by serial dilution. D6-kynurenine was diluted at 600 nM in LC-MS grade methanol (Sigma). Supernatants and calibration solutions (5 µL) were quenched with ice cold methanol containing the internal standard (D6-kynurenine at 600 nM). Samples were mixed and centrifuged at 3700 g for 15 min. 20 µL of ultrapure water (supplied by a MilliQ Advantage A10 purification system, Merck Millipore) was added prior to injection and the total dilution was x10. HPLC was set up to inject directly from the supernatant 8 mm above the pellet. ***Instrumentation and chromatographic conditions:*** measurements were performed on a HPLC Nexera X2 HPLC (Shimadzu) coupled to an API 5500^TM^ (AB Sciex) with positive ion electrospray ionization and operated in multiple reaction monitoring (MRM) mode. Data were collected and processed by Analyst^®^ 1.6.2 and the chromatographic separation was carried out on a Acquity HSS T3 1.8 m, 2.1×50 mm (Waters Corp) at 40 °C. The separation method consisted in a gradient elution of the mobile phase (0.1% formic acid in A: water and B: acetonitrile) as follows: 0% B from 0-0.25 minutes, 10% B at 0.3 minutes, 15% B at 1.1 minutes, 90% B at 1.2 minutes and held for 0.3 minutes before re-equilibration. Total run time was 2 minutes and all samples were analyzed with an injection of 2mL. The source parameters were curtain gas (CUR), nitrogen at 20 PSIG, collision gas at 6 PSIG, ion source gas-1 at 20 PSIG; ion source gas-2 at 20 PSIG, ion spray voltage (IS) at 4500 V, turbo heater temperature (TEM) at 500 °C and entrance potential (EP) at 10 V. The electrospray ionization was operated in positive multiple reaction monitoring mode (MRM) after optimization according to standard procedure. Compound specific values (mass transitions, declustering potential and collision energy) of mass spectrometer parameters were as follows for the two analytes: Kynurenine: Q1 (m/z) 209.05; Q3 (m/z) 192.05; DP (V) 43.0; CE (V) 12.5. D6-kynurenine: Q1 (m/z) 215.05; Q3 (m/z) 198.05; DP (V) 40.0; CE (V) 14.0.

### Statistical analysis

Data are reported as mean ± s.d. or mean ± s.e.m., and statistical significance was determined using the Mann-Whitney *U*-test, two-tailed Wilcoxon test, student’s t-test or Kruskal-Wallis test using Prism v9.0e (GraphPad), as specified in the respective figure legends. Differences were considered statistically significant if *P* displayed. Correlations were evaluated using Pearson correlation. Computational analyses were performed in R v.4.0.2, CytoMAP, Prism v9.0e, or python 3.8.8 (using packages: scipy1.7.1, numpy 1.21.1, statsmodels 0.12.2, pandas 1.3.1, scikit-learn 0.24.1, matplotlib 3.4.2, seaborn 0.11.1). Experiments were performed without duplicates because of material restrictions. The immunotherapy response based on IFNγ cytokine measurements was assessed by first calculating the receiver-operating characteristic (ROC) curves based on the delta values (immunotherapy treated condition minus untreated condition) for each parameter measured. Parameters that were strongly associated with IFNγ were selected based on the area under the curve (AUC)-ROC curve (**Figure S10A**).

## Results

### *Ex vivo* culture and immunotherapy of intact GBM explants from tumor center and periphery using perfusion bioreactors

To profile the region-specific GBM iTME by highly multiplexed microscopy and assess treatment responses to immune checkpoint inhibitors, we prospectively collected GBM specimens from 7 patients undergoing neuronavigated surgery (**Figure 1A** and Table S1). From each tumor, contrast medium enhancing, 5-aminolevulinic acid (5-ALA) positive, vital tissue from the center as well as 5-ALA low/negative tissue from the periphery / infiltration zone was acquired. Tissue acquisition was documented by intraoperative imaging (taking into consideration 5-ALA positivity and neuronavigation, Figure S1). After neuropathological examination by two board-certified neuropathologists, all samples were diagnosed as GBM, WHO grade IV, *isocitrate dehydrogenase 1/2 (IDH)* wild type. Further, GBM subtype analysis was performed by whole genome methylation analysis according to Capper et al. [6], documenting three receptor tyrosine kinase RTK II (classical) and four mesenchymal subtypes (**Figure 1B** and Table S1).

Intact tumor fragments (explants) were subsequently cultured in 3D tissue perfusion bioreactors [16]. This culture system provides flow of the media through the tissue, which enables culturing intact tissues of greater thickness and overcomes typical limitations of static cultures, including limited transport of nutrients and waste removal, particularly in the tissue center. We validated and optimized this technology and found superior tissue preservation in perfused cultures compared to non-perfused / static conditions (Figure S2). Specifically, an intact iTME, tumor cell proliferation, and invasion of GBM cells into the scaffold could be detected in explants cultured for up to three weeks (Figure S2).

To simulate a continuous local immunotherapy, we treated the perfused explants for 7 days with CNS-adapted doses [10] of an antibody against the microglia-macrophage checkpoint CD47 [40], with the clinically approved PD-1 checkpoint inhibitor nivolumab [28], or a combination of both (**Figure 1A**). Control samples were left untreated. To investigate potential differential effects of these immunotherapies on tumor center vs. periphery, we treated patient-matched samples from both regions simultaneously.

Our results indicate that intact GBM tissues from tumor center and periphery can be cultured alive for multiple days and provide a framework for GBM-tailored immunotherapy testing using multimodal analysis of the iTME.

### FFPE CODEX enables interrogating changes in iTME composition and architecture in GBM explants treated with immunotherapy

To analyze changes in immune cell type abundance and localization in the GBM iTME induced by immunotherapies, we performed highly multiplexed microscopy using CODEX on a total of 47 explants [29]. We selected and validated a panel of 55 antibodies for GBM iTME-specific protein markers on FFPE sections, which included markers for identifying immune cell types, functional markers (costimulatory proteins, immune checkpoints and molecules involved in apoptosis and proliferation), as well as auxiliary markers for tumor, vascular and stromal cells (**Figure 1C**, Tables S2 and S3).

After thorough validation of staining quality, marker expression and distribution in the tissue sections, iTME cell types were identified using a combined approach of unsupervised clustering, manual cluster refinement and by overlaying the clusters onto each individual tissue for morphological confirmation (**Figures 1D and** S3). Only markers that had a robust expression (moderate or strong signal intensity) throughout all the samples were used for clustering (see **Methods**). To validate the clustering result, we generated ‘cell passports’ for each cell type based on the expression of cell type-defining markers and appropriate negative controls (Figure S3). For all cell types, standardized mean fluorescence intensities across all the markers in the panel are shown in **Figure 1E**.

Tissue integrity after 7 days of perfusion culture was well preserved, and perfused samples were comparable to fresh tissues based on hematoxylin & eosin (H&E) staining (**Figure 2A** and data not shown). Using composite multicolor overlays derived from the CODEX dataset, we visualized each of the 10 annotated, final cell types in the tissue context based on phenotype-defining markers and morphology (**Figure 2B**). Vasculature-defining markers were CD34, CD31, and collagen IV. Tumor cells and astrocytes were positive for GFAP and NGFR, with a subset of proliferating, Ki-67^+^ tumor cells. CD45, CD68, CD11b, Iba1 and HLA-DR were used to identify macrophages and microglia. CD163 was used as a surrogate marker of an M2-like state for both microglia and macrophages; microglia and M2-like microglia were discriminated from macrophages by TMEM119 positivity. Granulocytes represented another major myeloid population, and were characterized by MPO, CD11b, S100A9 and CD15 expression. In line with previous studies [20], the adaptive immune compartment was relatively low-abundant in our sample set, and was dominated by CD3^+^CD4^+^ and CD3^+^CD8^+^ T cells and a CD45 high-expressing, morphologically defined lymphocytic population negative for T cell markers, which we called “lymphocytes” (Figure S3).

**Figure 2.**
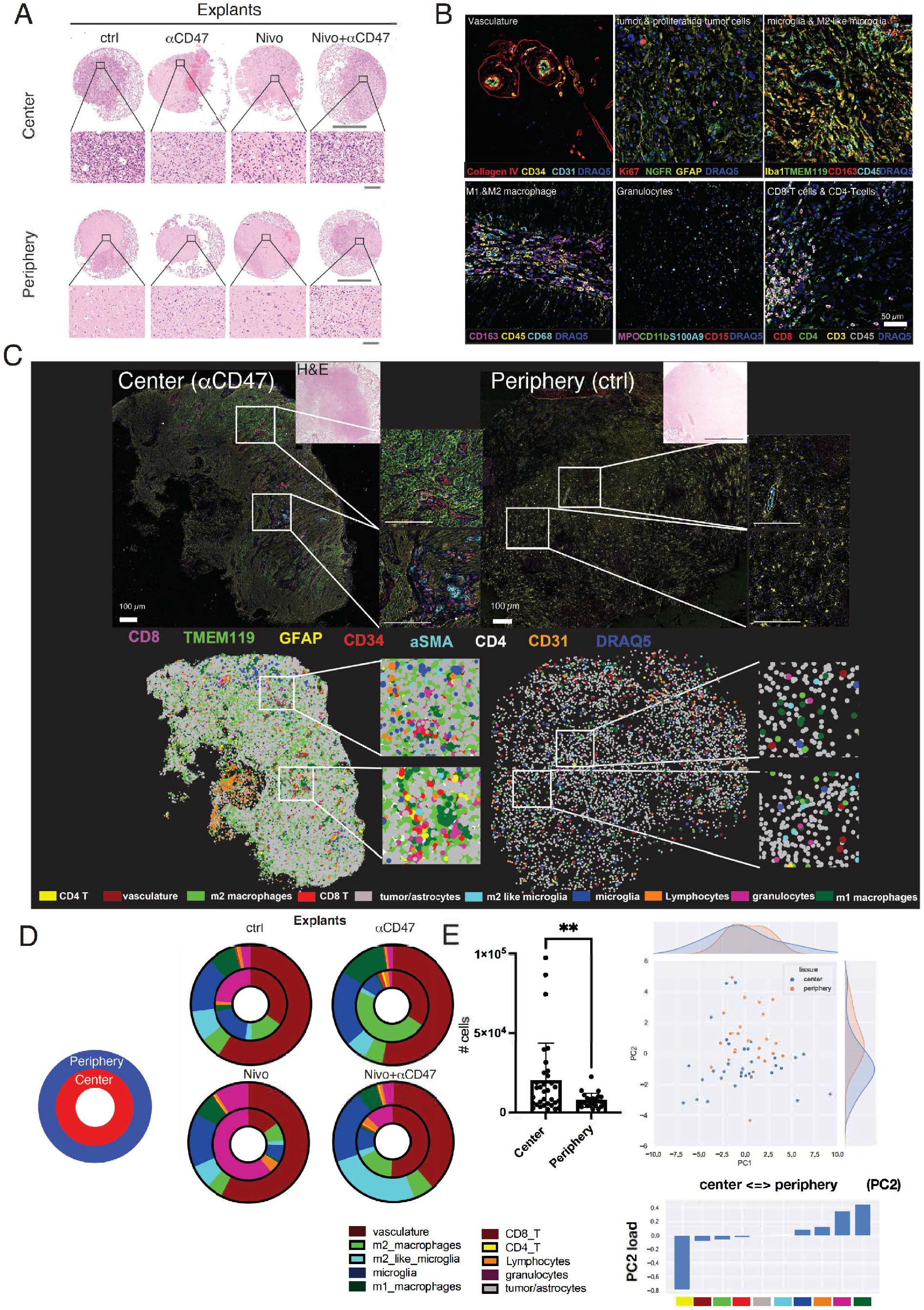
Differential composition of cellular phenotypes across biopsy locations. **(A)** Representative H&E-stained images of FFPE GBM center and periphery explants after 7d of culture in perfusion bioreactors. Scale bars: 1000 μm (overview), and 50 μm (close-up). **(B)** Multiplexed immunofluorescence overlay images of the most important cell types and their defining markers after definitive cell type annotation. **(C)** Upper-panel: FFPE-CODEX stained representative GBM bioreactor sample from center and periphery (tumor ID 588). 7 out of 54 markers are depicted indicating microglia (TMEM119^+^), tumor cells/reactive astrocytes (GFAP^+^), CD8 T cells (CD8^+^), CD4 T cells (CD4^+^) and vasculature (CD31^+^CD34^+^ SMA^+^). Scale bars: 100 μm. *Lower panel:* Assembly of annotated cell types according to x-y coordinates of the corresponding tissue sample in the upper panel. **(D)** Cell type frequency distribution throughout all center and periphery samples used in the experiment. Inner circle: center samples, outer circle: periphery samples. **(E) Left:** overall cellularity among all bioreactor samples in center vs. periphery (n=31 center samples, mean 20995 cells/sample, n=22 periphery samples, mean = 8467 cells/sample; p=0.0095, unpaired t-test with Welch’s correction). Individual cellular abundances are visualized in Figure S6 and Figure 5G; **right:** PCA correlating abundances in center vs. periphery samples (across pooled bioreactor samples). Statistics: (E) Student’s t-test; \*\**P* < 0.01.

To obtain an overview of the overall tissue composition with respect to the 10 cell type clusters identified, we used a color-coded graphical representation, and correlated this to the morphology and multicolor overlays (**Figure 2C**). We observed great intratumoral heterogeneity in explants from the tumor center, exemplified by islands of pronounced microglia accumulation in parts of an anti-CD47 treated center explant, and a perivascular enrichment of CD4^+^ and CD8^+^ T cells (**Figure 2C**, left lower panel, insets). In an untreated explant from the tumor periphery, we observed the typical stellate morphology of astrocytes with homogeneously intermingled microglia, and only few adaptive immune cells (**Figure 2C**, right panels). By spatially arranging the annotated, color-coded clusters based on their X-Y coordinates, differences in cellular density and spatial distribution between the exemplary center and periphery explants can be appreciated (**Figure 2C**, lower panels).

### Cell frequencies and composition differ between explants from tumor center and periphery

We observed differences in the cell type composition and density of explants from tumor center and periphery (**Figure 2D** and Figures S4**-6**). Consistent with our biopsy selection, the total cell density in explants from the tumor center was significantly higher compared to that in explants from the tumor periphery (mean total cell counts per center explant: 20,779, periphery explant: 8,467, p=0.0095, **Figures 2C, lower panels; 2E and** Figures S4**-5**). Moreover, tumor center explants contained significantly higher numbers of CD4^+^ T cells (p=0.004) and lymphocytes (p=0.0055), and tended towards higher M2 macrophage (p=0.11) numbers. Conversely, M2-like microglia (p=0.0014) and M1 macrophages (p=0.00019) were more abundant in periphery explants (Figure S5). After performing a principal component analysis (PCA) of the relative abundance of the annotated cell types across samples, we observed marked differences between explants from tumor center and periphery (**Figure 2E**). The first principal component (PC1) did not significantly differ between the biopsy locations. Notably, the second principal component (PC2), which was more prevalent in tumor center explants (p=0.0041), had a positive weight for CD4^+^ T cells, CD8^+^ T cells, vasculature and M2 macrophages (**Figure 2E**).

### Spatial analysis identifies GBM iTME tissue compartments conserved across samples and patients

To further investigate whether immunotherapies affect the spatial localization of cell types in the GBM iTME, and whether this depends on the type of tumor sample (center vs. periphery), we identified GBM tissue compartments (TCs) using a raster-scanned radius method [34]. This analysis resulted in 7 distinct TCs that were conserved across samples (**Figure 3A**). The cell type composition of each TC, as well as the abundance of the different TCs across all samples, are visualized as a heatmap in **Figure 3B**, and their frequencies across explants from center and periphery are depicted in **Figure 3C**. As expected, the most prevalent TC (TC1) was composed mainly of tumor cells / astrocytes, and therefore corresponded to the “*bulk tumor/astrocyte compartment*” (**Figures 3B-C, TC1, dark blue**). TC2 was enriched in immunosuppressive cell types such as M2-like microglia and granulocytes, but also M1-macrophages; we therefore named this TC the “*myeloid compartment*“ (**Figures 3B-C, TC2, cyan**). TC3, the second most abundant compartment in all samples, consisted of a mixture of tumor cells and immune cells; we therefore named this compartment the “*tumor-immune interface*” (**Figures 3B-C, TC3, green**). TC4 morphologically corresponded to GBM-typical “*glomeruloid vascular proliferations*” (**Figures 3B-C, TC1, dark blue**, and Figure S3). CD4^+^ and CD8^+^ T cells, lymphocytes and M2 macrophages were enriched in TC5 (“*adaptive immune compartment*”) (**Figure 3B, TC5, orange**), while microglia defined TC6 (“*microglia-enriched compartment*”) (**Figure 3B, TC6, red**). Finally, TC7 was a vascular compartment with immune cell enrichment, which we termed “*vascular-immune interface*” (**Figure 3B, TC7, brown**).

**Figure 3.**
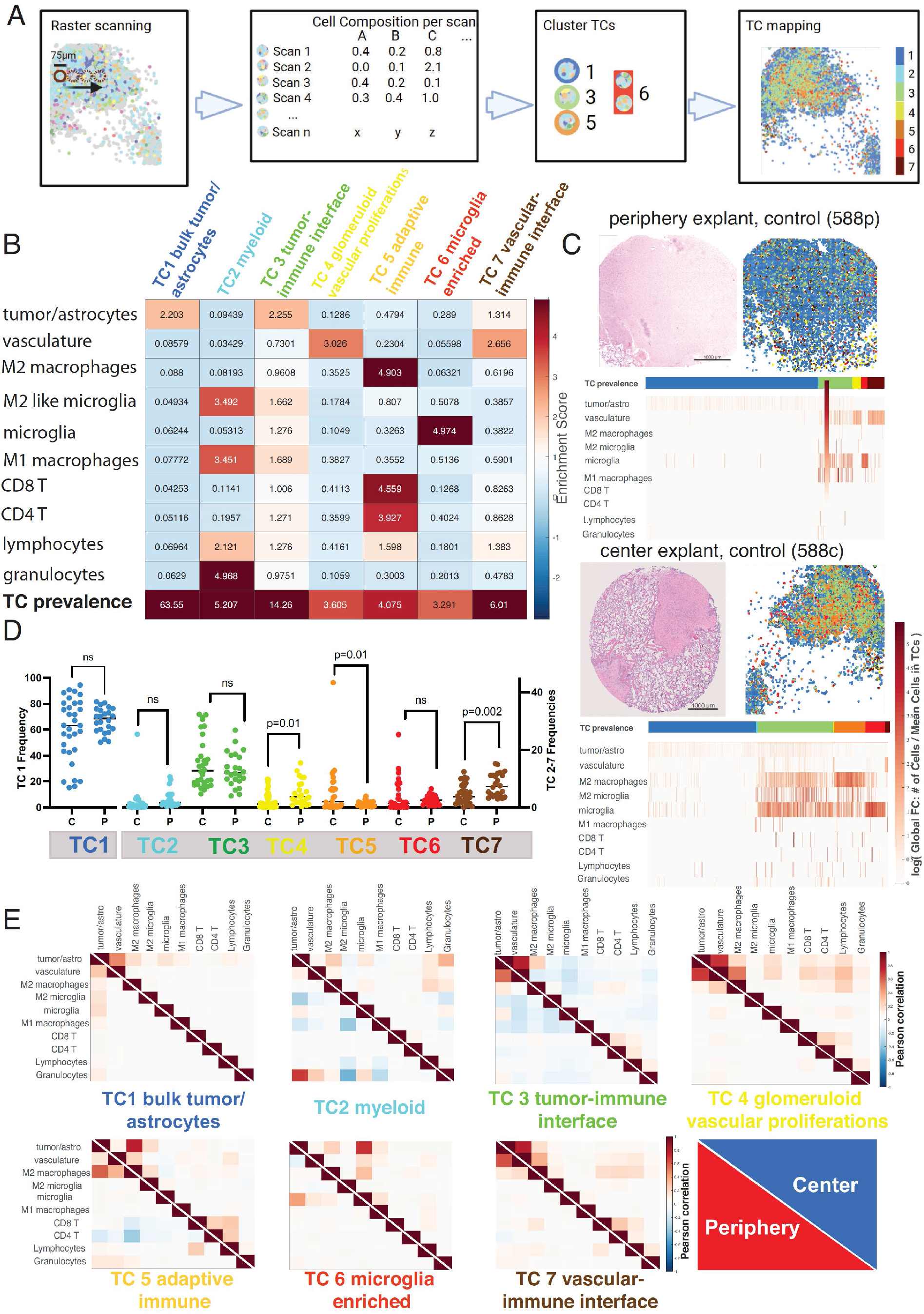
GBM explants can be subclassified into distinct tissue compartments which differ depending on biopsy location. **(A)** Schematic of neighborhood tissue compartment (TC) identification: Tissue regions were m diameter (best focus area). The cellular composition of each scan was recorded, and subsequently clustered using the NN self-organizing map (SOM) algorithm and the Davies-Bouldin criterion. This resulted in 7 distinct neighborhoods. **(B)** Region enrichment score depicting the fold change of cellular composition in each region, and their overall prevalence throughout all explants. Color code for TCs is used for all subsequent analysis. TCs were named based on their most prevalent cell types and functionality: TC1 (blue): bulk tumor-astrocyte compartment; TC2 (cyan): myeloid compartment; TC3 (green), tumor-immune interface; TC4 (yellow): glomeruloid vascular proliferations; TC5 (orange): adaptive immune compartment; TC6 (red): microglia enriched compartment; TC7 (brown): vascular-immune interface. **(C)** Representative TC map on bioreactor tissue from untreated periphery (upper image) and center (bottom image). H&E stained tissue section is supplemented by its corresponding TC overlay. Individual region heatmaps display the prevalence of TCs as well as the cellular composition of each region (log of cell number/total cells per TC). **(D)** Frequencies of TCs in all bioreactor samples depending on biopsy location (center C vs. periphery P). Each data point represents one bioreactor sample, and the horizontal line the median of all samples. Statistical analysis: Student’s t-test. **(E)** Normalized cell-cell correlation per TC in periphery (red, lower triangle) and center (blue, upper triangle). Pearson correlation coefficients (−1 to 1) between individual cell types per region are plotted. Reading example: in TC6, microglia and M2 macrophages have a stronger spatial interaction with tumor cells in the center than in the periphery.

### GBM iTME tissue compartment distribution points towards higher adaptive immune activity in tumor center explants

To visualize the different TCs and their spatial contexts in the tissue, we overlaid them onto the respective tissue samples (**Figure 3C** and Figure S4). In samples from the tumor periphery, we observed, as expected, a lower cell density in bioreactor explants examined, and a more homogenous distribution of the TCs (**Figure 3C**, *upper panel*). In contrast, in samples from the tumor center, higher cell densities and a more heterogenous TC distribution prevailed, and the region heatmap analyses showed a more pronounced infiltration by adaptive and especially innate immune cells, corresponding mainly to the presence of the respective TCs “M2 macrophages”, “M2-like microglia’’ and “microglia” (**Figure 3C**, *lower panel*). We performed this analysis for each patient sample, biopsy location and treatment condition, cross-comparing H&E stainings, cell density plots, cell type and TC localizations and cell type enrichments (Figure S6). We then analyzed TC frequencies between all bioreactor center (n=34) and periphery (n=24) samples independently of the applied treatment. The frequencies of TC4 (“glomeruloid vascular proliferations”), TC5 (“adaptive immune compartment”), and TC7 (“vascular-immune interface”) significantly differed in their frequency between tumor center and periphery explants (**Figure 3D**).

The higher frequency of TC5 in tumor center samples suggest a higher activity of adaptive immunity there. Interestingly, vascular TCs (TCs 4 and 7) were more abundant in the tumor periphery, which could portend neovascularization at the tumor invasive front [8]. When attempting to stratify the composition of the individual TCs according to individual treatments and location of the biopsies, no clear trends of composition changes were noticed (Figure S7**, A-B**).

### Cell-cell correlations within TCs have distinct patterns depending on biopsy location

Next, we computed cell-cell correlations for each TC across all explants, comparing tumor center and periphery. These relationships were quantified using the Pearson correlation coefficients of the numbers of cells per TC (**Figure 3E**). Such cell-cell spatial correlation analyses indicate cell populations preferentially co-localizing in the same TC. Notable differences of cell-cell correlation were observed in certain TCs. For example, adaptive immune cells correlated more strongly with vasculature in TC4 (*“glomeruloid vascular proliferations”*) in samples from the tumor center compared to those from the periphery (**Figure 3E, TC4, center**). This indicates a higher lymphoid cell infiltration of vascular proliferates in the tumor center. Furthermore, CD4^+^ T cells negatively correlated with tumor cells, vasculature and M2 macrophages in TC5 (*“adaptive immune compartment”*) in samples from the periphery, suggesting a blunted CD4^+^ T cell response at the tumor invasive front (**Figure 3E, TC5, periphery**). In TC6 (*“microglia-enriched compartment”*), microglia and M2 macrophages correlated more strongly with tumor cells in samples from the tumor center, as did M2 microglia and M1 macrophages (**Figure 3E, TC6, center**). These correlations point towards a stronger GAM-mediated immune response within the tumor center.

In summary, our spatial and cell-cell correlation analyses comparing explants from tumor center and periphery suggest that innate and adaptive anti-tumoral immune responses are enriched within the tumor center, whereas neovascularization and vasculature-associated processes are more prominent at the tumor periphery. This could have implications for designing locally targeted immunotherapies and antiangiogenic therapies, especially for tumor recurrences.

### Patient-personalized immunotherapy assessment by integrating explant-specific TC immune cell enrichment and secreted protein profiles

To pinpoint the effects of immunotherapies targeting innate and adaptive immune cell types in our patient-personalized model, we integrated the multidimensional information from both spatial imaging data and soluble factor profiles after 7 days of bioreactor treatment. Despite carefully navigated biopsies, explants from patient 580’s tumor periphery contained areas with higher cell density, whereas patient 583’s center explants partly showed a lower cell density. This indicates a greater regional heterogeneity in these tumors, which was not fully captured by 5-ALA neuronavigation (Figure S4**, page 3**). Moreover, tumor heterogeneity was prominent across patients, was higher in tumor center than periphery, but was relatively low within each patient’s center and periphery explants in the different treatment conditions (Figure S6). For example, in patient 587, we observed a relatively homogeneous distribution of cell densities in all tumor center explants (**Figure 4A**), whereas some variation was seen in the explants from other patients (Figure S6). Treatment-specific cell type and TC compositions were analyzed, and results are displayed as cell type pie charts (**Figures 4B and** S6), TC overlay plots (**Figures 4C and** S6), TC-specific cell type enrichment heatmaps (**Figures 4D and** S6), and TC-specific cell type composition heatmaps (**Figures 4E and** S6). In this exemplary tumor, immunotherapy-treated tumor center explants showed increased enrichment of CD8^+^ and especially CD4^+^ T cells in TC5 (*“adaptive immune compartment”*) compared to the untreated control (**Figure 4D**, TC5 orange, immunotherapies vs. control). Moreover, we observed an increased enrichment of M1 macrophages, granulocytes and lymphocytes in TC2 (“*myeloid compartment*“) in immunotherapy-treated explants (**Figure 4D**, TC2 light blue, immunotherapies vs. control).

**Figure 4.**
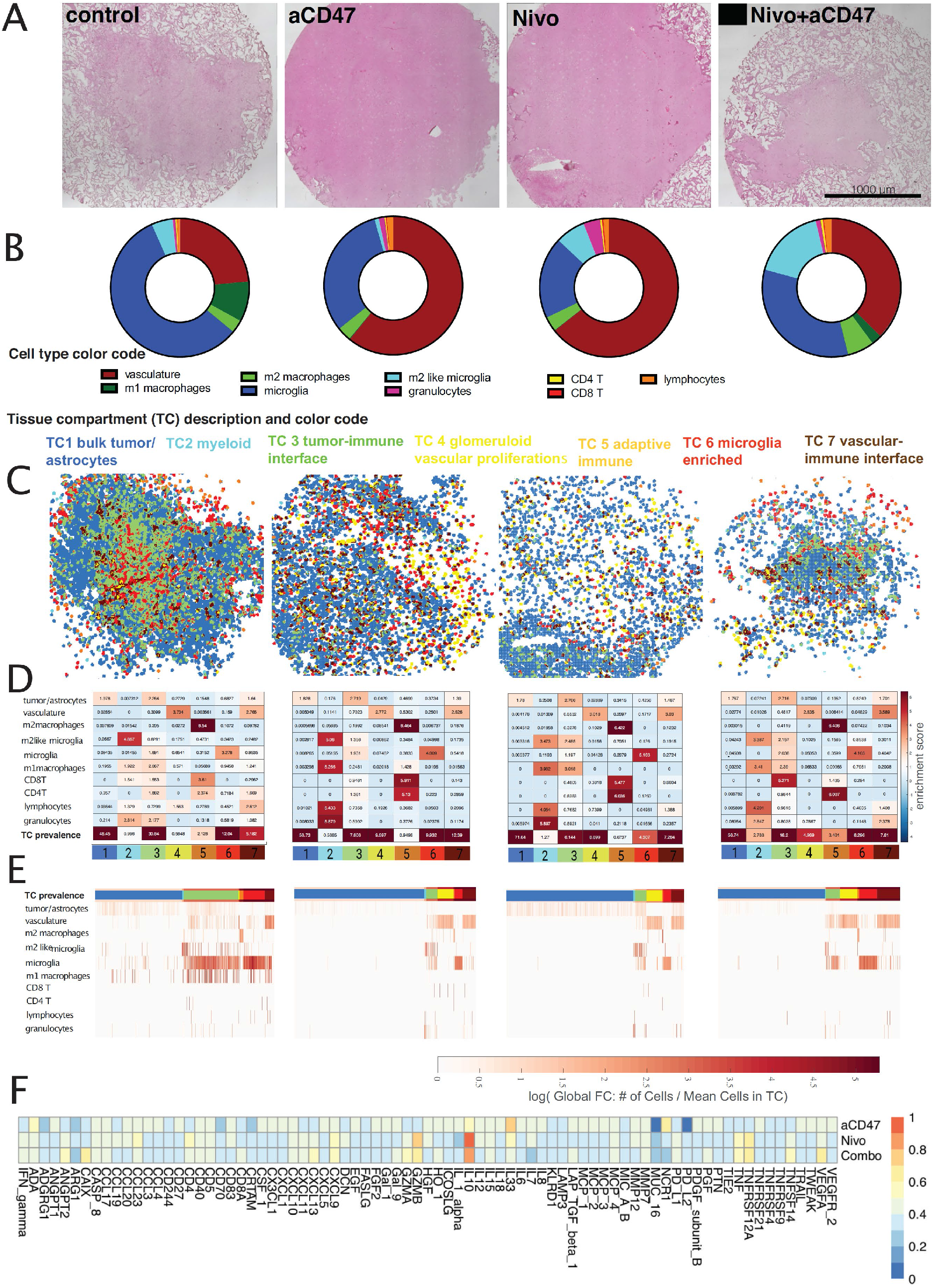
Personalized local immunotherapy assessments in GBM explants by integrated analysis of soluble proteins and spatial iTME rearrangements. Representative example of patient-specific treatment assessment towards ‘local’ immunotherapy (patient 583, center explant). **(A)** H&E stained tissue sections of bioreactor samples after 7 days of immunomodulatory treatment, after CODEX run. **(B)** Cell frequency distribution pie chart per specific explant, excluding tumor cells and astrocytes. **(C)** TC mapping and overlay on tissue sections based on x-y coordinates **(D)** Individual TC enrichment plots depicting fold change of cell type enrichment per TC, and TC prevalence. **(E)** Individual heatmap of the TC composition (percentage of each cell phenotype per TC) after SOM clustering. Individual TCs are denoted by the color bar at the top of the graph; **(F)** individual cytokine profiles resulting from multiplexed proximity extension assays, displayed as fold change in relation to untreated controls. Values are sorted from low to high based on the combinatorial treatment condition (Nivolumab + anti-CD47).

### IFN**γ** levels distinguish immunotherapy responder and non-responder tumor explants

To establish additional criteria for the differentiation between explants responding or not to immunotherapy, we measured 92 soluble proteins including cytokines and chemokines in the media of each bioreactor. For example, in patient 587, we observed increases in several cytokines including interferon-γ(IFNγ) after immunotherapy (**Figures 4F and** S5). Explants from the tumor center displayed higher global soluble protein levels than those from the periphery when integrating all 92 analytes per sample (**Figure 5A**). This was irrespective of the type of immunotherapy applied; however, when comparing soluble protein levels of center vs. periphery in each individual treatment group, no significant differences were identified (**Figure 5B**). Nevertheless, many soluble proteins were present at significantly higher levels in immunotherapy-treated tumor center vs. periphery explants, including IFNγ IL-15 and CXCL-13, amongst others (Figure S9).

**Figure 5.**
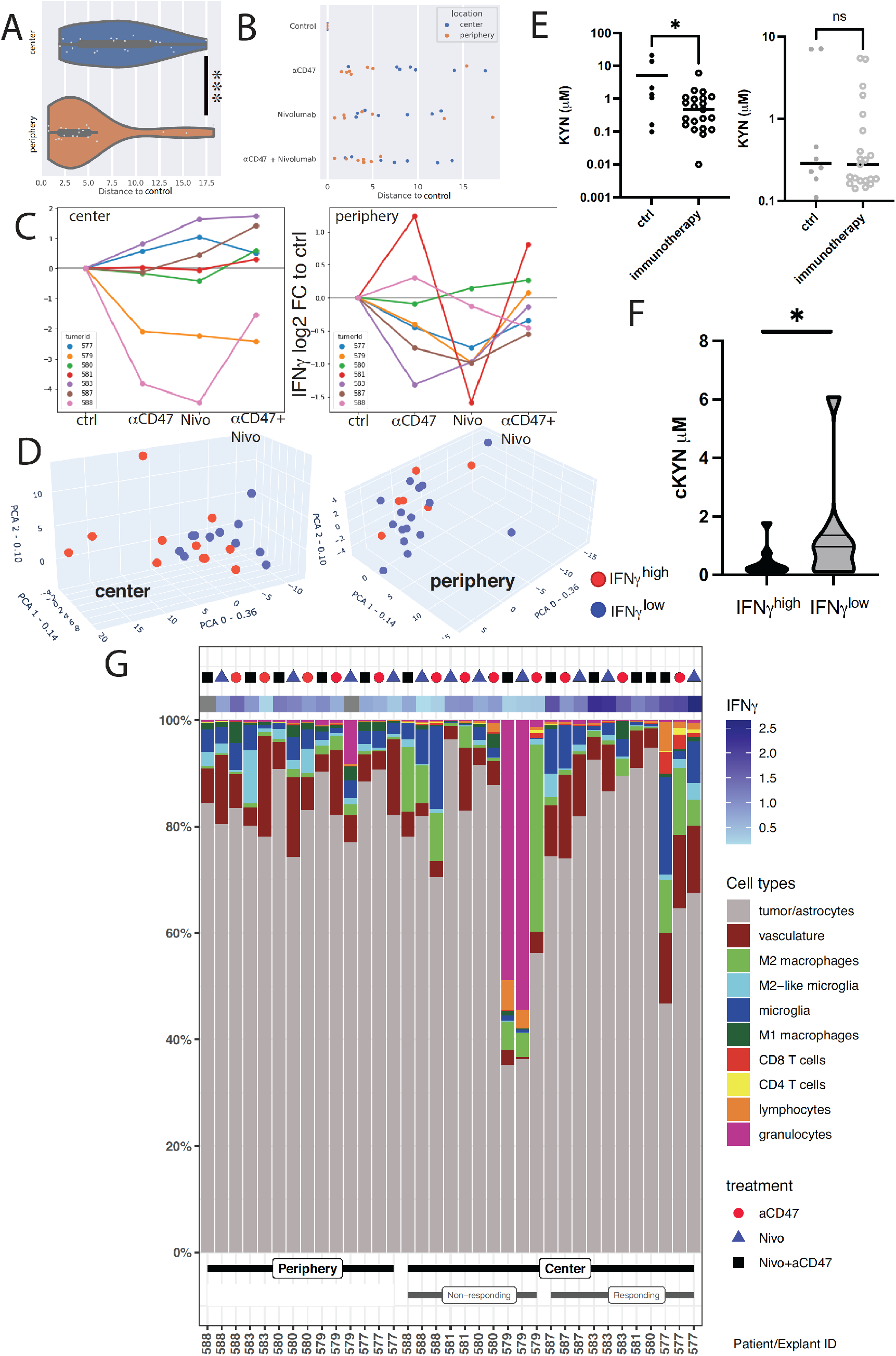
Stratification based on IFNγ levels in culture media reveals a subset of tumor center explants responding to immunotherapy. **(A)** Global soluble protein analysis of bioreactor media in relation to their reference (control samples).Distances to control correspond to euclidean distances between sample and the corresponding control on the first two components of the principal component analysis (PCA) of global cytokine response - resp. 36% and 14% of the total variance. **(B)** Relative distance to control of global soluble protein expression values among immunotherapy treated center and periphery explants. **(C)** Log2 fold change of IFNγ secretion vs. control in center and periphery, per individual patients, conditions, and biopsy location. Samples with a ratio treated/control > 1 were counted as IFN**γ**^high^, independent of the applied treatment regimen. **(D)** 3D principal component analysis (PCA) of global cytokine response in the center (left) and periphery (right) samples, overlayed with information on IFNγ secretion. Red dots: IFN**γ**^high^, blue dots: IFN**γ**^low^. For individual PC weights, see Figure S8. **(E)** Measurement of kynurenine (KYN) levels in bioreactor supernatants after local immunomodulatory treatments depending on biopsy location, left graph representing center biopsies, and the right periphery biopsies. For individual measurements per treatment modality refer to Figure S11. **(F)** Pooled KYN values among immunotherapy treated IFNγ^high^ responder and IFNγ^low^ non-responder explants (student’s t-test). **(G)** Cellular composition (as percentage of total cell count per explant) of individual treated explants stratified according to periphery, IFN**γ**^low^ center, and IFN**γ**^high^ center. IFNγ values represent log2 fold change to untreated control explant. *Statistics:* (A) Mann-Whitney U test; (E) two-tailed Welch’s test, (F) student’s t-test.*p<0.05, **p<0.01, ***p<0.005

To define immunotherapy ‘responder’ and ‘non-responder’ explants, we used IFNγ levels as a surrogate marker of an antitumoral immune response [38]. Compared to untreated controls, IFNγ levels were higher in media from a subset of tumor center explants in all immunotherapy conditions (**Figure 5C**, left panel). In contrast, only few explants from the tumor periphery showed enhanced IFNγ levels upon immunotherapy (**Figure 5C**, right panel). In tumor center explants, combination immunotherapy increased IFNγ levels more than did single treatments, and blocking PD-1 generally induced higher IFNγ levels than did blocking CD47 (**Figure 5C**, left panel). Interestingly, combination immunotherapy of explants from the tumor periphery only increased IFNγ levels in patient 580, whose peripheral samples had a high cell density similar to tumor center biopsies (Figures S4**-5**), and in patient 581 (tissue from tumor periphery not available for analysis).

In addition, to stratify explants and investigate immune response-related protein signatures in an unbiased manner, we performed PCA analysis and displayed explants with high IFNγ levels (IFNγ^high^) as blue symbols and those with low IFNγ levels (IFNγ^low^) as red symbols (**Figure 5D**). In the periphery, no clear segregation pattern was observed, and IFNγ levels were generally uniformly distributed (**Figure 5D**, right panel). In contrast, in the tumor center, IFNγ^high^ responder explants were associated with higher values in the 2 most highly weighted PCs (**Figure 5D**, left panel). PC0 with the highest weight (36%) was characterized by CASP-8, LAP-TGFβ, CSF1, CD40, TNF, MMP7, GAL9 and TNFRSF12A, while PC1 (weight: 14%) was determined by IL-8, followed by myeloid chemokines MCP1, MCP3, CCL4, MCP2, IL-6, and CCL20. In PC2 (weight: 9%), TIE, TRAIL and CD244 were the most important determinants (Figure S8). γ, area under the curve (AUC) receiver-operating characteristics analysis identified other mediators that were potentially associated with an immunotherapy-induced immune response: VEGFA, CXCL10, CXCL9, CCL23, GZMH, IL10, TNFRSF12A and CD4 reached an AUC > 0.78 (**Figure S10A**). Heatmap clustering of IFNγ^high^ and IFNγ ^low^ explants from the tumor center, and the immune-blunted periphery samples visualized the distinct, patient-specific cytokine profiles per treatment (**Figure S10B**).

### Reduced kynurenine levels in media from immunotherapy-treated tumor center explants

As another surrogate marker for immunotherapy response, we measured the metabolite kynurenine (KYN) by LC-MS in bioreactor media after 7 days of culture. KYN is produced from the amino acid L-tryptophan by the enzymes indoleamine 2,3-dioxygenase (IDO) and tryptophan 2,3-dioxygenase (TDO), which have been linked to immunosuppressive iTME states [41]. In untreated control explants, KYN concentrations tended to be higher in the media of tumor center compared to periphery (mean 5.5 vs. 2.1 μM; p=0.08, Figure S11). Immunotherapy-treated center explants had significantly lower KYN levels than untreated M, p=0.01); this effect was not observed in explants from the M, p=0.23) (**Figure 5E**). KYN levels in bioreactor media inversely correlated with IFNγ levels when stratifying the immunotherapy treated explants from tumor centers into IFNγ^high^ responders and IFNγ^low^ non-responders (**Figure 5F**). Individualized assessment showed changes per treatment and location, and in a subset of explants, lower KYN levels after both anti-CD47, anti-PD-1, or combination immunotherapy were observed, even in explants from the periphery (Figure S11). These findings support the notion of an overall stronger activation of antitumoral immunity upon immunotherapy in explants from the tumor center.

### Neoadjuvant GBM therapy and high numbers of intratumoral myeloid cells might portend tumor explant immunotherapy non-response

We compared the cellular composition of IFNγ^high^ responder and IFNγ^low^ non-responder explants in our cohort (**Figure 5G**), and attempted to correlate the explant response to clinical behavior. While no statistically significant differences in cell type composition were seen between responding and non-responding explants, the iTME composition of individual explants could potentially bear important information on the later clinical disease course. Interestingly, the non-responder explants from patient 579 (mesenchymal subtype, methylguanine DNA methyltransferase [MGMT] unmethylated, overall survival 505 days) had very high numbers of granulocytes and M2 macrophages, which might be responsible for an intense immunosuppression in this patient’s explants. The other non-responding explants were from patient 588 (mesenchymal subtype, MGMT methylated), who was initially biopsied and underwent neoadjuvant chemotherapy with temozolomide and radiation therapy before surgical resection. This patient’s tumor center explants contained high numbers of immunosuppressive M2 macrophages, which could explain non-response. Moreover, the overall survival of patient 588 was only 204 days. In patients 580 and 581 (both classical subtype, MGMT methylated), we observed an IFNγ^high^ response in tumor center explants only upon combination immunotherapy.

Interestingly, treated explants from the tumor periphery also displayed IFNγ^high^ responses in those patients, despite their rather low immune cell contents. Importantly, those two patients had better clinical outcomes, with an overall survival of 651 days (patient 580) and >700 days (patient 581), with patient 581 being alive at the time of concluding this study.

### IFN**γ** responder explants show distinct immune cell compositions in their tissue compartments

We next used the information derived from the soluble protein analysis to identify potential immunotherapy-induced changes in iTME architecture. Since an overall treatment response was mainly detectable in explants from the tumor center, we focused our analysis on this group and differentiated between immunotherapy-treated responder and non-responder explants. Differences between these groups were assessed in TC enrichment diagrams to depict TC prevalence and cell type composition per TC (fold change) (**Figure 6A**). In non-responders, no significant changes within the TCs and their prevalence were detected after treatment. In responders, immunotherapy led to a significant lymphocyte diminution in TC2 (“*myeloid compartment”*) and TC7 (*“vascular-immune interface”*) (**Figure 6A**, right lower panel, # in TC2, ## in TC7).

**Figure 6.**
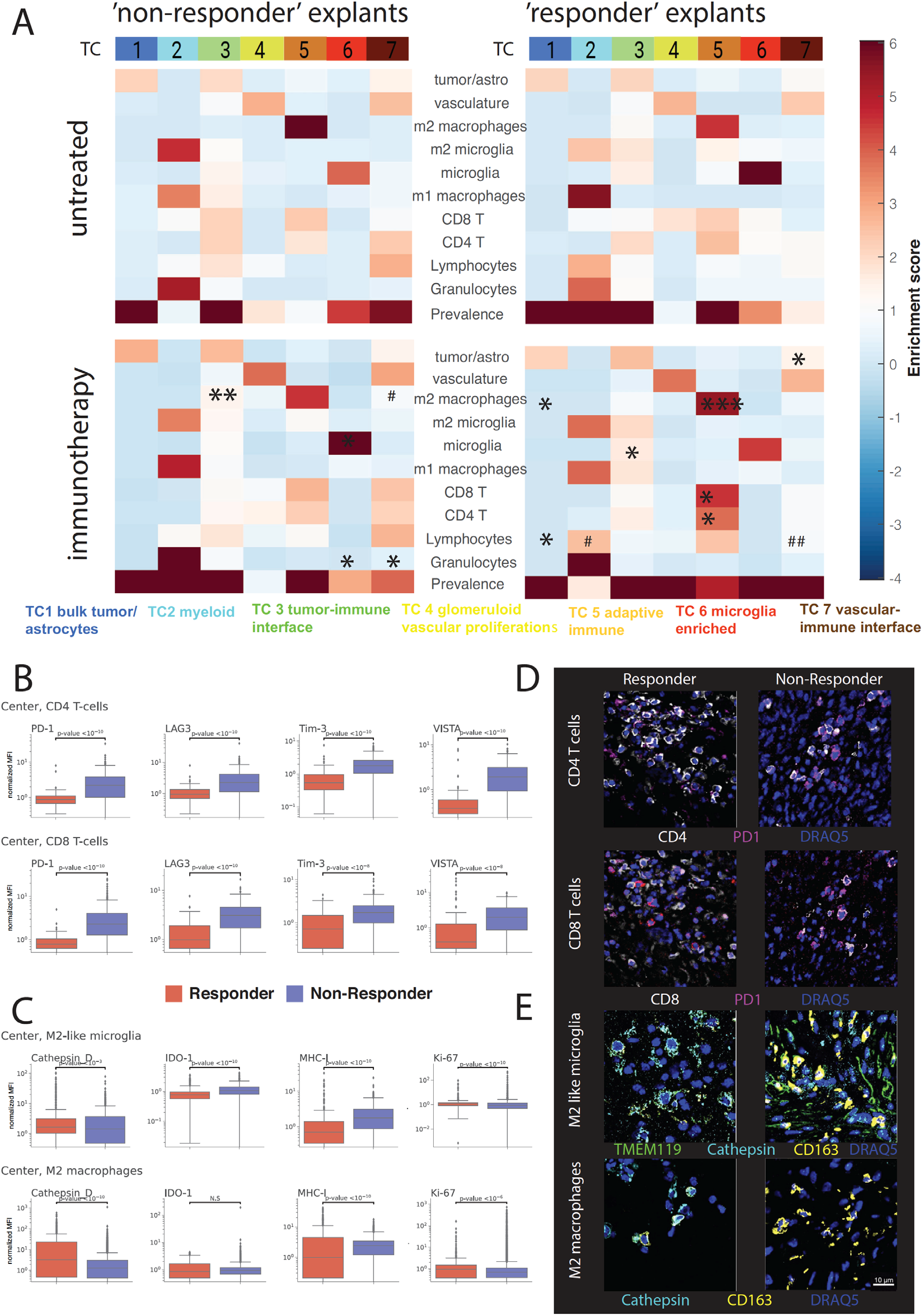
Immunotherapy induces distinct cellular composition shifts within tissue compartments and changes cellular expression levels of functional markers responding explants. **(A)** Enrichment score (ES) of cellular composition in center samples, stratified according to IFNγ secretion (pooled analysis). ***Horizontal analysis in-between treated samples IFN***γ ***vs IFN***γ (Asterisks in condition with higher FC display significant results): TC5: CD8 T cells: mean ES from 1.8 to 4.24; CD4^+^ T cells, mean ES from 1.9 to 4.17; M2 macrophages: mean ES from 4.07 to 5.69; TC6 microglia: mean ES from 5.03 to 4.38. ***Vertical Analysis between untreated and treated samples*** (# in conditions with higher ES display significant results). **(B)** Normalized inhibitory checkpoint molecule expression (mean fluorescent intensity within a specific annotated cell type) on CD4^+^ and CD8^+^ T cells in ***IFN***γ***^low^ vs IFN***γ***^high^*** center explants. **(C)** Normalized expression of functional markers and surrogates of immunosuppression in M2-like microglia and m2 macrophages in ***IFN***γ ***vs IFN***γ center explants. For other cell types and additional markers refer to Figure S12. **(D)** Immunofluorescent representation of PD1 expression on CD4^+^ and CD8^+^ T cells from the original CODEX-stained explants in a responding (Pat. 577, combinatorial condition) and a non-responding (Pat. 579, aCD47 condition) explant. Cells were initially identified based on the cell passports in Figure S3**. (E)** Immunofluorescent representation of Cathepsin D expression on M2-like microglia and M2 macrophages from a responding (Pat. 583, combinatorial condition) and a non-responding (Pat. 588, combinatorial condition) explant. Scale bar: 10 μm. Statistics: (A) * or # p<0.05, ** or ## p<0.01, *** or ### p<0.005, two-tailed Welch t-tests, (B) and (C) Kruskal-Wallis H test with correction for multiple comparisons for median marker expression values.

Moreover, when comparing treatment outcomes between non-responders and responders, the most prominent differences affected TC5 (*“adaptive immune compartment”*), where a significantly higher enrichment of CD4^+^ and CD8^+^ T cells, as well as M2 macrophages was observed in immunotherapy-treated responders (**Figure 6A**, * in lower panels). Conversely, in immunotherapy-treated non-responder explants, M2 macrophages were more enriched in TC3 (*“tumor-immune interface”*). Among treated samples, region prevalences of TC4 (responders 5.8%, non-responders 1%, p=0.001) and TC7 (responders 7.5%, non-responders 3.7%, p=0.01) were increased in the responding samples (Figure S7**, C**). This confirmed not only intra-TC composition shifts, but also overall TC prevalence changes upon immunomodulatory treatments.

### Cell type-specific functional marker expression profiles indicate innate and adaptive immune activation in explants responding to immunotherapy

We then investigated whether individual cell types within explants changed their phenotypic and functional properties upon immunotherapy. In line with our previous observations, diverging activation patterns among cell types per tumor region were noted: CD4^+^ T cells in center explants from responders displayed lower surface levels of the immune checkpoints PD-1, LAG3, TIM-3 and VISTA. In contrast, CD4^+^ T cells from the periphery showed no significant changes in the expression of those immune checkpoints between immunotherapy and control conditions (**Figure 6B, D**, and Figure S12).

Similar observations were made for CD8^+^ T cells (**Figure 6B, D**). Further, CD8^+^ T cells expressed lower levels of IDO-1, which is consistent with our findings of lower KYN levels in this group (Figures S10 and S12). The “lymphocyte” cluster showed an alternate pattern in terms of inhibitory immune checkpoint expression with partial upregulation of PD-1, LAG-3 and TIM-3 in both center and periphery. This subset was driven to proliferate in the responder center explants as assessed by enhanced Ki67 staining intensity (Figure S12, lymphocytes).

M1 macrophages displayed a double-sided response among responders and non-responders. Overall, a stronger proliferation (Ki67) was observed in the responding center explants, albeit accompanied by higher levels of IDO-1. The immune checkpoints PD-1, LAG-3, TIM-3 and VISTA were upregulated in responders, conversely, the lysosomal enzyme Cathepsin D, suggestive for a higher phagocytic activity, was also elevated [3]. We did not observe marked differences between center and periphery (Figure S12, M1 macrophages).

Interestingly, changes in M2 macrophages, which were the most abundant myeloid immune phenotype in the cohort and are an important cell population to address in GBM immunotherapy, were more pronounced in responders: Ki67 and Cathepsin D were significantly enhanced in IFNγ samples. MHC-I, a potential ‘dont eat me signal’ [2], was downregulated in responding center explants. Also, IDO-1 was downregulated in the responder periphery upon treatment. M2 macrophages might be the most important source of IDO-1 activity in the tumors under investigation, and the suppression of IDO-1 in this cell type could be a surrogate for an effective intratumoral immune response. In terms of other inhibitory checkpoints, a downregulation of PD-1, TIM-3 and LAG-3, and in contrast upregulation of the macrophage checkpoint VISTA was observed (**Figure 6C, E,** Figure S12, M2 macrophages), which might equally have implications on M2 macrophage tumoricidal activity [15].

Finally, we assessed marker emergence in microglia and M2-like microglia. In line with findings in M2 macrophages, the phagocytic marker Cathepsin D was elevated in responding samples, whereas expression of IDO was lower in this group, pointing towards reduced immunosuppressive features. Moreover, responsive microglia tended to proliferate more in both center and periphery, independent of their polarization state. In all 3 examined treatments, we found a profound downregulation of MHC-I. Concerning classical immune checkpoint expression, microglia and M2-like microglia tended to downregulate PD1, but the potential compensatory immune checkpoint VISTA was increased in responding samples (**Figure 6C, E** Figure S12, microglia and M2-like microglia).

Although we were focusing on the iTME in this study, we also noted specific marker intensity changes in tumor cells/astrocytes as a consequence of iTME remodeling. Most importantly, surrogates of an antitumoral response included downregulation of CD47 and PD-L1 in both center and periphery responder explants, lower levels of tumor-derived IDO-1, and a less pronounced cellular proliferation by Ki-67 staining intensity (Figure S12).

Taken together, this study provides an in-depth proof of concept analysis of a potential immunotherapy response in patient-derived GBM explants with divergence between tumor center and periphery.

## Discussion

The iTME in GBM is a complex, highly immunosuppressive ecosystem. To date, immunotherapies against GBM have mostly failed. However, data in systemic, neoadjuvant application of T cell checkpoint inhibitors point towards a tumor intrinsic immune response [28]. Here, we modeled the iTME of *IDH* wild type GBM and its potential amenability to local immunomodulatory treatments in a patient-individualized way. For this, we combined multiple technologies including neuronavigated intraoperative tissue acquisition, live, intact 3D tissue cultures, GBM-relevant immunotherapy, FFPE-CODEX, soluble protein analysis, and metabolomics.

Our integrative framework enabled us to describe components of the GBM iTME in depth, and to monitor immune-mediated responses on a combined spatial and biomarker-based basis. We identified cell composition differences between center and periphery, described seven conserved TCs of the GBM iTME across explants, and probed their organization, functional states, and communication.

In a subset of tumor center explants, we identified solid surrogates of an intratumoral immune response by TC shifts, intra- and inter-TC composition changes, distinct functional and phenotypic changes of the annotated cell types under question, and a favorable reversion of immunosuppressive features. To our surprise, a strong adaptive immune response was almost exclusively ensuing in the center explants, whereas the peripheral invasion zone acted as an “immune desert” zone and displayed much lower responses in both cellular rearrangements and soluble factor levels. This observation might be of translational clinical relevance and might support the concept of local neoadjuvant immunomodulatory treatments before bulk tumor resection or after re-emergence of contrast-enhancing tumor areas.

However, the periphery was not completely inert to immunotherapies. At least in responding tumors, the dominating M2-like microglia and M2 macrophages substantially changed their marker profile towards antitumoral action. Although the study was iTME centered, we also observed perturbations of the tumor cells/astrocytes in responding explants. Importantly, responding explant tumor cells/astrocytes had lower levels of CD47, PD-L1, IDO-1, and CD73. These expression level differences on the tumor cells might impede a successful anti-tumoral immune response, but could also have been induced by a successful reversion of the iTME by the treatments under investigation. Clearly, responding explants displayed lower tumor cell proliferation, which serves as a direct surrogate of anti-tumor efficacy.

Recent discoveries in *ex vivo* culture systems have equally unraveled the potential of tumor resident T-cells to contribute to an intratumoral immune response upon T-cell checkpoint inhibition, albeit in extracerebral tumors [38]. However, patient-derived data on innate immune modulators in a prototypic, myeloid-dominated, immunologically ‘cold’ tumor such as GBM are scarce.

Efforts to model GBM *ex vivo* in a patient-centered fashion are needed to find potential treatment combinations that are directly translatable to patient care. Few reports on other 3D GBM *ex vivo* culture systems have been described: for example, GBM organoids serve as a valid tool to study personalized therapies, and can even be co-cultured with chimeric antigen receptor T cells [18]. However, organoids lack the integration of microenvironment constituents and are therefore more suited to study direct tumor targeting drugs. Further, 3D bioprinted and vascularized models have been established that artificially add components of the GBM iTME such as microglia to the bioprinted system [22]. Other authors utilized spheroids, organotypic slice cultures or tumoroids to study the tumor complexity as reviewed by Souberan et al. [33]. However, most of the current models lack an integrative GBM iTME, which has important implications for immunotherapy response assessment. The perfused bioreactor approach partly overcomes this limitation with the caveat that the contribution influx of immune cells from the peripheral circulation was not assessed in our proof-of-concept study.

Our proof-of-concept study has several limitations. Although we were analyzing more than 800’000 single cells in-depth, the patient sample size of the study was small, limiting the interpretation of systematic correlations with clinical outcomes, survival or transcriptomic GBM subtype or methylation data. However, we identified two non-responding patients that were either pre-treated by standard of care therapy (patient 588) or had a massive immunosuppressive granulocyte and M2 macrophage infiltration (patient 579). These explants were not able to raise a sufficient IFNγ-associated response, and had no spatial rearrangement of adaptive immune cells within their iTME despite their high cellularity and presence of adaptive immune cells. Further, two patients (Pat. 580 and 581) displayed an intermediate or ‘partial’ response based on the cytokine exploration, and combinatorial immunotherapies with both anti-CD47 and anti-PD1 antibodies raised an IFNγ response, which hinted towards a synergistic effect of targeting both innate and adaptive immune compartments. Interestingly, these two patients had the longest overall survival among this cohort, but this might also be related to other more favorable prognostic factors including the patients’ relatively young age and positive MGMT status.

Furthermore, intratumoral heterogeneity even within patient-matched experimental series posed a problem to the interpretation of the results, necessitating more biopsies per treatment condition. The small sample size forced us to pool different immunotherapy modalities (e.g., T cell checkpoint inhibition vs. myeloid checkpoint inhibition) in the TC composition analysis, and precluded us from identifying cellular reactions to either modality alone.

The granularity of our clustering strategy, especially concerning T cell subsets, was lower than expected in our study. We were not able to identify NK T cells, regulatory T cells or B cells with certainty, and summarized these cells, if they were present, into a ‘lymphocyte’ cluster that precluded us from a more detailed T or B cell analysis. However, we are confident that the normalized functional marker expression analysis in the T cells within explants strictly speaks in favor of phenotype switches in the cell types under question. Hence, GBM is dominated by myeloid cells, and we provide an in-depth phenotypic characterization of microglia and macrophage subsets within GBM. After treatment, compensatory checkpoints were upregulated, e.g., in the case of the checkpoint protein VISTA (V-type immunoglobulin domain-containing suppressor of T-cell activation) in both center and periphery M2 macrophages and microglia responding explants. Whether this observation represents an important immune escape or resistance mechanism is subject to further investigation, but underscores the plasticity of the innate iTME upon therapeutic interventions. Our study focused on immune cells and included only few tumor cell markers which impeded us from discriminating tumor cells from reactive astrocytes, and from assigning different oncogenic driver mutations to tumor cell subclones. Nevertheless, we were able to assess potential predictive markers such as CD47 or PD-L1 quantitatively on tumor cells/astrocytes with implications on immunotherapy response.

Overall, the proposed strategy of combining complex 3D tissue culturing with multiplexed imaging and targeted secretome analysis is not yet standardized and far from reaching implications for clinical practice. Here, demonstrated the feasibility of this concept, and future developments may implicate to concentrate patient stratification efforts on miniaturization of bioreactor systems, more center biopsies per patient, fewer informative functional iTME markers and IFNγ-associated cytokines with predictive capacity. This approach would enable us to test a variety of immunotherapy combinations and tumor targeting drugs on a larger patient cohort and lead to key data that could be correlated with clinical outcomes.

## Acknowledgments

We thank Prof. Alfred Zippelius (Department of Oncology, University Hospital Basel, Switzerland), Prof. Doron Merkler (Department of Pathology, University Hospital Geneva, Switzerland), Prof. Jens Schittenhelm (Department of Pathology and Neuropathology, University Hospital Tübingen, Germany) and Prof. Sheila Singh (Neurosurgery Department, University of Hamilton, Ontario, Canada) for critically reading the manuscript. The anti-PD-1 and anti-CD47 antibodies for bioreactor treatment were a generous gift from Prof. Heinz Läubli (Department of Oncology, University Hospital Basel, Switzerland). We are grateful to Prof. Stephan Frank and Dr. Jürgen Hench (Department of Pathology, University Hospital Basel, Switzerland) for histopathological and molecular workup of the GBM specimens. This work was supported by a Swiss National Science Foundation Professorial Fellowship (PP00P3_176974); the ProPatient Forschungsstiftung, University Hospital Basel (Annemarie Karrasch Award 2019); the Department of Surgery, University Hospital Basel, to G.H.; and by The Brain Tumour Charity Foundation, London, UK (GN-000562) to G.H. and C.M.S. Research in the G.P.N. laboratory was supported by the US NIH (5U54CA20997103, IDIQ17X149); the US DOD (W81XWH-14-1-0180); the US FDA (DSTL/AGR/00980/01); Cancer Research UK (C27165/A29073); the Bill and Melinda Gates Foundation (OPP1113682); the Parker Institute for Cancer Immunotherapy; the Beckman Center for Molecular and Genetic Medicine; and the Rachford & Carlotta A. Harris Endowed Chair. C.M.S. was supported by an Advanced Postdoc Mobility Fellowship from the Swiss National Science Foundation (P300PB_171189, P400PM_183915), and an International Award for Research in Leukemia from the Lady Tata Memorial Trust, London, UK. T.S. was supported by an UICC Technical Fellowship (TF/18/625070). D.P. was supported by an NIH T32 Fellowship (AR007422), an NIH F32 Fellowship (CA233203), a Stanford Dean’s Postdoctoral Fellowship, and Stanford’s Dermatology Department. S.S.B. was supported by aStanford Bio-X Interdisciplinary Graduate Fellowship and Stanford’s Bioengineering Department. G.L.B was supported by an NIH T32 Fellowship (5T32AI007290-34).

## Author contributions

Conceptualization: G.H. and C.M.S. Methodology: T.S., C.P.Z., E.M.B., W.D., A.T.W., T.A.M., S.Z., F.L.G., M.G.M., M-F.R., C.M.S. and G.H. Clinical data and patient integration: P.S. and G.H. Software: C.P.Z., E.M.B., W.D., S.S.B., G.L.B. Validation: T.S., C.P.Z., E.M.B., W.D., J.F., C.P., D.P., C.M.S. and G.H. Formal Analysis: T.S., C.P.Z., E.M.B., W.D., S.H., A.T.W., M.M.E., C.M.S. and G.H. Investigation: T.S., C.P.Z., E.M.B. and G.H. Resources: T.S., M.G.M., C.M.S. and G.H. Writing - Original Draft: G.H. and C.M.S. Writing - Review and Editing: all authors Supervision: G.H. and C.M.S. Project Administration: G.H. and C.M.S. Funding acquisition: T.S., G.P.N., C.M.S. and G.H.

## Data and material availability

All data needed to evaluate the conclusions in the paper are present in the paper and/or the Supplementary Materials. Single cell data table of clustered, annotated cell types with metadata can be downloaded from Mendeley (https://data.mendeley.com/datasets/f9hfcfyt93/1). Raw primary imaging data can be obtained from the authors directly upon reasonable request.

## Supplementary Figure Legends

**Figure S1. Biopsy Locations.** Biopsies were acquired using the intraoperative navigation system Brainlab Automatic Image Registration (Brainlab). Preoperative planning of surgeries and labeling of important eloquent structures was performed in all cases, based on diffusion tensor imaging (DTI) or anatomical considerations. Intraoperative screen captures of both contrast enhancing center and peripheral invasion zone were acquired to document accuracy of the biopsy location. Intraoperative 5-ALA fluorescence was used as an additional stratifier for center-periphery discrimination (5-ALA fluorescence high for vital, contrast enhancing tumor vs. 5-ALA fluorescence low for infiltration zone).

**Figure S2. Bioreactor setup and validation. (A)** Workflow of assembling bioreactor 3D perfusion culture directly after tumor resection. **(B)** *Left:* Representative H&E stained microphotographs of individual tumor samples (center biopsy) at day 0, and after 7, 14 and 21 days of either perfusion culture or static culture condition (40x magnification). *Right:* Analysis of nuclear counts per 10 high power fields in 3 independent samples in relation to the D0 sample in perfused and static samples. **(C)** *Left:* Representative anti-Ki-67 immunohistochemical stained slides comparing static and perfused culture conditions at day 0, and 14 and days 21 after ex vivo culture. Insert: proliferating tumor cells invading the bioreactor scaffold after 21 days of *ex vivo* culture (40x magnification). *Right:* Relative quantification of Ki-67^+^ proliferative cells after 14 and 21 days of *ex vivo* culture in static and perfused conditions, compared to freshly obtained GBM tissue (D0). **(D)** *Left:* Representative anti-CD11b immunohistochemical stained slides comparing static and perfused culture conditions at day 0, and 14 and days 21 after *ex vivo* culture. Insert: myeloid cells invading the bioreactor collagen scaffold after 21 days of *ex vivo* culture (40x magnification). *Right:* Relative quantification of CD11b^+^ myeloid cells after 14 and 21 days of ex vivo culture in static and perfused conditions, compared to freshly obtained GBM tissue (D0). Data are obtained from 3 individual explants in each static and perfusion culture, and represented as mean +/- SEM. *Statistics:* *p<0.05, **p<0.01, ***p<0.005, student’s t-tests.

**Figure S3. Cell Passports.** *<u>Page 1-4:</u>* After clustering, single cells from the 173 resulting initial clusters were overlaid on the raw data fluorescent images and on H&E stains of TMAs based on X/Y positions and visually verified based on marker expression profiles, morphology and localization within the tissue. Similar clusters were manually merged, resulting in 10 final clusters. Here, representative clusters are shown as green crosses based on X-Y coordinates of the cells contained in that cluster, overlaid on the montages of different samples. For each of these clusters, examples of markers important for the cluster identification (positive and negative) and DRAQ5 nuclear stain are shown. *<u>Page 5</u>:* Furthermore, flow-cytometry-like plots gated on DRAQ5/DAPI double-positive single cells were generated to confirm the presence of phenotype defining markers on the cell population of interest. As an example, CD4 T (CD4^+^, CD3^+^), CD8 T (CD3^+^, CD8^+^), microglia (CD45^+^, TMEM119^+^, CD68^+^, CD11^+^, CD163^-^), M2-like microglia (CD45^+^, TMEM119^+^, CD68^+^, CD11b^+^, CD163^+^), M1 macrophages (CD45^+^, CD11b^+^, CD68^+^), M2 macrophages (CD45^+^, CD11b^+^, CD68^+^, CD163^+^), granulocytes (CD45^+^, CD11b^+^, CD15^+^, S100A9^+^), tumor cells (GFAP^+^, CD45^-^, CD31^-^) and vasculature (CD45^-^, CD31^+^, CD34^+^) in patient 577, center explant, are depicted.

**Figure S4: Cell density calculations across center and periphery explants** *Page 1:* exemplary center explant. Top: H&E overview (left), scatter plot of cells with size representation (right). Middle: KDE (Kernel Density Estimation) of cells according to location based on x-y coordinates (left), and in consideration of cell size (right). Bottom: density distribution histogram and cell number. *Page 2:* exemplary periphery explant: Top: H&E overview (left), scatter plot of cells with size representation (right). Middle: KDE (Kernel Density Estimation) of cells according to location based on x-y coordinates (left), and in consideration of cell size (right).. Bw = bandwith, which determines how smooth the KDE becomes. Bottom: density distribution histogram and cell number. *Page 3:* smoothened cell density histograms across center and periphery explants. PDF=probability density function (y-axis), approximate 2D number density (x-axis).

**Figure S5. Frequencies of cellular phenotypes (center vs. periphery) in explants. Proportions of cellular phenotypes per biopsy location in explants: (A)** The proportions of annotated cells per explant are plotted against biopsy location (center vs. periphery). Each dot in the bar graphs corresponds to one bioreactor sample. N=30 center samples and n=23 periphery samples. **(B)** zoomed-in representation of CD4 T cells and lymphocytes. Statistics: Wilcoxon tests.

**Figure S6. Individualized treatment outcomes per patient and per region, individualized cytokine measurements per patient and per region.** For each patient (center and periphery), a set of personalized calculations were generated. *1st row*: H&E (stitched together after CODEX run) stains of explants control (GBM) and treatment conditions, scale bar: 1000μm. *2nd row:* point density plot (x-y plot is divided into a 100×100 grid, in which every rectangle is colored corresponding to the number of elements within this square). *3rd row:* Cell overlay, each dot represents an individual, annotated cell type according to the final cell type annotation. Color code of cell types displayed on the right; 4th row: pie chart of cell type proportions within each individual explant. *5th row*: tissue compartment (TC) overlay, according to 7 TCs outlined in Figure 3 (cf. Figure 3 and text for TC naming and numbering). *6th row:* TC region heatmap depicting the TC frequency (top bar, similar color code as in TC overlay), and the global fold change (number of cells per mean cells in TC) of specific cell types per TC. *7th row*: region enrichment heatmaps depicting the fold change enrichment of cell type composition per TC, and the overall percentage of the TC within an individual sample.

**Figure S7. TC enrichment scores per therapeutic condition and location; TC prevalence changes in respondig vs. non responding explants.** Individual enrichment scores of cell types in respective TCs stratified per treatment and biopsy locations in pooled explants-(**A**) center explants: n=6 for control, n=7 for anti-CD47, n=7 for Nivolumab and n=7 for anti-CD47+Nivolumab; (**B**) periphery explants: n=5 for control, n=5 for anti-CD47, n=4 for Nivolumab and n=5 for anti-CD47+Nivolumab. (**C**) TC prevalence changes in non-responding vs. responding center explants, cf. Fig. 6A, lower panels. Statistics: *p<0.05, **p<0.01, Welch’s t-test,

**Figure S8. PCA Loadings of unbiased cytokine analysis in center and periphery.** PCA loadings of the principal component analysis in Figure 5D.

**Figure S9. Differential expression of cytokines in immunotherapy-treated center and periphery explants.** Results from multiplexed bead based proximity extension assay of 92 analytes in the bioreactor culture supernatant. Comparison of pooled treated (all conditions) center (n=21) and periphery (n=21) explants independent of response. Only significant results are displayed. Values are displayed as mean +/- SEM. NPX: Log2 scaled **normalized protein expression**. Statistics: ANOVA with multiple comparison adjustments. Welch’s t-test between the groups.

**Figure S10. AUC-ROC analysis of cytokines associated with IFNγ, heatmap clustering of cytokines stratified per IFNγ status and location. (A)** Based on the IFNγ stratification criterion, we performed an AUC-ROC analysis of other associated cytokines in center explants. Cytokines with a AUC > 0.78 were determined to have a high association with IFNγ and are listed. **(B)** Normalized heatmap depicting Z-scores of the the overall cytokine response per treatment in IFNγ^high^ and IFNγ^low^ center explants, and periphery explants. Samples were normalized to internal control condition.

**Figure S11. KYN modulation in individual patient explants after immunotherapy, and based on IFNγ status.** Assessment of KYN concentrations (nM) in explant culture supernatant by LC-MS in control and immunotherapy treated center and periphery explants. Each dot represents an individual LC-MS measurement. The coloured connecting lines are pairs of individual patients. *Statistics:* *p<0.05, **p<0.01, ***p<0.005, Wilcoxon test.

**Figure S12. Assessment of immune checkpoint, proliferation and activation markers on cellular subsets, stratified per IFNγ and IFNγ responses, and biopsy location.** Patient samples were stratified according to IFNγ response after immunomodulatory treatment (see Figure 5C and G), and pooled. Log of normalized mean fluorescent intensity (MFI) values of the respective markers in individual cellular phenotypes per region (center vs. periphery) and per IFNγ response (low vs. high) were plotted. *Page 1*: center CD4 T cells, *page 2*: periphery CD4 T cells; *page 3*: center CD8 T cells, *page 4:* periphery CD8 T cells; *page 5*: center lymphocytes; page 6: periphery lymphocytes; *page 7*: center M1 macrophages, *page 8*: periphery M1 macrophages; *page 9*: center M2 like microglia, *page 10*: periphery M2 like microglia; *page 11*: center M2 macrophages, *page 12*: periphery M2 macrophages; *page 13*: center microglia, page 14: periphery microglia; *page 15*: center tumor cells/astrocytes; page 16: periphery tumor cells/astrocytes. Statistics: non-parametric Kruskal-Wallis H-test [21], using Benjamini-Hochberg procedure to control the false discovery rate.

## Supplementary Tables

**Table S1:** Clinical data and pathological-molecular tumor characteristics.

**Table S2:** Purified antibodies used in the CODEX analysis (first sheet), CODEX multicycle scheme, exposure times and antibody dilutions (second sheet), list of oligonucleotides used (third sheet).

