## Supplementary material for "Immunotherapy of glioblastoma explants induces interferon-γ responses and spatial immune cell rearrangements in tumor center, but not periphery": Suppl Figures S1-12

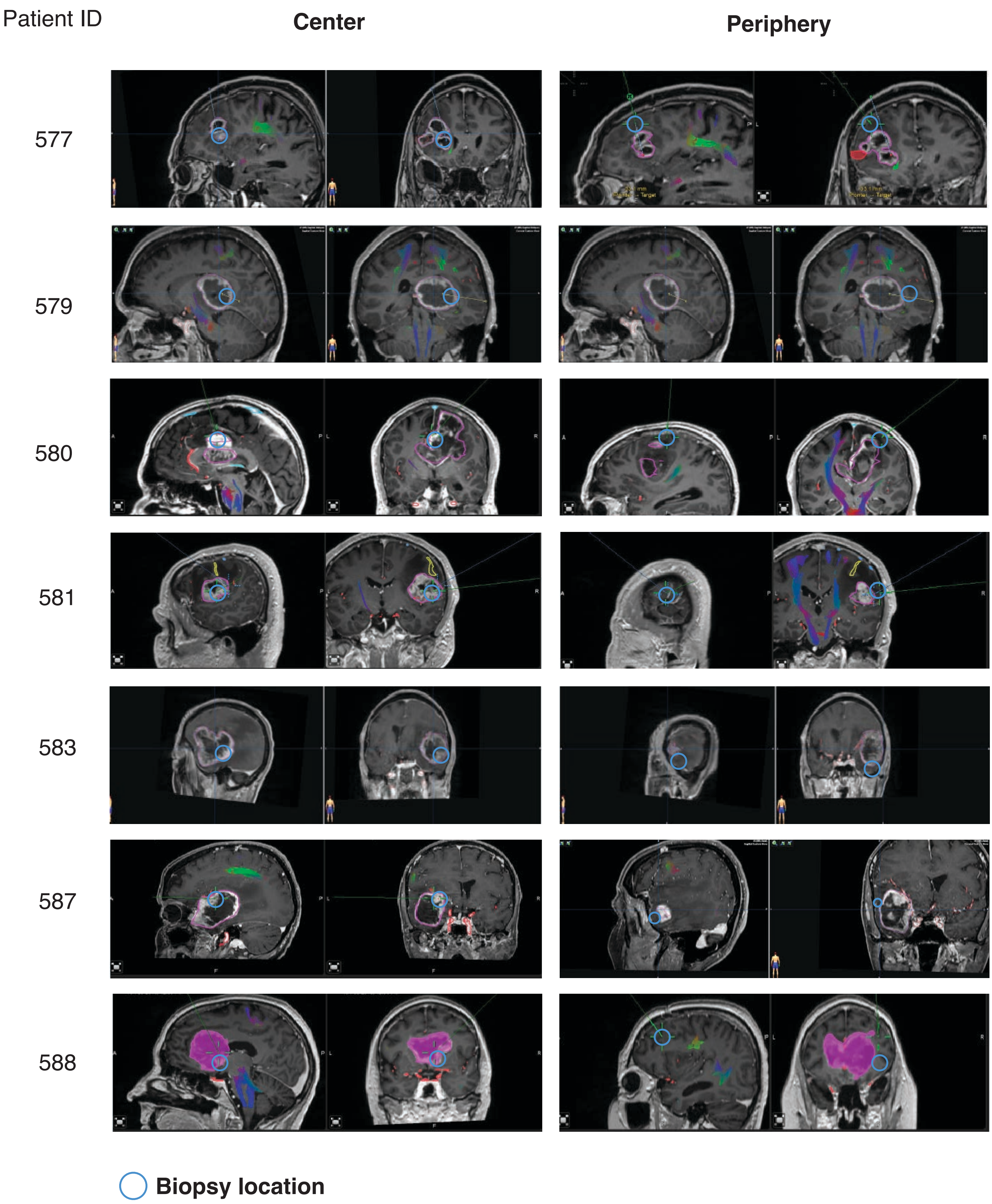

Patient ID
Center
Periphery
577
579
580
581
583
587
588
Biopsy location


### Figure S1

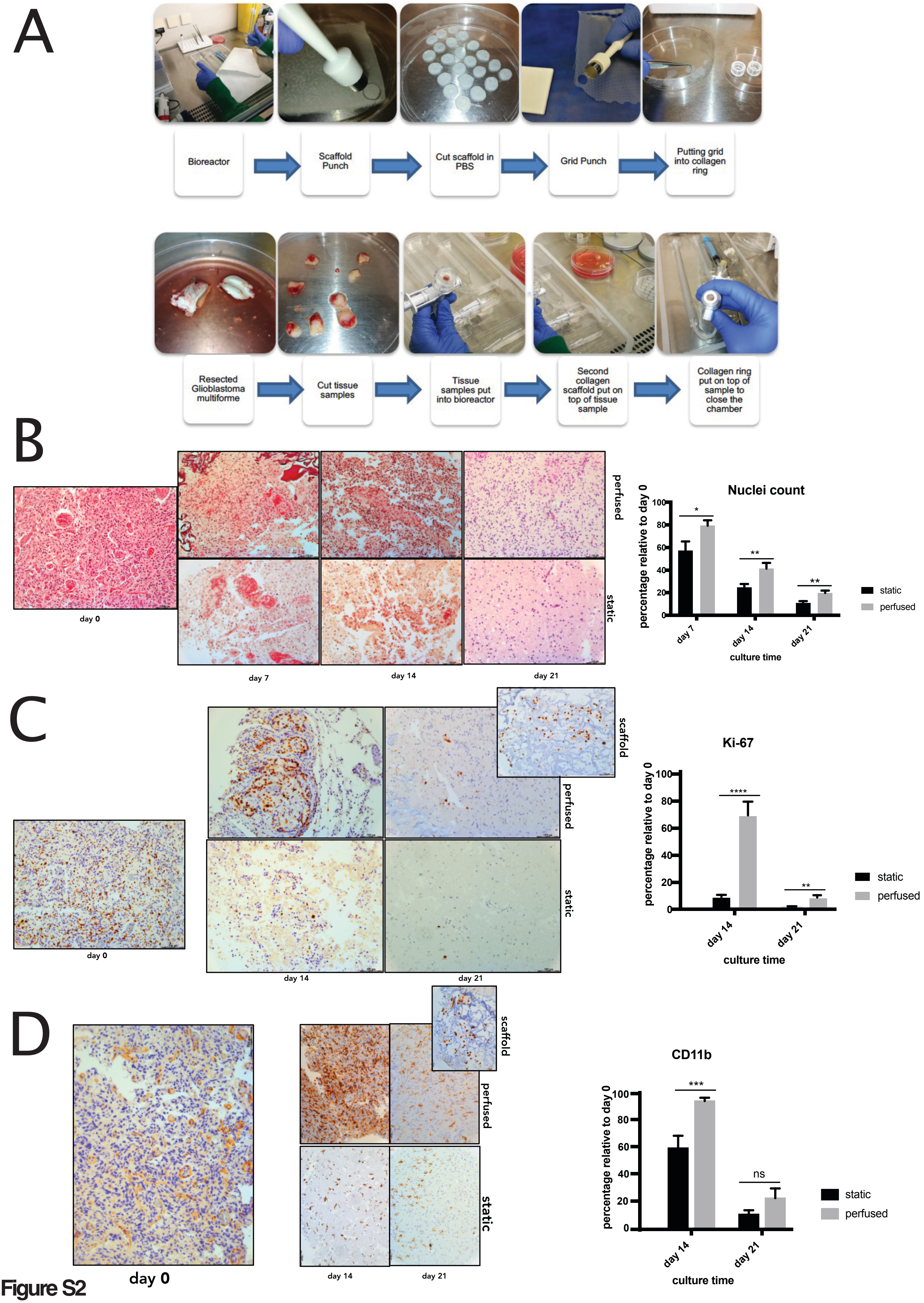
A
Bioreactor
Scaffold Punch
Cut scaffold in PBS
Grid Punch
Putting grid into collagen ring
Resected Glioblastoma multiforme
Cut tissue samples
Tissue samples put into bioreactor
Second collagen scaffold put on top of tissue sample
Collagen ring put on top of sample to close the chamber
B
perfused
static
day 0
day 7
day 14
day 21
Nuclei count
percentage relative to day 0
culture time
static
perfused
C
scaffold
perfused
static
day 0
day 14
day 21
Ki-67
percentage relative to day 0
culture time
static
perfused
D
scaffold
perfused
static
day 0
day 14
day 21
CD11b
percentage relative to day 0
culture time
static
perfused
ns

Figure S2

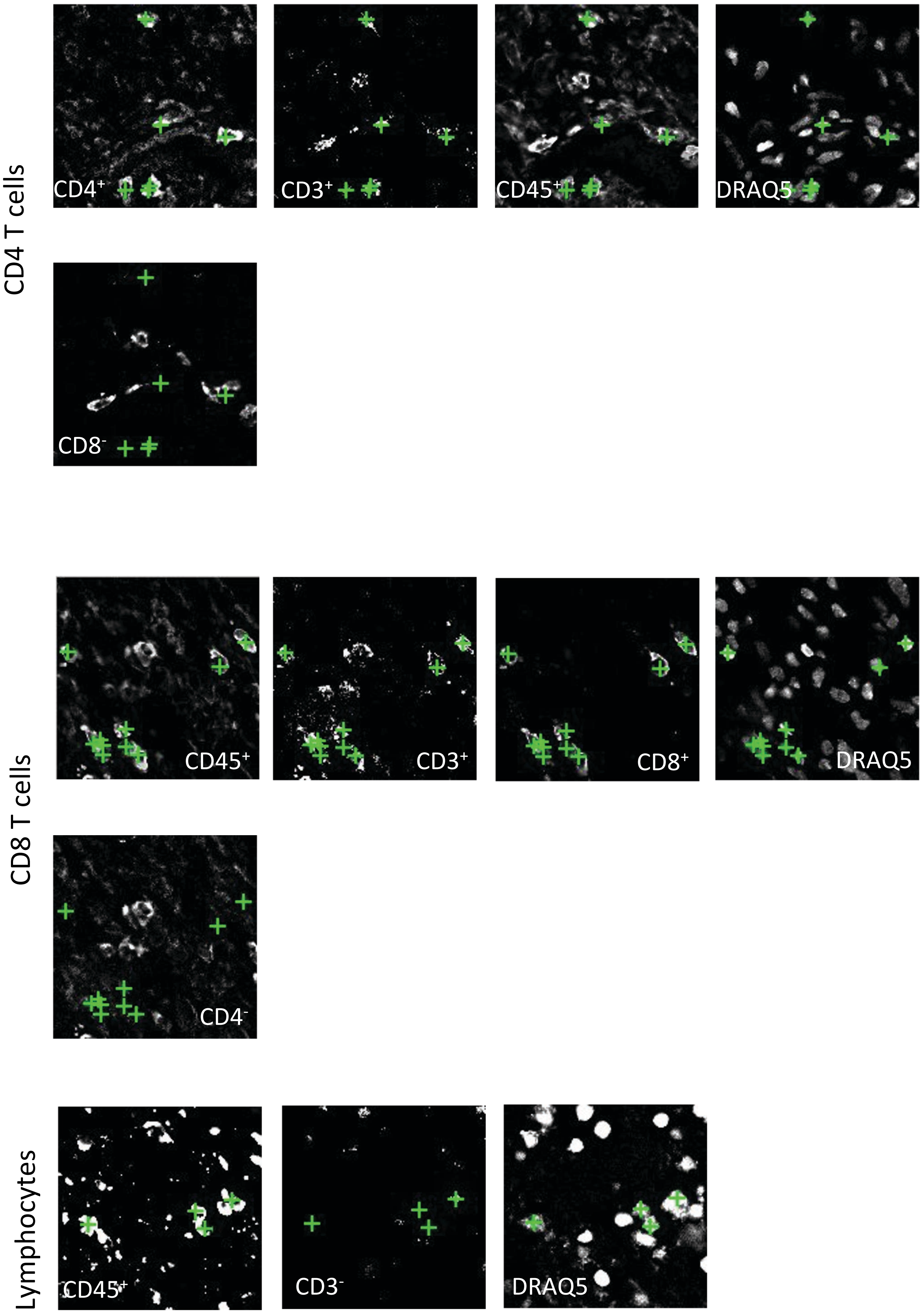
CD4 T cells
CD4+
CD3+
CD45+
DRAQ5
CD8-
CD8 T cells
CD45+
CD3+
CD8+
DRAQ5
CD4-
Lymphocytes
CD45+
CD3-
DRAQ5



**Figure S3**

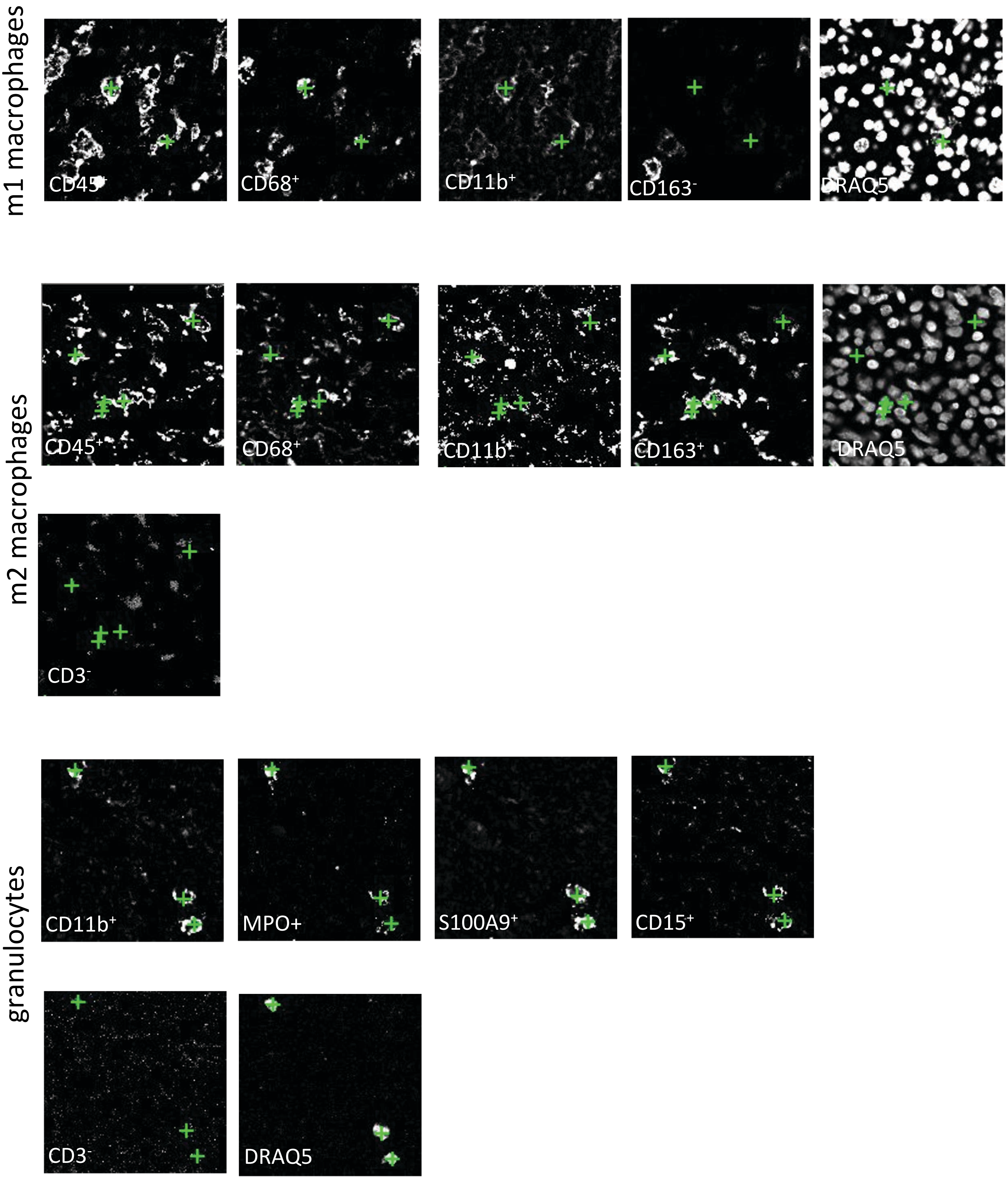
m1 macrophages
CD45+
CD68+
CD11b+
CD163-
DRAQ5
m2 macrophages
CD45+
CD68+
CD11b+
CD163+
DRAQ5
CD3-
granulocytes
CD11b+
MPO+
S100A9+
CD15+
CD3-
DRAQ5

**Figure S3**

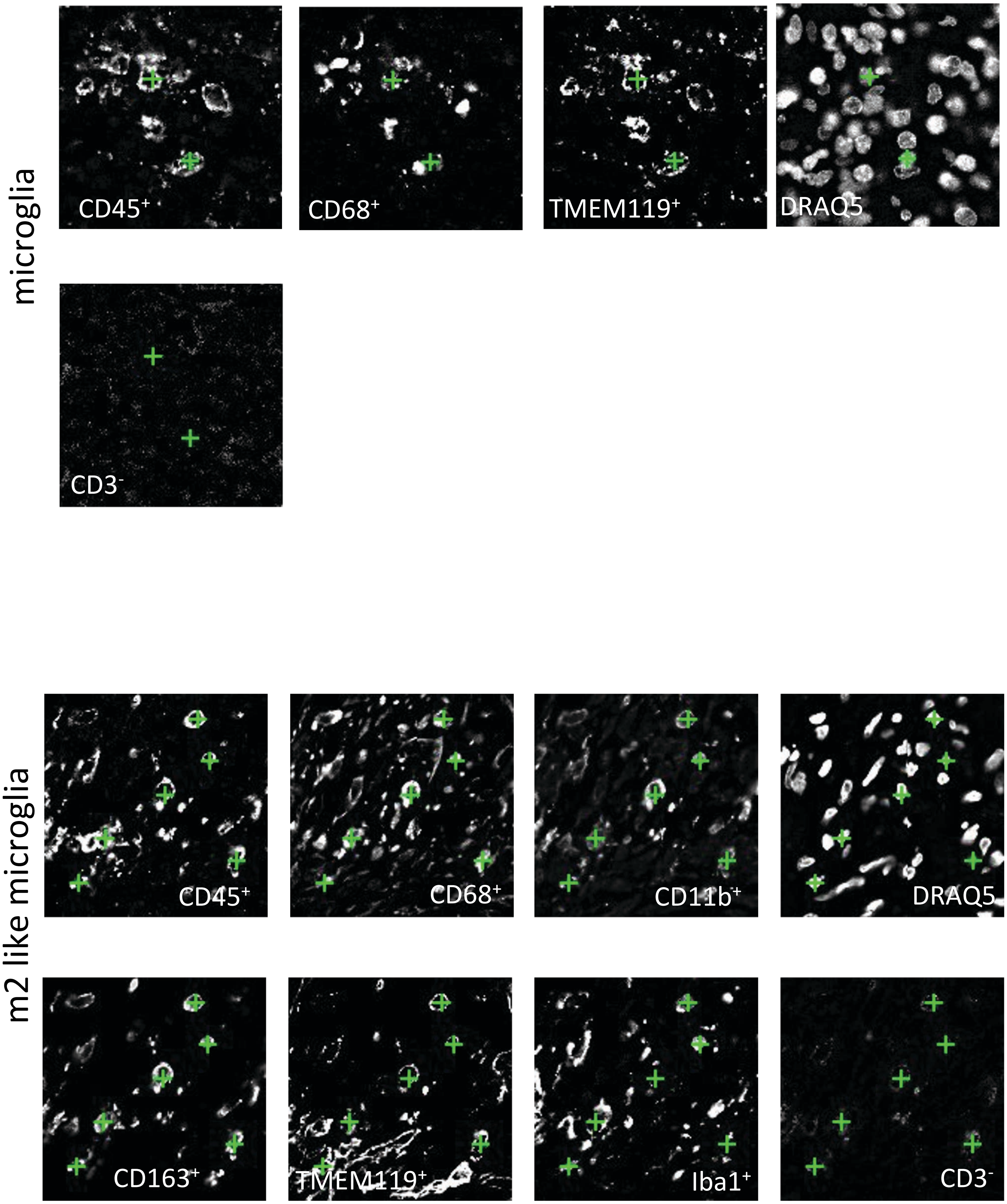
microglia
CD45⁺
CD68⁺
TMEM119⁺
DRAQ5
CD3⁻
m2 like microglia
CD45⁺
CD68⁺
CD11b⁺
DRAQ5
CD163⁺
TMEM119⁺
Iba1⁺
CD3⁻

**Figure S3**

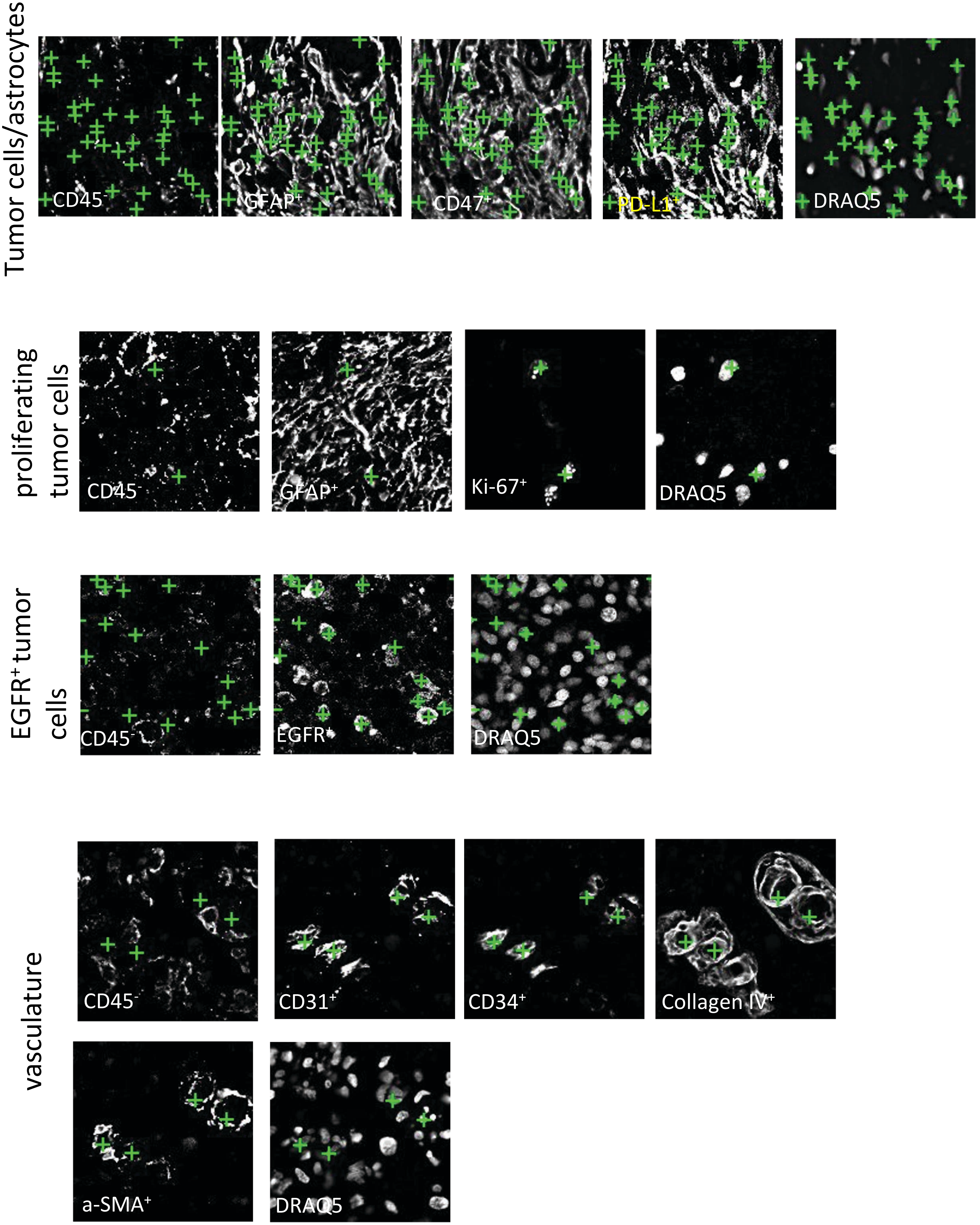
Tumor cells/astrocytes
CD45⁻
GFAP⁺
CD47⁺
PD-L1⁺
DRAQ5
proliferating tumor cells
CD45⁻
GFAP⁺
Ki-67⁺
DRAQ5
EGFR⁺ tumor cells
CD45⁻
EGFR⁺
DRAQ5
vasculature
CD45⁻
CD31⁺
CD34⁺
Collagen IV⁺
a-SMA⁺
DRAQ5

**Figure S3**

**representative center sample**

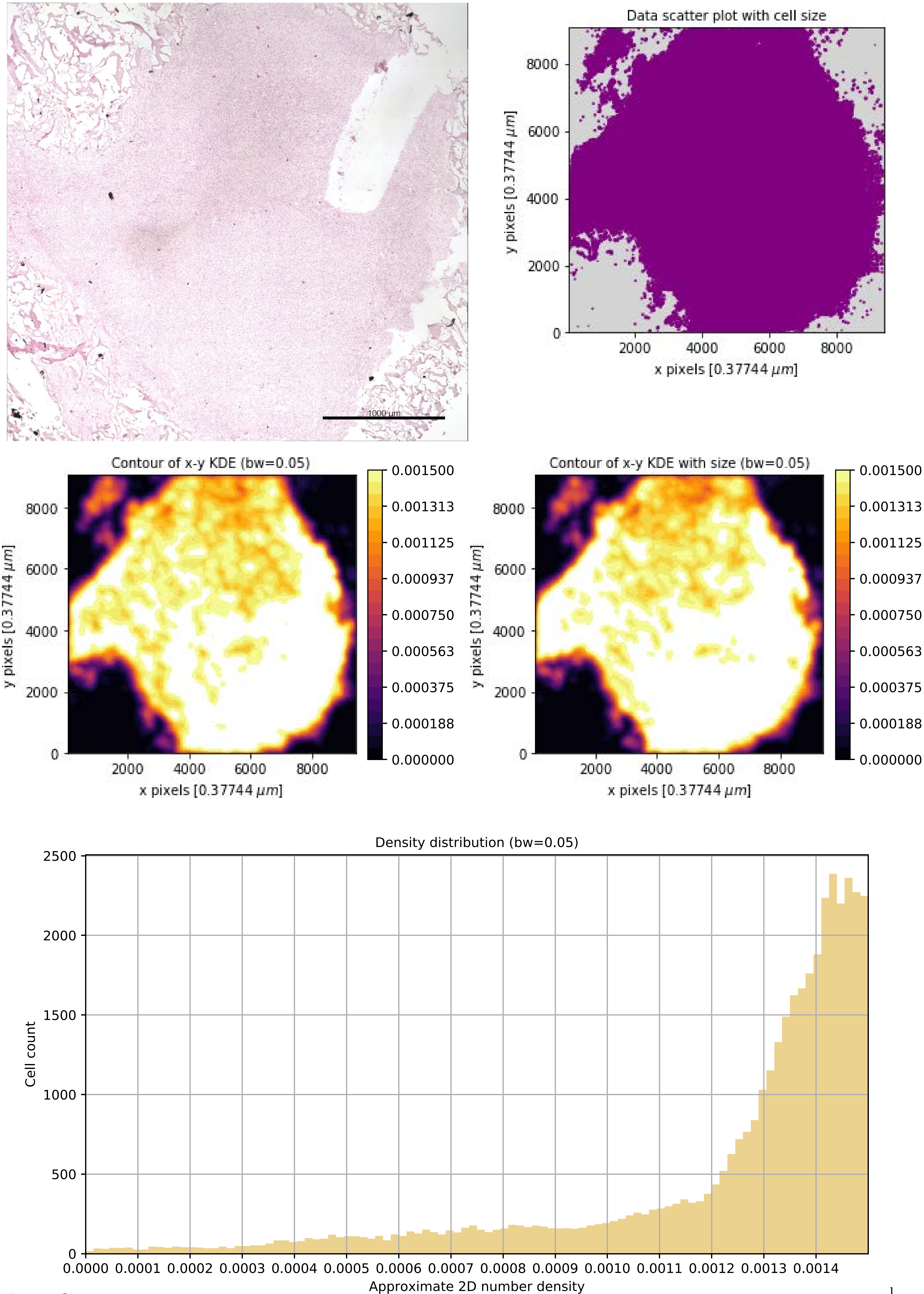


**Figure S4**

**representative periphery sample**

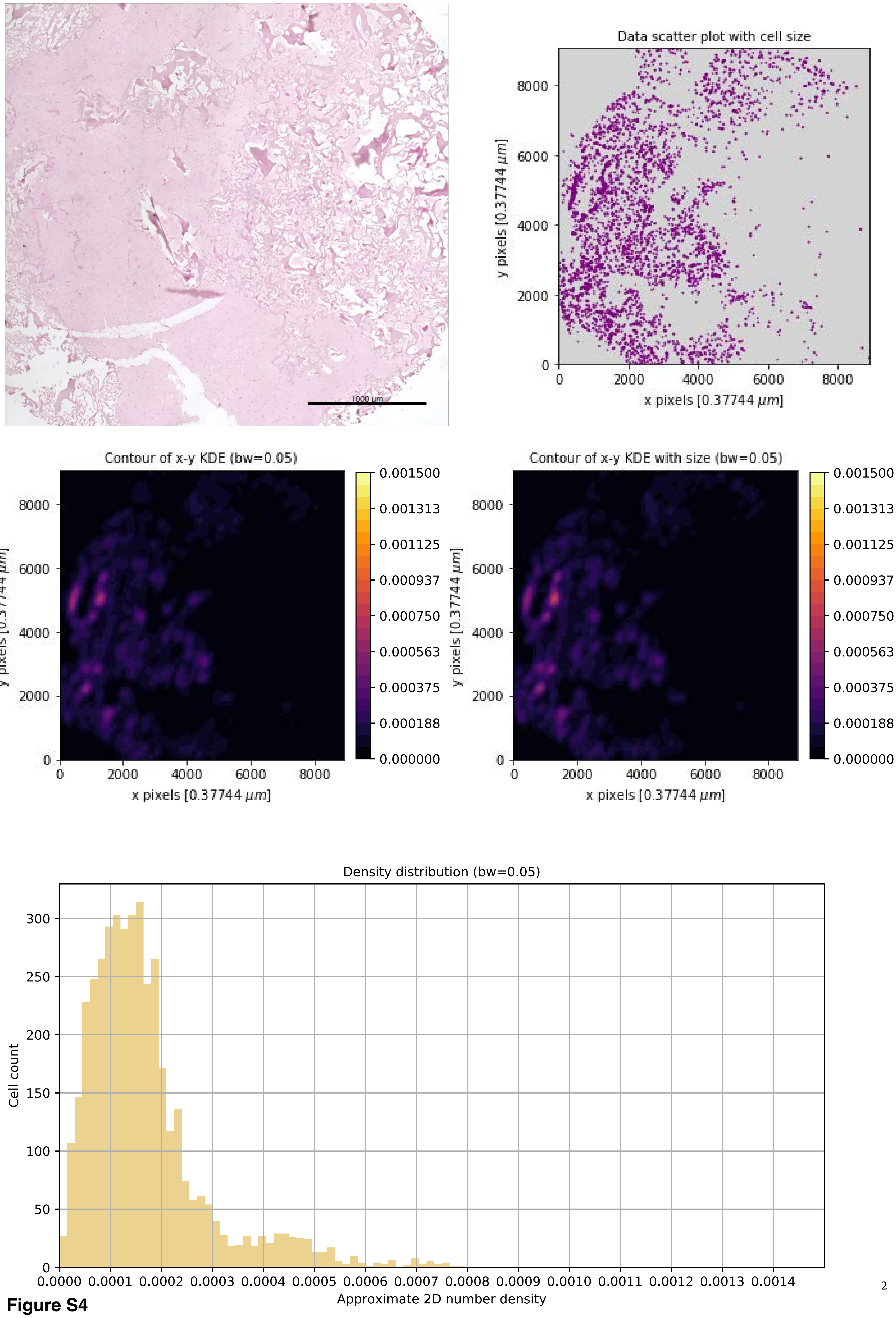


**Figure S4**

**Figure S4**

#### Overview all samples - density histograms

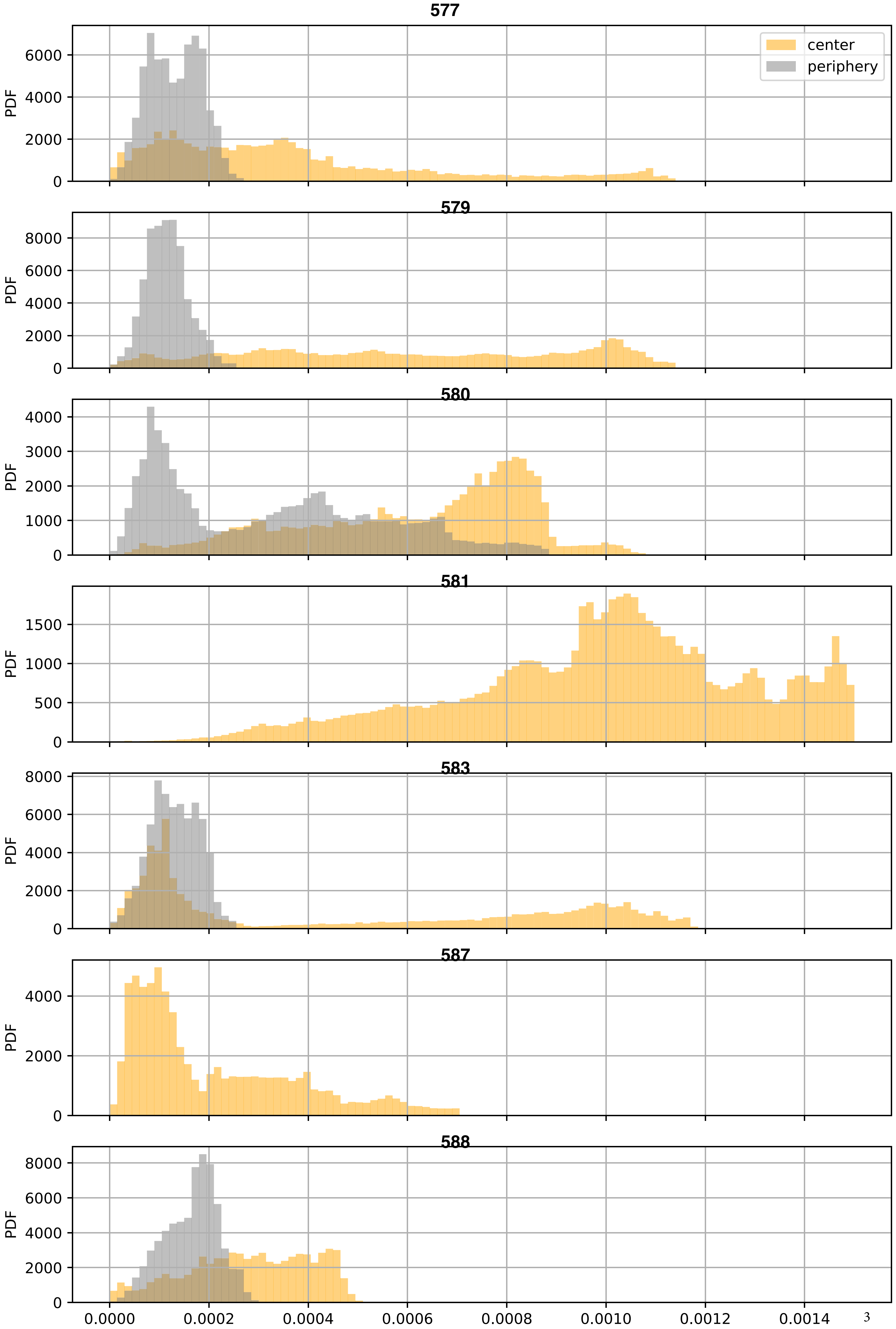

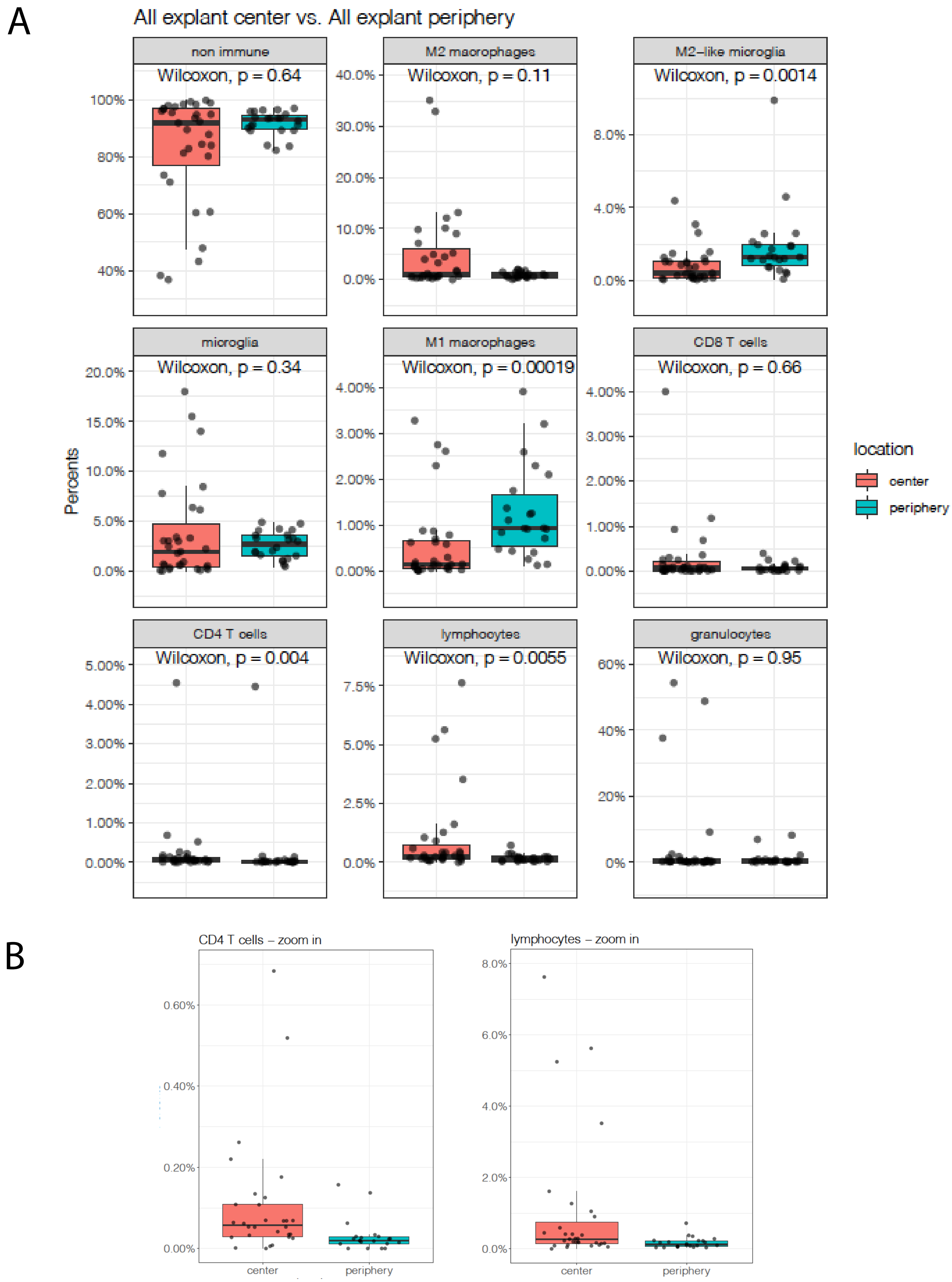
A
All explant center vs. All explant periphery
non immune
Wilcoxon, p = 0.64
M2 macrophages
Wilcoxon, p = 0.11
M2–like microglia
Wilcoxon, p = 0.0014
microglia
Wilcoxon, p = 0.34
M1 macrophages
Wilcoxon, p = 0.00019
CD8 T cells
Wilcoxon, p = 0.66
CD4 T cells
Wilcoxon, p = 0.004
lymphocytes
Wilcoxon, p = 0.0055
granulocytes
Wilcoxon, p = 0.95
Percents
location
center
periphery
B
CD4 T cells – zoom in
lymphocytes – zoom in
center
periphery

**Figure S5**

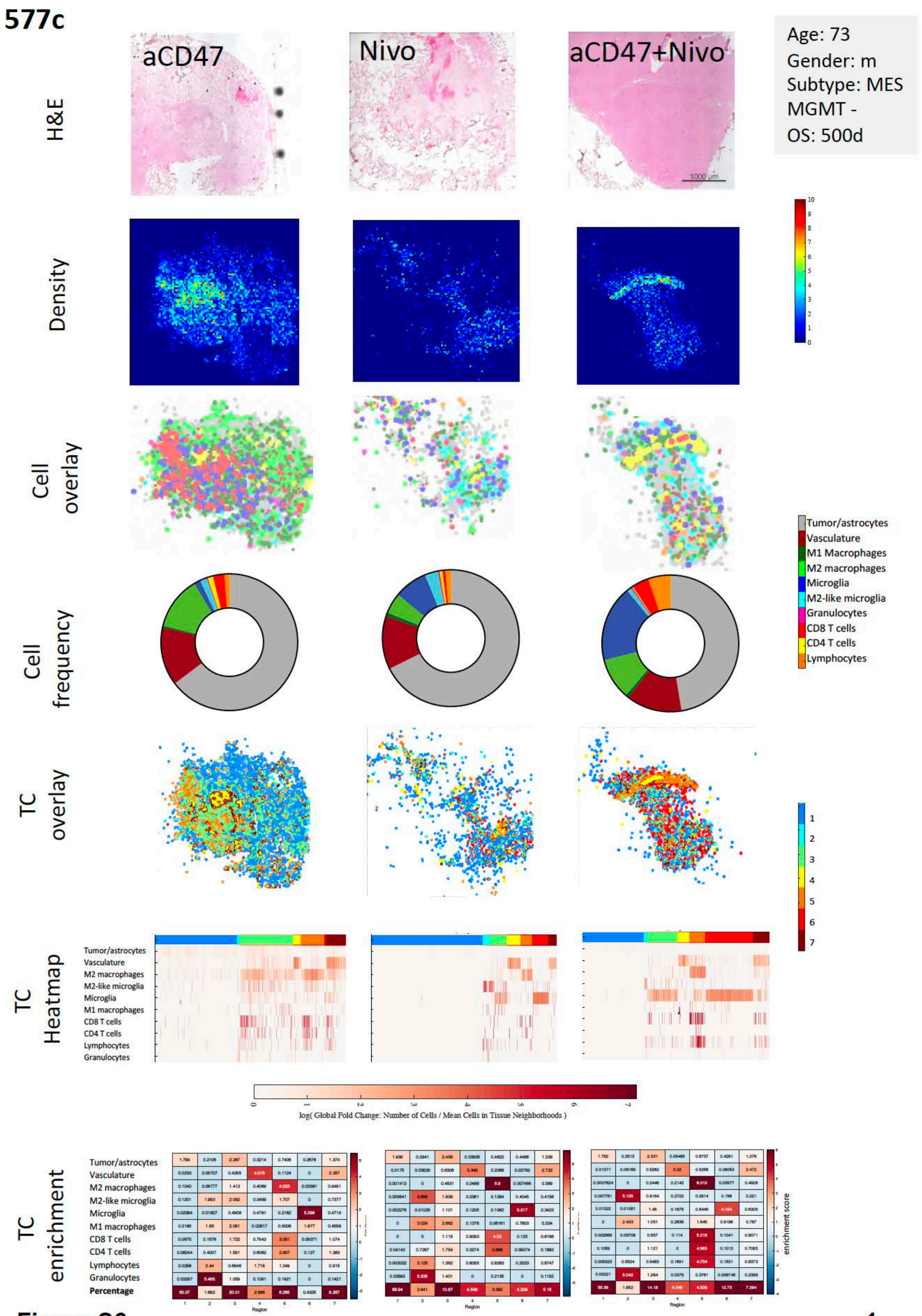
577c
aCD47
Nivo
aCD47+Nivo
Age: 73
Gender: m
Subtype: MES
MGMT -
OS: 500d
H&E
Density
Cell overlay
Cell frequency
TC overlay
TC Heatmap
TC enrichment
1000 µm
Tumor/astrocytes
Vasculature
M1 Macrophages
M2 macrophages
Microglia
M2-like microglia
Granulocytes
CD8 T cells
CD4 T cells
Lymphocytes
M1 macrophages
Percentage
log( Global Fold Change: Number of Cells / Mean Cells in Tissue Neighborhoods )
Region
enrichment score

Figure S6

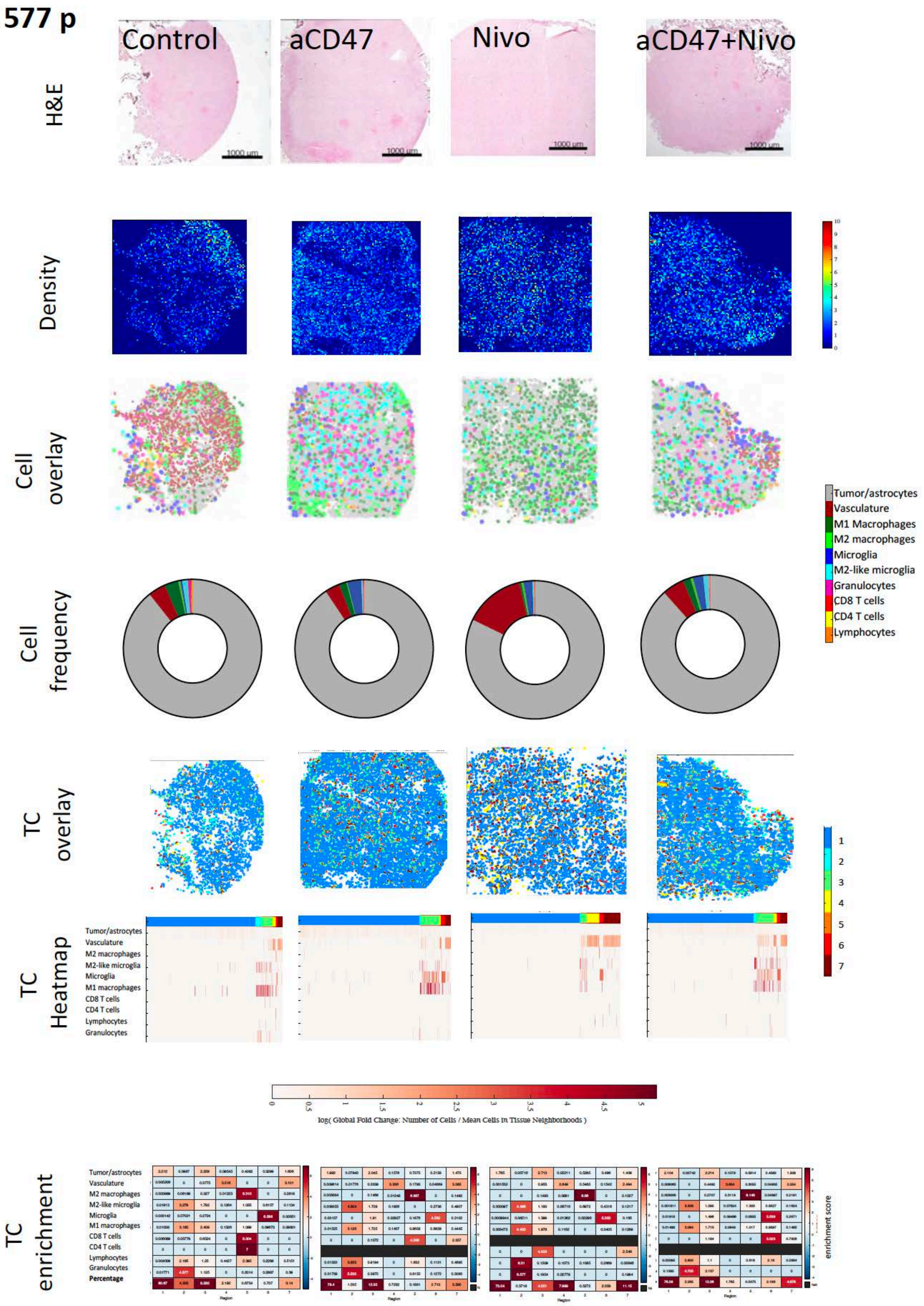

577 p
Control
aCD47
Nivo
aCD47+Nivo
H&E
1000 um
Density
Cell overlay
Cell frequency
Tumor/astrocytes
Vasculature
M1 Macrophages
M2 macrophages
Microglia
M2-like microglia
Granulocytes
CD8 T cells
CD4 T cells
Lymphocytes
TC overlay
TC Heatmap
M2 macrophages
M2-like microglia
M1 macrophages
log( Global Fold Change: Number of Cells / Mean Cells in Tissue Neighborhoods )
TC enrichment
Percentage
Region
enrichment score


**Figure S6**

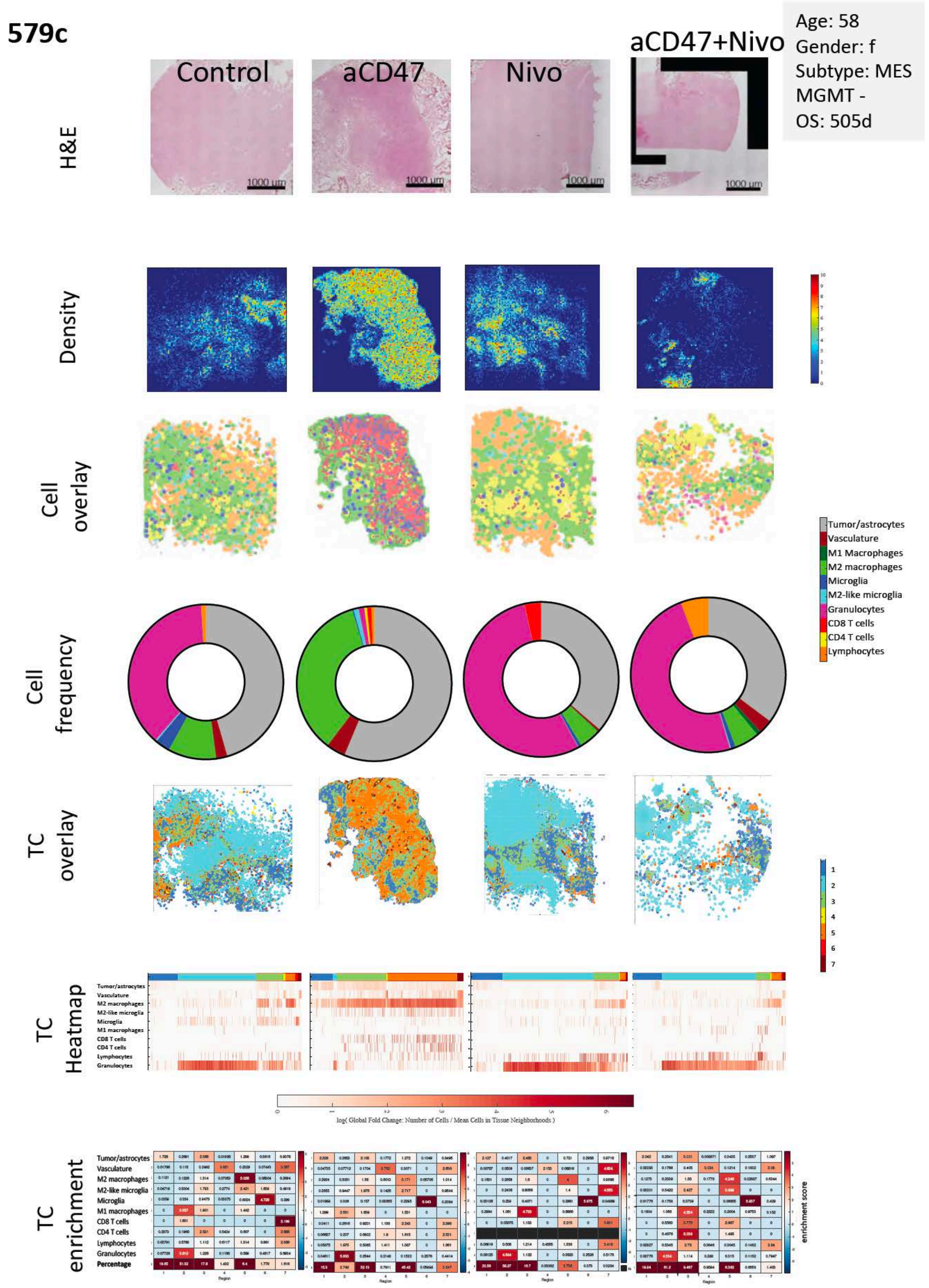

579c
Age: 58
Gender: f
Subtype: MES
MGMT -
OS: 505d
Control
aCD47
Nivo
aCD47+Nivo
H&E
1000 um
Density
Cell overlay
Cell frequency
TC overlay
TC Heatmap
TC enrichment
Tumor/astrocytes
Vasculature
M1 Macrophages
M2 macrophages
Microglia
M2-like microglia
Granulocytes
CD8 T cells
CD4 T cells
Lymphocytes
Percentage
log( Global Fold Change: Number of Cells / Mean Cells in Tissue Neighborhoods )
Region
enrichment score


Figure S6

**579 p**

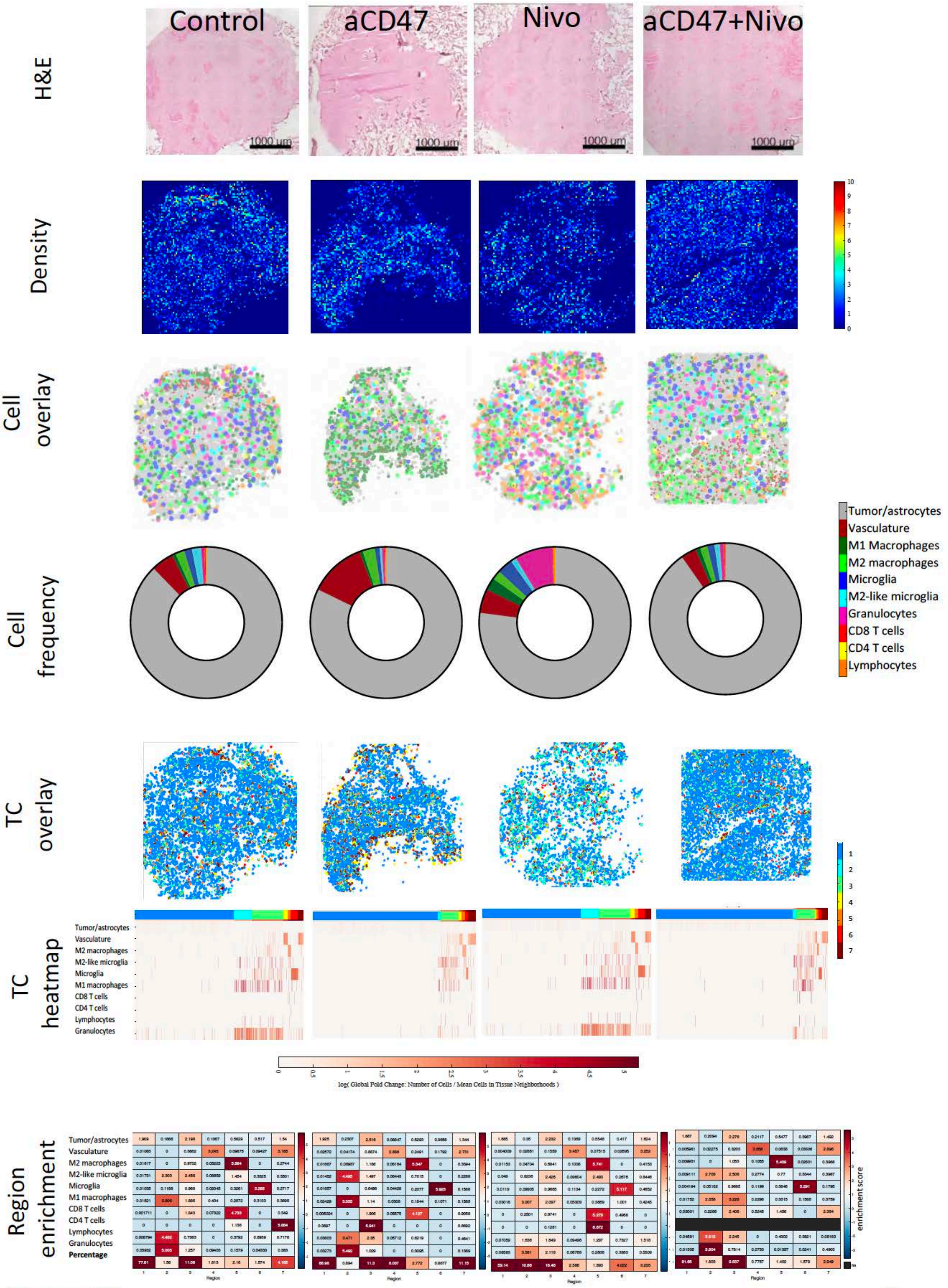


**Figure S6**

## 580 c

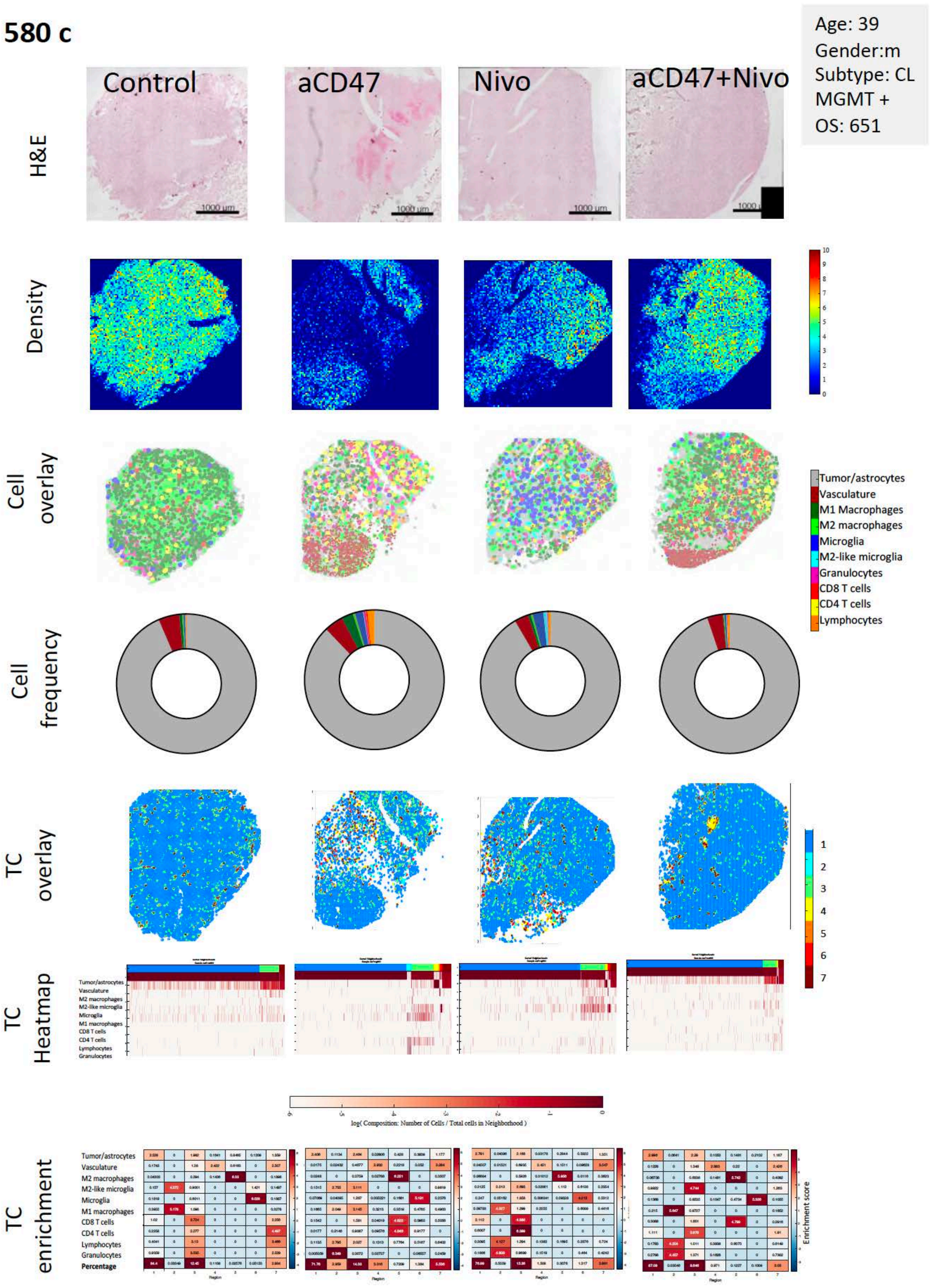


**Figure S6**

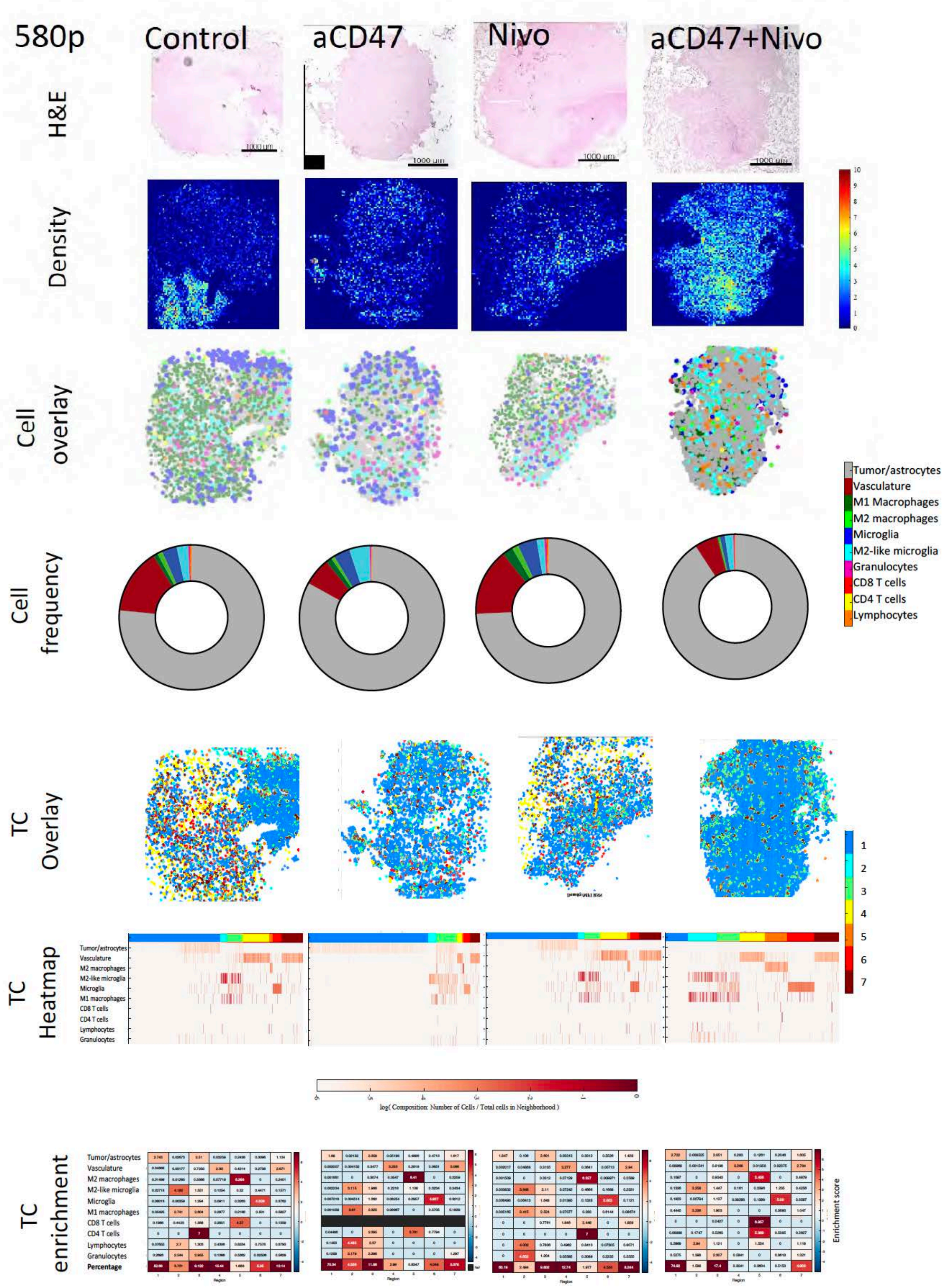
580p
Control
aCD47
Nivo
aCD47+Nivo
H&E
1000 µm
Density
Cell overlay
Cell frequency
Tumor/astrocytes
Vasculature
M1 Macrophages
M2 macrophages
Microglia
M2-like microglia
Granulocytes
CD8 T cells
CD4 T cells
Lymphocytes
TC Overlay
TC Heatmap
Tumor/astrocytes
Vasculature
M2 macrophages
M2-like microglia
Microglia
M1 macrophages
CD8 T cells
CD4 T cells
Lymphocytes
Granulocytes
log( Composition: Number of Cells / Total cells in Neighborhood )
TC enrichment
Tumor/astrocytes
Vasculature
M2 macrophages
M2-like microglia
Microglia
M1 macrophages
CD8 T cells
CD4 T cells
Lymphocytes
Granulocytes
Percentage
Region
Enrichment score

**Figure S6**

## 581c

Age: 59
Gender: m
Subtype: CL
MGMT +
OS: >700

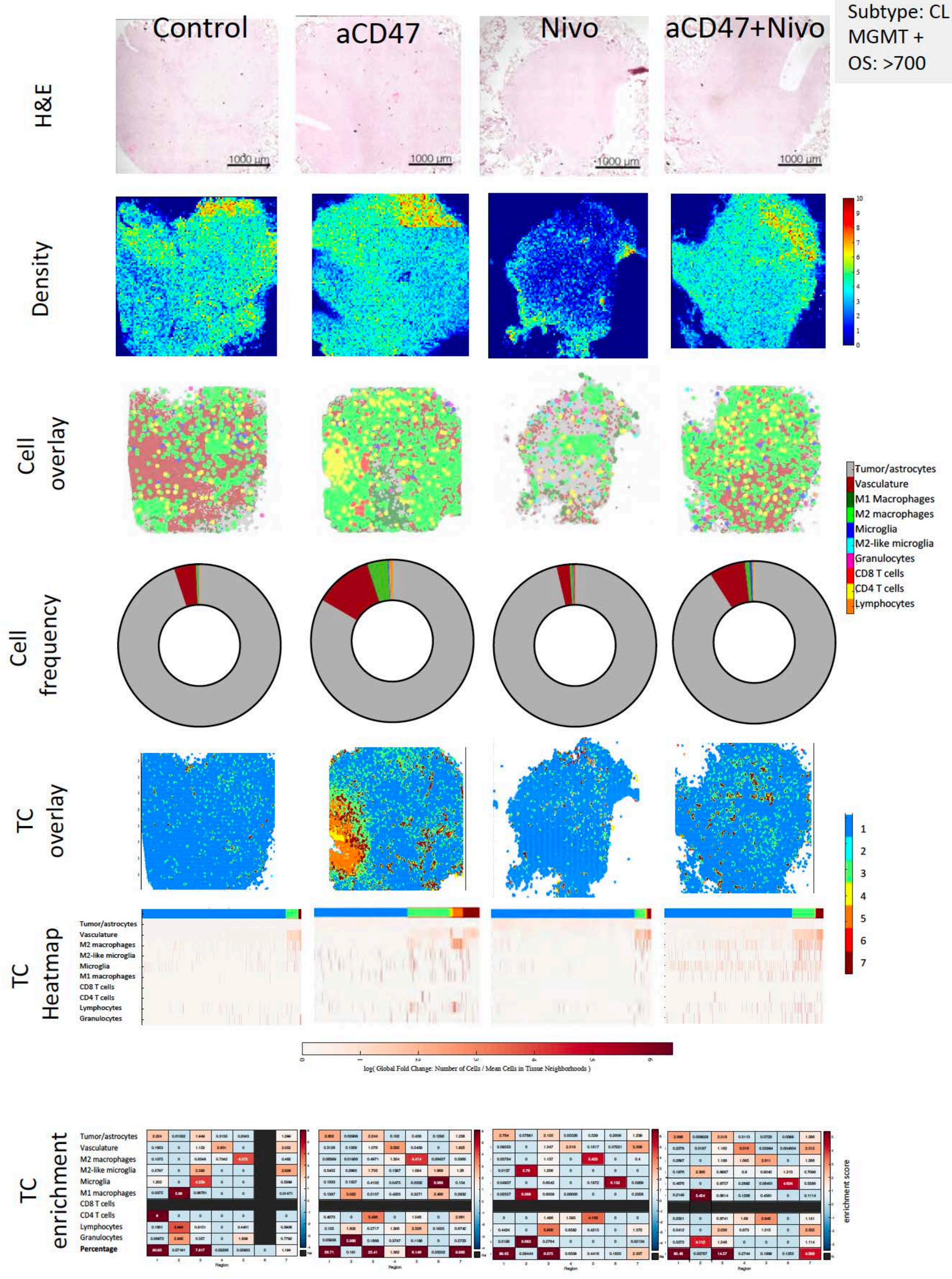


Figure S6

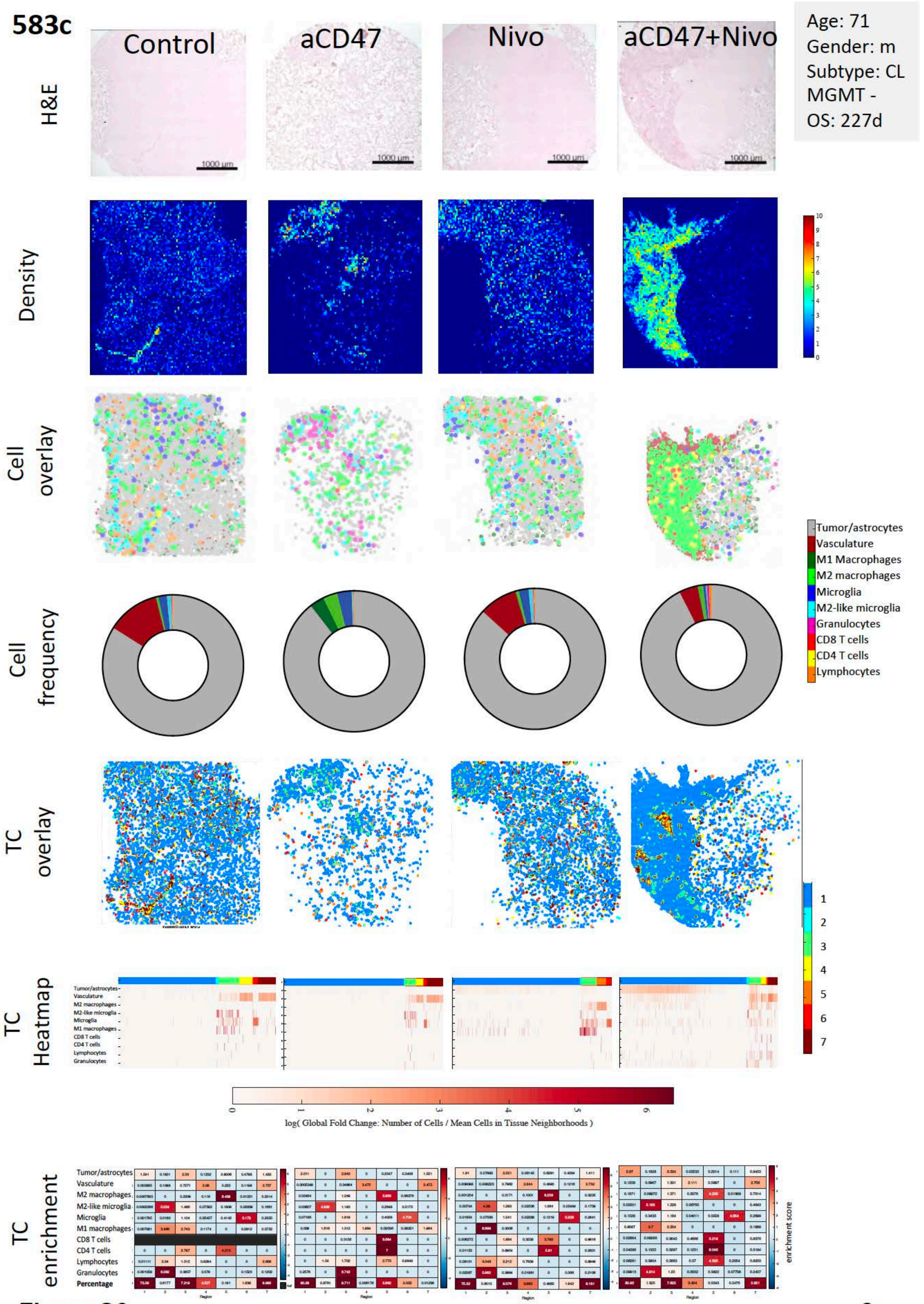
583c
Control
aCD47
Nivo
aCD47+Nivo
Age: 71
Gender: m
Subtype: CL
MGMT -
OS: 227d
H&E
Density
Cell overlay
Cell frequency
TC overlay
TC Heatmap
TC enrichment
Tumor/astrocytes
Vasculature
M1 Macrophages
M2 macrophages
Microglia
M2-like microglia
Granulocytes
CD8 T cells
CD4 T cells
Lymphocytes
log( Global Fold Change: Number of Cells / Mean Cells in Tissue Neighborhoods )
Percentage
enrichment score

**Figure S6**

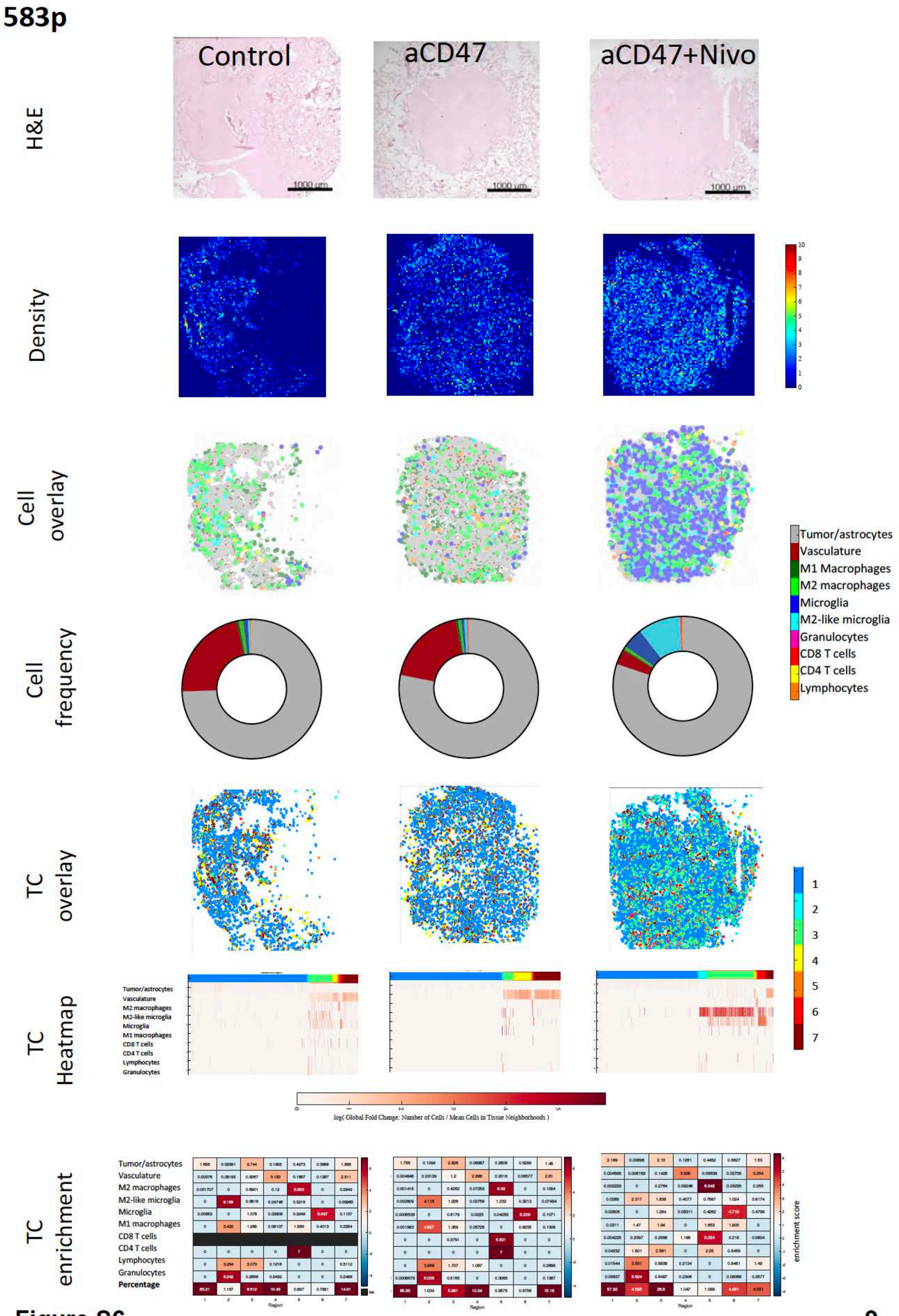
583p
Control
aCD47
aCD47+Nivo
H&E
Density
Cell overlay
Cell frequency
TC overlay
TC Heatmap
TC enrichment
Tumor/astrocytes
Vasculature
M1 Macrophages
M2 macrophages
Microglia
M2-like microglia
Granulocytes
CD8 T cells
CD4 T cells
Lymphocytes
Percentage
Region
enrichment score

Figure S6

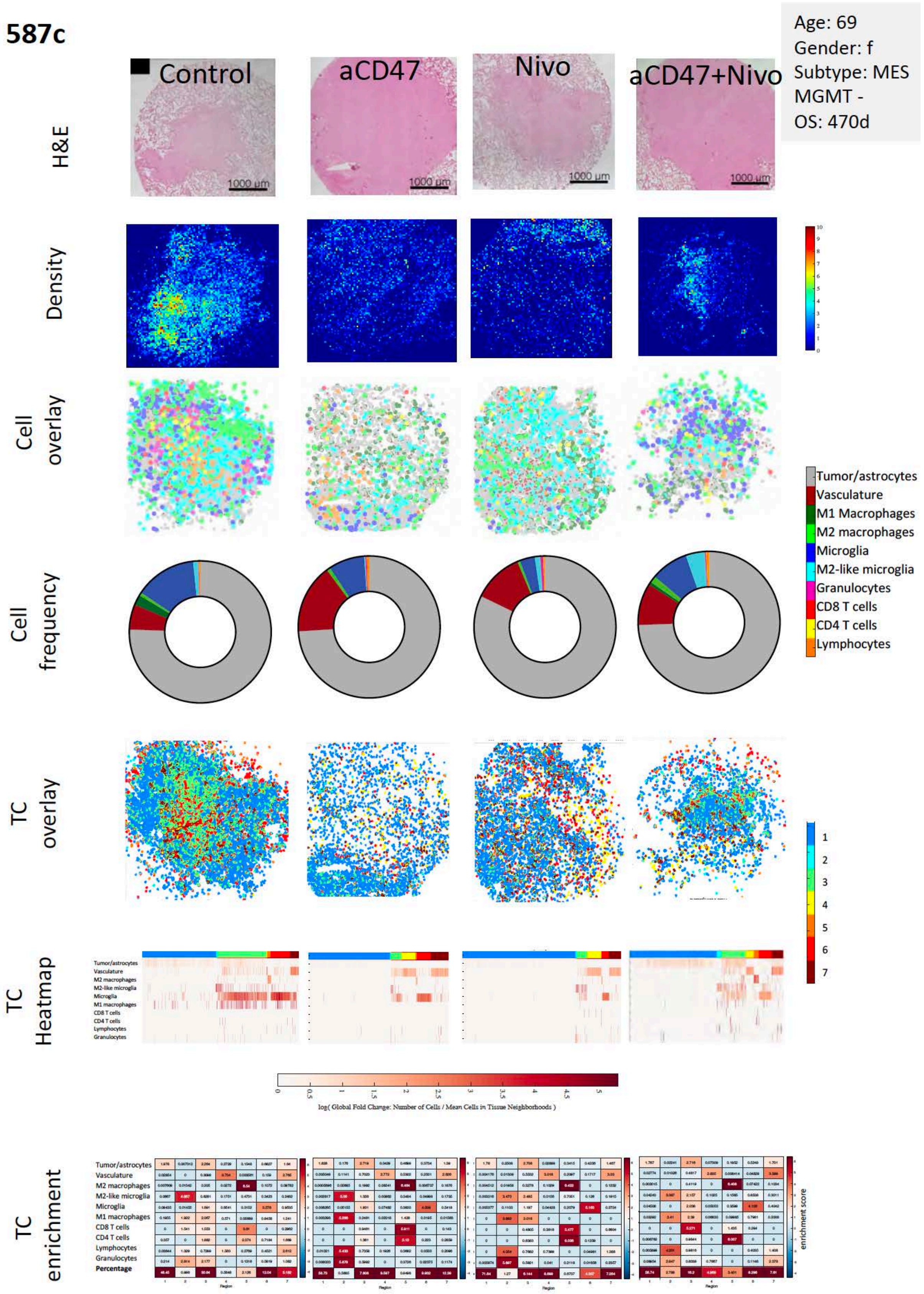
587c
Age: 69
Gender: f
Subtype: MES
MGMT -
OS: 470d
Control
aCD47
Nivo
aCD47+Nivo
H&E
1000 µm
Density
Cell overlay
Cell frequency
Tumor/astrocytes
Vasculature
M1 Macrophages
M2 macrophages
Microglia
M2-like microglia
Granulocytes
CD8 T cells
CD4 T cells
Lymphocytes
TC overlay
TC Heatmap
log( Global Fold Change: Number of Cells / Mean Cells in Tissue Neighborhoods )
TC enrichment
Percentage
Region
enrichment score

**Figure S6**

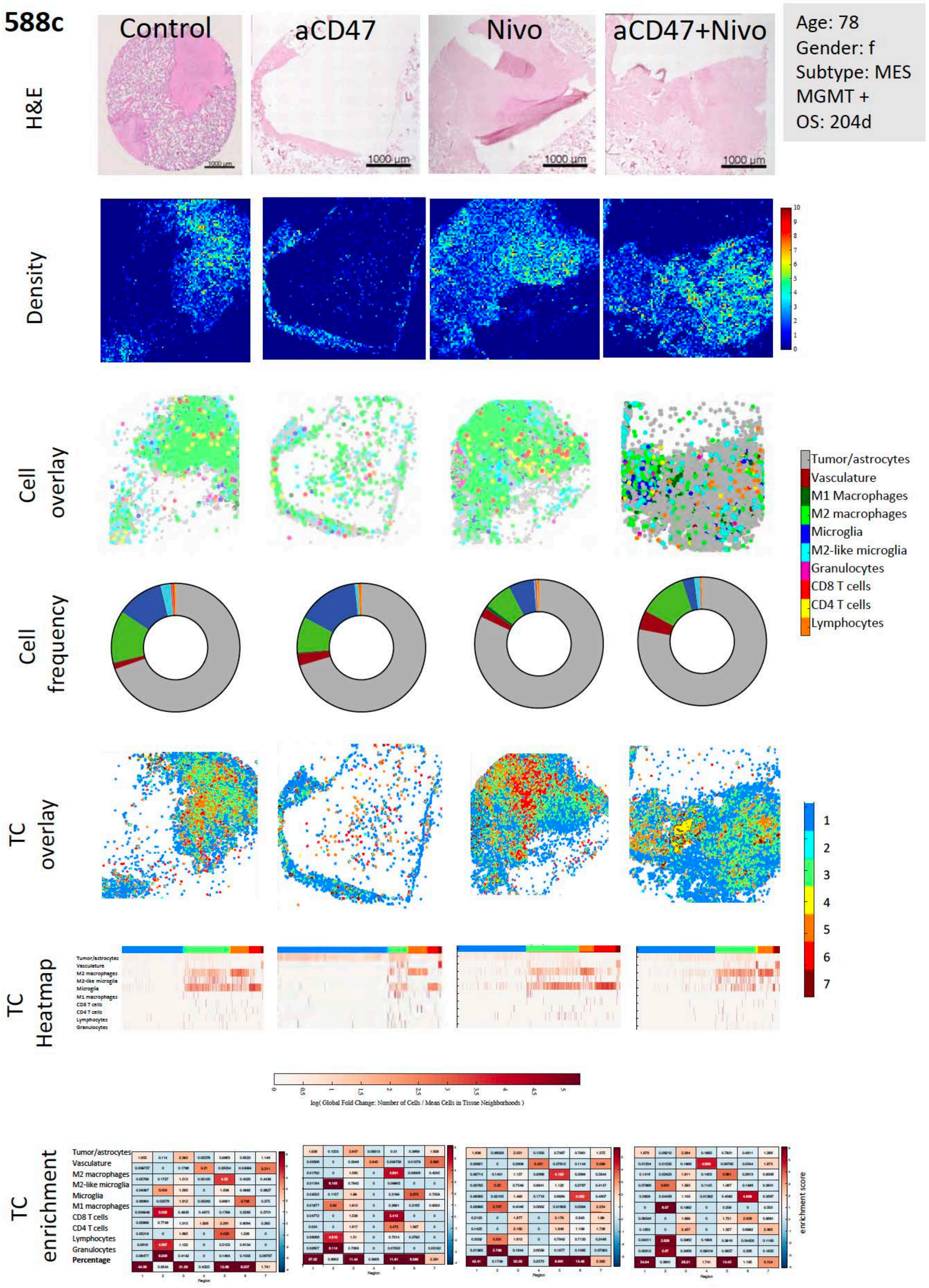
588c
Control
aCD47
Nivo
aCD47+Nivo
Age: 78
Gender: f
Subtype: MES
MGMT +
OS: 204d
H&E
1000 µm
Density
Cell overlay
Tumor/astrocytes
Vasculature
M1 Macrophages
M2 macrophages
Microglia
M2-like microglia
Granulocytes
CD8 T cells
CD4 T cells
Lymphocytes
Cell frequency
TC overlay
1
2
3
4
5
6
7
TC Heatmap
log( Global Fold Change: Number of Cells / Mean Cells in Tissue Neighborhoods )
TC enrichment
Percentage
Region
enrichment score

Figure S6 

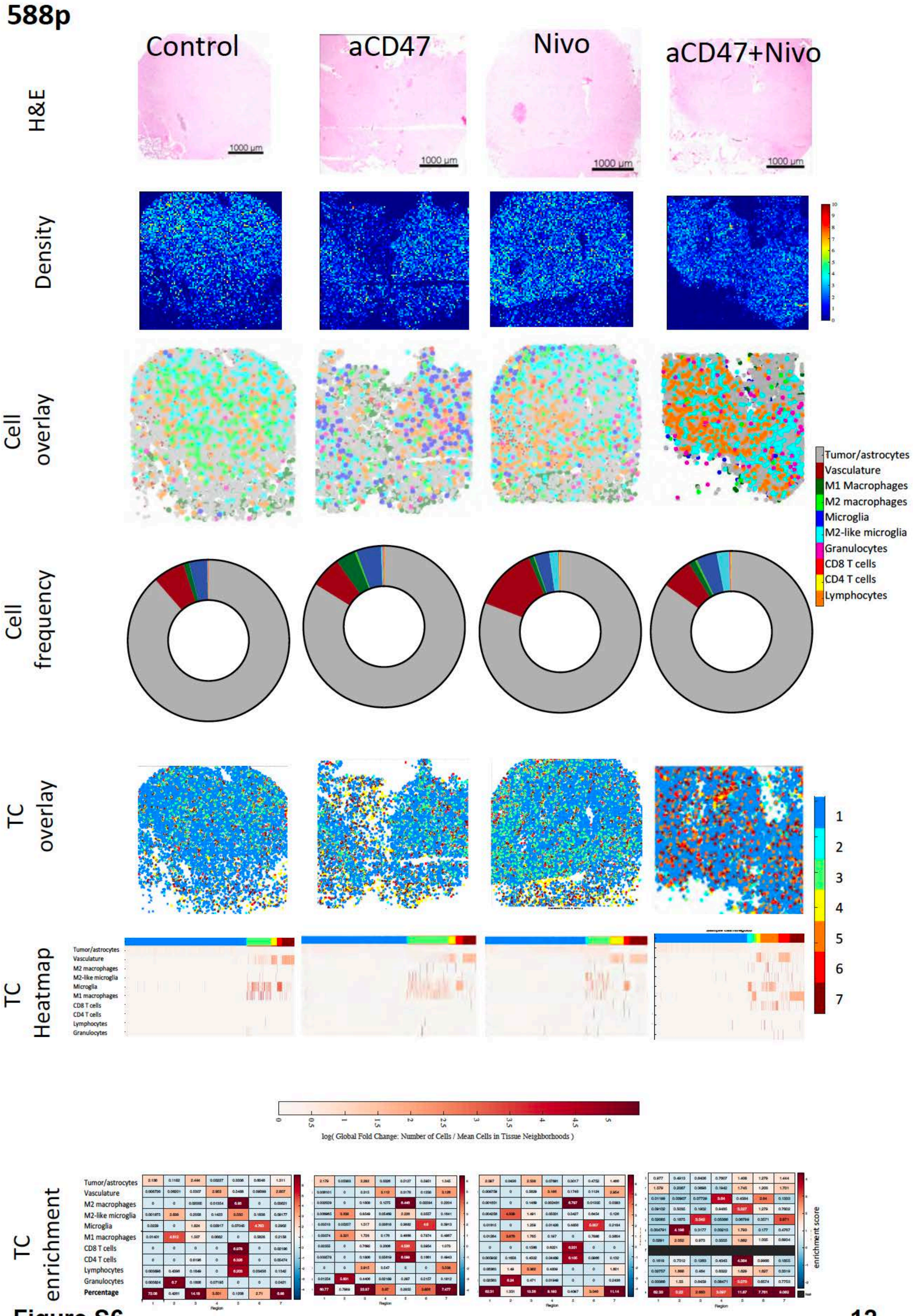
588p
Control
aCD47
Nivo
aCD47+Nivo
H&E
1000 μm
Density
Cell overlay
Tumor/astrocytes
Vasculature
M1 Macrophages
M2 macrophages
Microglia
M2-like microglia
Granulocytes
CD8 T cells
CD4 T cells
Lymphocytes
Cell frequency
TC overlay
1
2
3
4
5
6
7
TC Heatmap
Tumor/astrocytes
Vasculature
M2 macrophages
M2-like microglia
Microglia
M1 macrophages
CD8 T cells
CD4 T cells
Lymphocytes
Granulocytes
log( Global Fold Change: Number of Cells / Mean Cells in Tissue Neighborhoods )
TC enrichment
Percentage
Region
enrichment score

Figure S6

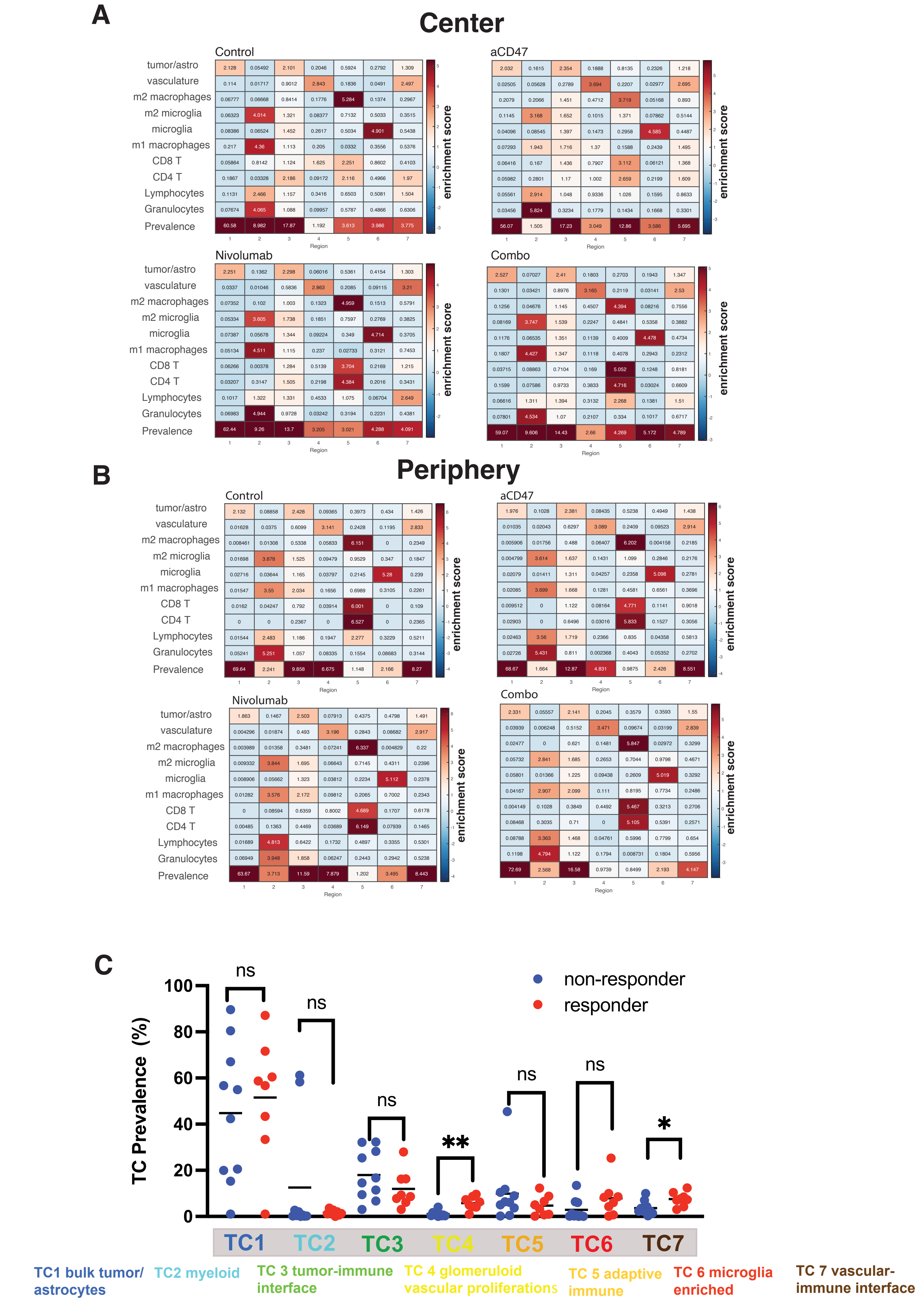
A
Center
Control
aCD47
Nivolumab
Combo
tumor/astro
vasculature
m2 macrophages
m2 microglia
microglia
m1 macrophages
CD8 T
CD4 T
Lymphocytes
Granulocytes
Prevalence
Region
enrichment score
B
Periphery
C
TC Prevalence (%)
non-responder
responder
ns
**
*
TC1
TC2
TC3
TC4
TC5
TC6
TC7
TC1 bulk tumor/ astrocytes
TC2 myeloid
TC 3 tumor-immune interface
TC 4 glomeruloid vascular proliferations
TC 5 adaptive immune
TC 6 microglia enriched
TC 7 vascular-immune interface

Figure S7

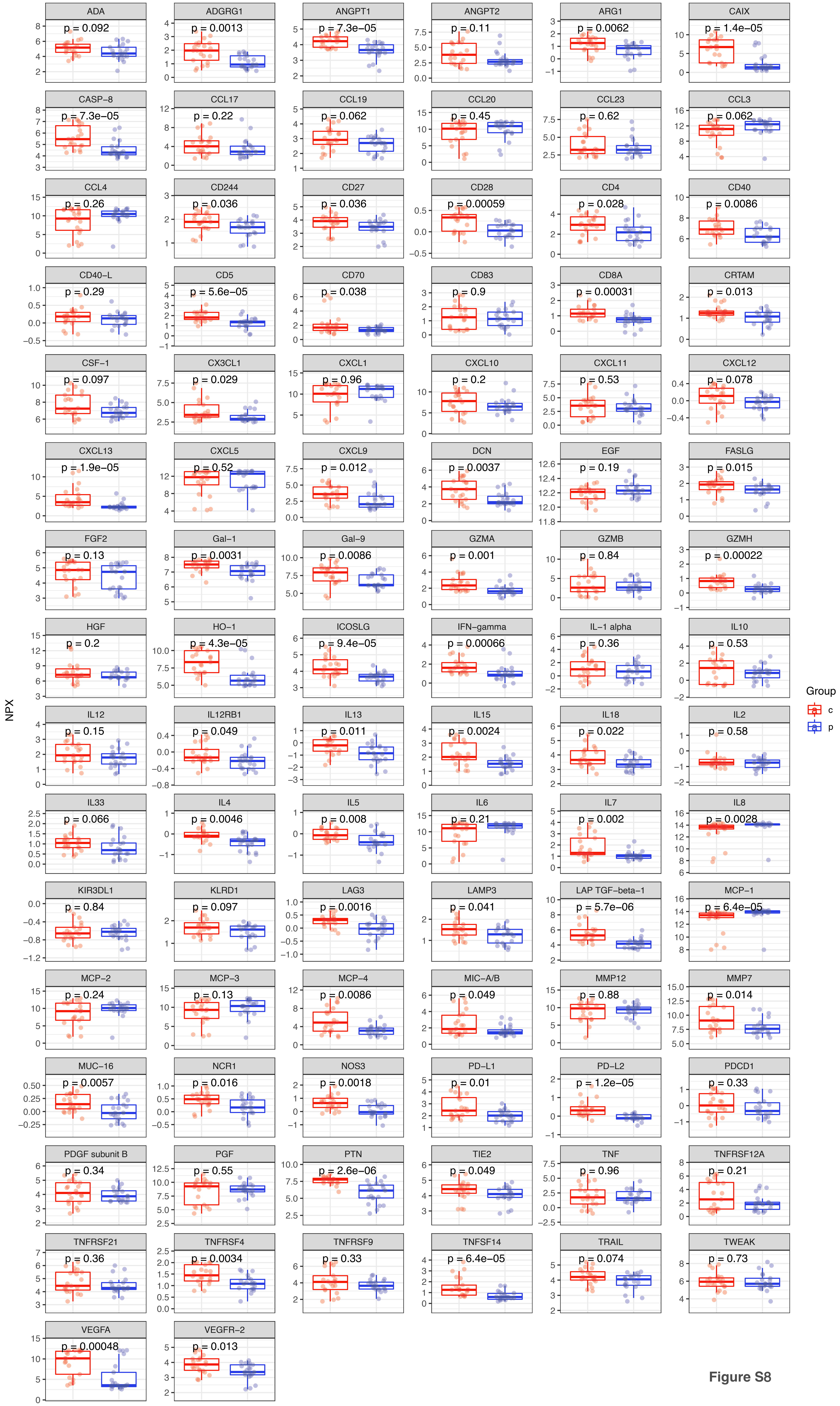
NPX
Group
c
p
ADA p = 0.092
ADGRG1 p = 0.0013
ANGPT1 p = 7.3e−05
ANGPT2 p = 0.11
ARG1 p = 0.0062
CAIX p = 1.4e−05
CASP−8 p = 7.3e−05
CCL17 p = 0.22
CCL19 p = 0.062
CCL20 p = 0.45
CCL23 p = 0.62
CCL3 p = 0.062
CCL4 p = 0.26
CD244 p = 0.036
CD27 p = 0.036
CD28 p = 0.00059
CD4 p = 0.028
CD40 p = 0.0086
CD40−L p = 0.29
CD5 p = 5.6e−05
CD70 p = 0.038
CD83 p = 0.9
CD8A p = 0.00031
CRTAM p = 0.013
CSF−1 p = 0.097
CX3CL1 p = 0.029
CXCL1 p = 0.96
CXCL10 p = 0.2
CXCL11 p = 0.53
CXCL12 p = 0.078
CXCL13 p = 1.9e−05
CXCL5 p = 0.52
CXCL9 p = 0.012
DCN p = 0.0037
EGF p = 0.19
FASLG p = 0.015
FGF2 p = 0.13
Gal−1 p = 0.0031
Gal−9 p = 0.0086
GZMA p = 0.001
GZMB p = 0.84
GZMH p = 0.00022
HGF p = 0.2
HO−1 p = 4.3e−05
ICOSLG p = 9.4e−05
IFN−gamma p = 0.00066
IL−1 alpha p = 0.36
IL10 p = 0.53
IL12 p = 0.15
IL12RB1 p = 0.049
IL13 p = 0.011
IL15 p = 0.0024
IL18 p = 0.022
IL2 p = 0.58
IL33 p = 0.066
IL4 p = 0.0046
IL5 p = 0.008
IL6 p = 0.21
IL7 p = 0.002
IL8 p = 0.0028
KIR3DL1 p = 0.84
KLRD1 p = 0.097
LAG3 p = 0.0016
LAMP3 p = 0.041
LAP TGF−beta−1 p = 5.7e−06
MCP−1 p = 6.4e−05
MCP−2 p = 0.24
MCP−3 p = 0.13
MCP−4 p = 0.0086
MIC−A/B p = 0.049
MMP12 p = 0.88
MMP7 p = 0.014
MUC−16 p = 0.0057
NCR1 p = 0.016
NOS3 p = 0.0018
PD−L1 p = 0.01
PD−L2 p = 1.2e−05
PDCD1 p = 0.33
PDGF subunit B p = 0.34
PGF p = 0.55
PTN p = 2.6e−06
TIE2 p = 0.049
TNF p = 0.96
TNFRSF12A p = 0.21
TNFRSF21 p = 0.36
TNFRSF4 p = 0.0034
TNFRSF9 p = 0.33
TNFSF14 p = 6.4e−05
TRAIL p = 0.074
TWEAK p = 0.73
VEGFA p = 0.00048
VEGFR−2 p = 0.013

**Figure S8**

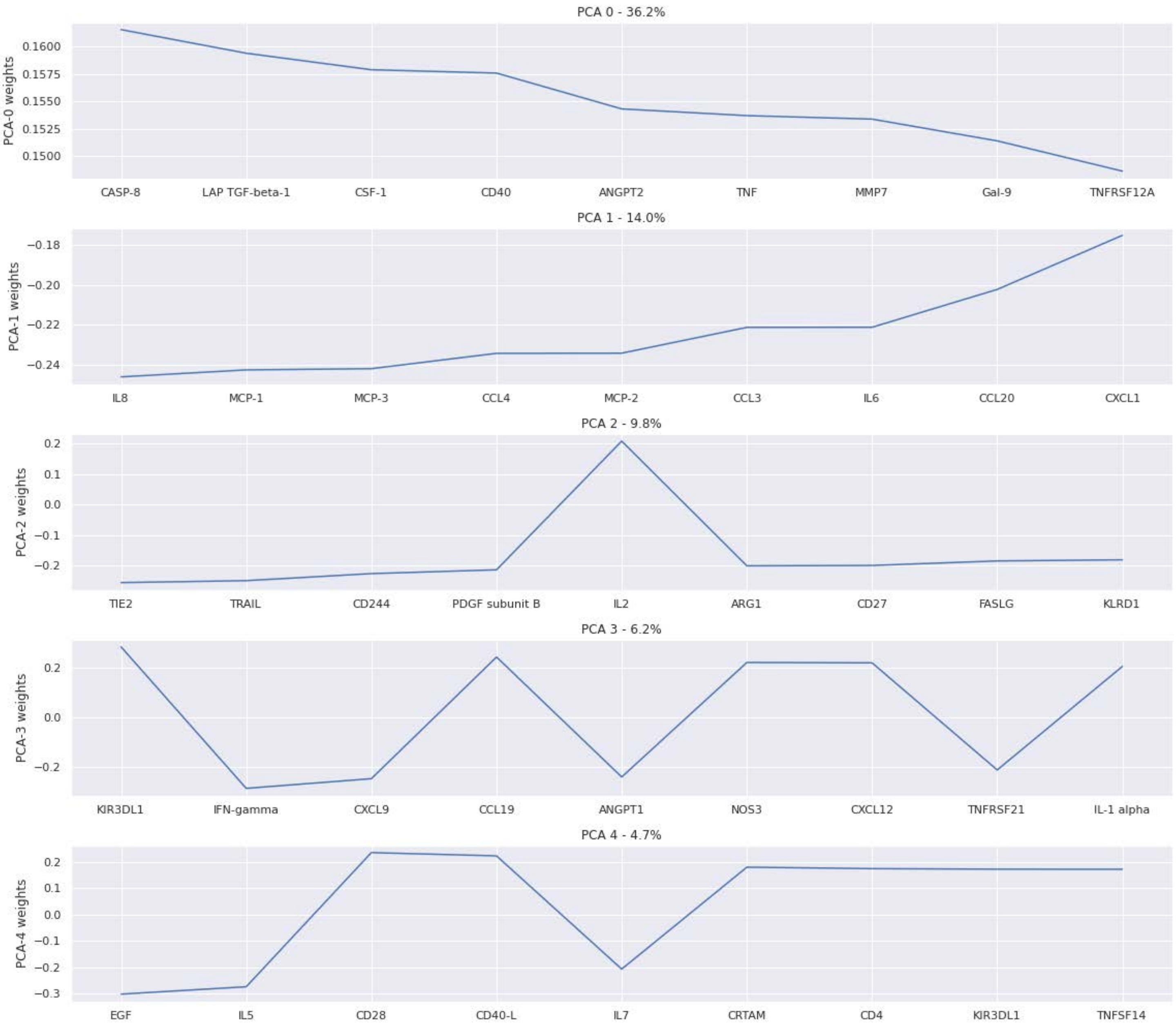
PCA 0 - 36.2%
PCA-0 weights
0.1600
0.1575
0.1550
0.1525
0.1500
CASP-8
LAP TGF-beta-1
CSF-1
CD40
ANGPT2
TNF
MMP7
Gal-9
TNFRSF12A
PCA 1 - 14.0%
PCA-1 weights
−0.18
−0.20
−0.22
−0.24
IL8
MCP-1
MCP-3
CCL4
MCP-2
CCL3
IL6
CCL20
CXCL1
PCA 2 - 9.8%
PCA-2 weights
0.2
0.1
0.0
−0.1
−0.2
TIE2
TRAIL
CD244
PDGF subunit B
IL2
ARG1
CD27
FASLG
KLRD1
PCA 3 - 6.2%
PCA-3 weights
0.2
0.0
−0.2
KIR3DL1
IFN-gamma
CXCL9
CCL19
ANGPT1
NOS3
CXCL12
TNFRSF21
IL-1 alpha
PCA 4 - 4.7%
PCA-4 weights
0.2
0.1
0.0
−0.1
−0.2
−0.3
EGF
IL5
CD28
CD40-L
IL7
CRTAM
CD4
KIR3DL1
TNFSF14

**Figure S9**

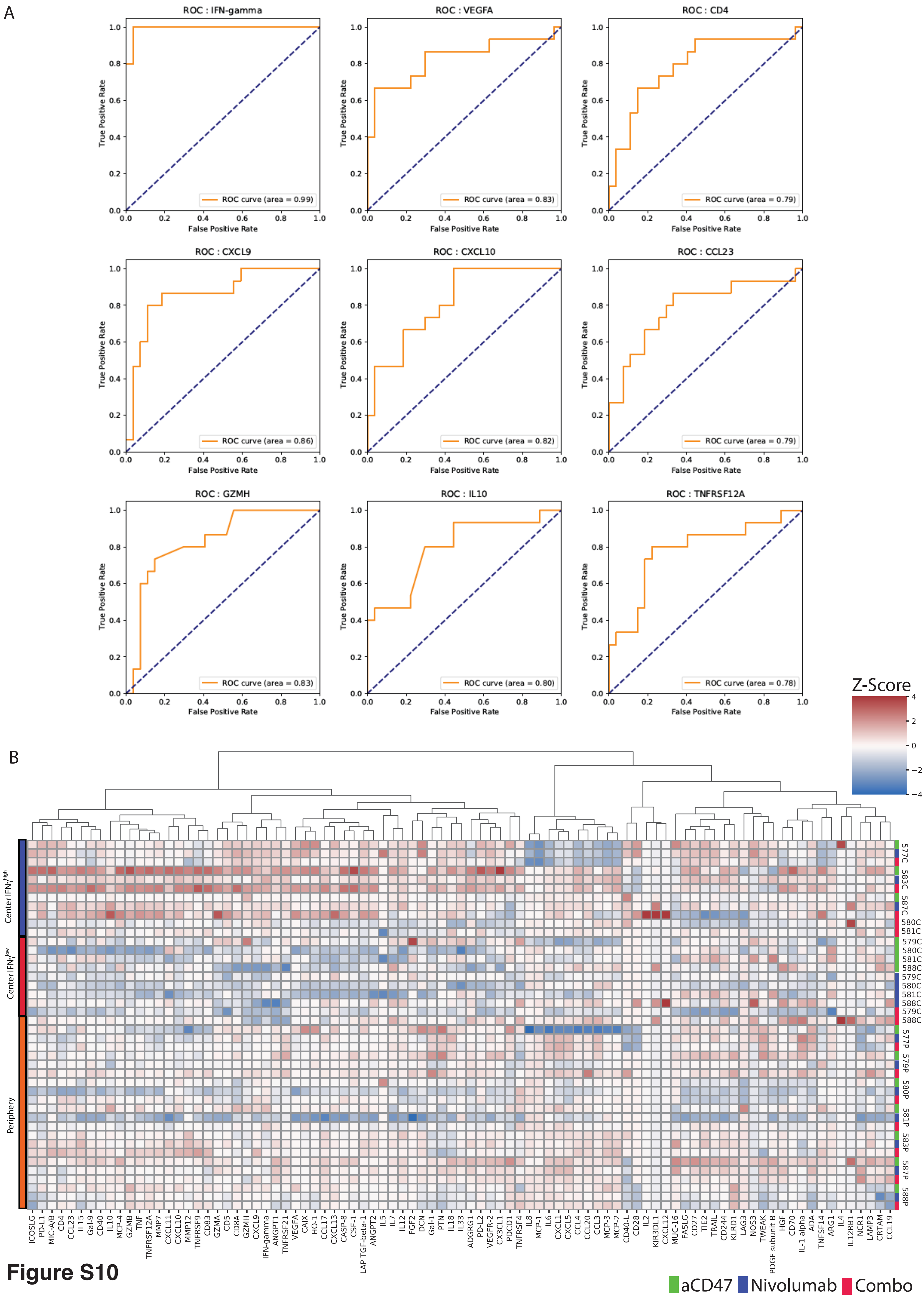
A
ROC : IFN-gamma
ROC curve (area = 0.99)
ROC : VEGFA
ROC curve (area = 0.83)
ROC : CD4
ROC curve (area = 0.79)
ROC : CXCL9
ROC curve (area = 0.86)
ROC : CXCL10
ROC curve (area = 0.82)
ROC : CCL23
ROC curve (area = 0.79)
ROC : GZMH
ROC curve (area = 0.83)
ROC : IL10
ROC curve (area = 0.80)
ROC : TNFRSF12A
ROC curve (area = 0.78)
True Positive Rate
False Positive Rate
B
Z-Score
Center IFNγ high
Center IFNγ low
Periphery
577C
583C
587C
580C
581C
579C
580C
581C
588C
579C
580C
581C
588C
579C
588C
577P
579P
580P
581P
583P
587P
588P
ICOSLG
PD-L1
MIC-A/B
CD4
CCL23
IL15
Gal-9
CD40
IL10
MCP-4
GZMB
TNF
TNFRSF12A
MMP7
CXCL11
CXCL10
MMP12
TNFRSF9
CD83
GZMA
CD5
CD8A
GZMH
CXCL9
IFN-gamma
ANGPT1
TNFRSF21
VEGFA
CAIX
HO-1
CCL17
CXCL13
CASP-8
CSF-1
LAP TGF-beta-1
ANGPT2
IL5
IL7
IL12
FGF2
DCN
Gal-1
PTN
IL18
IL33
ADGRG1
PD-L2
VEGFR-2
CX3CL1
PDCD1
TNFRSF4
IL8
MCP-1
IL6
CXCL1
CXCL5
CCL4
CCL20
CCL3
MCP-3
MCP-2
CD40-L
CD28
IL2
KIR3DL1
CXCL12
MUC-16
FASLG
CD27
TIE2
TRAIL
CD244
KLRD1
LAG3
NOS3
TWEAK
PDGF subunit B
HGF
CD70
IL-1 alpha
ADA
TNFSF14
ARG1
IL4
IL12RB1
NCR1
LAMP3
CRTAM
CCL19
aCD47
Nivolumab
Combo

**Figure S10**

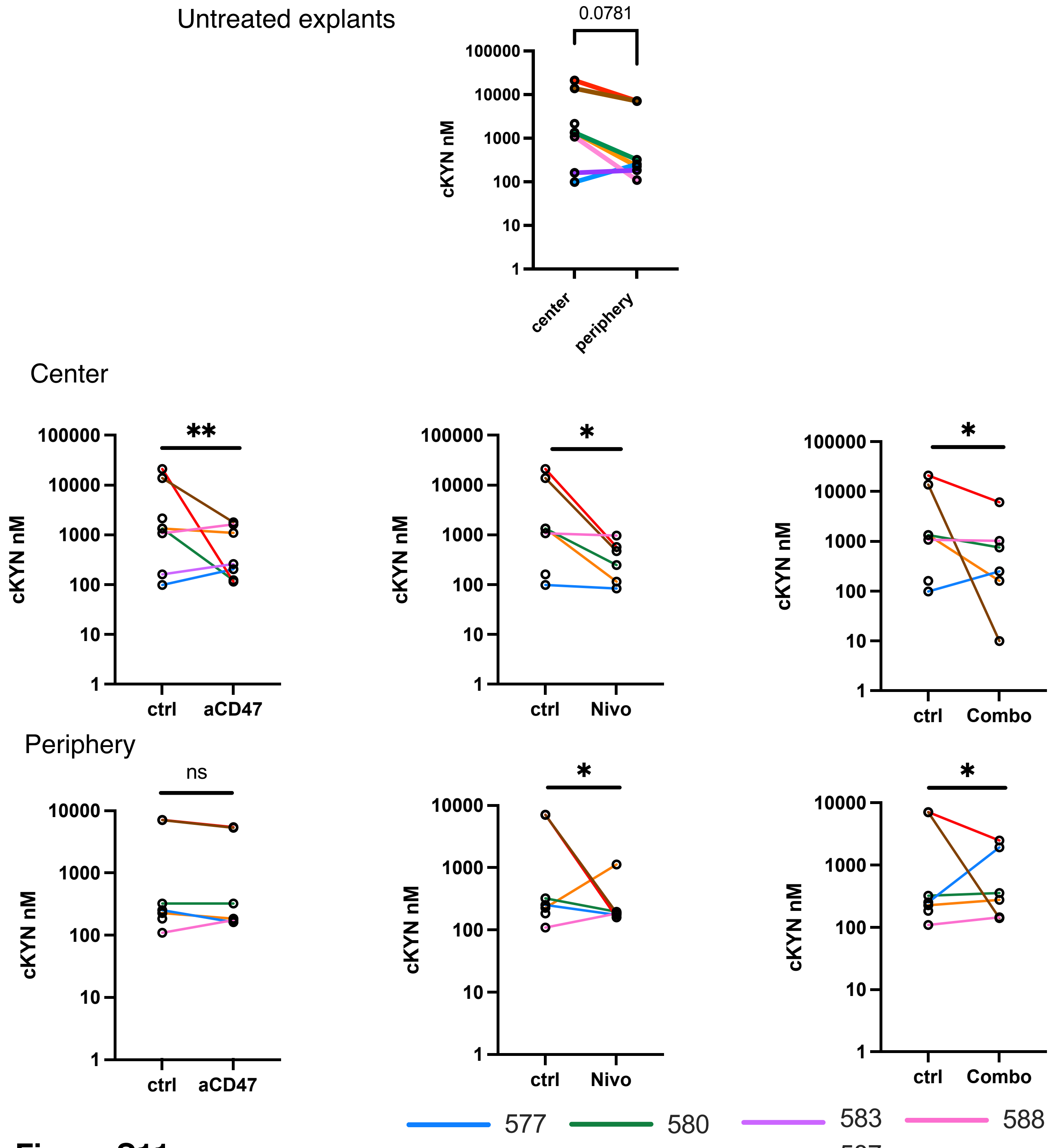
Untreated explants
0.0781
cKYN nM
100000
10000
1000
100
10
1
center
periphery
Center
**
*
*
ctrl
aCD47
Nivo
Combo
Periphery
ns
577
580
583
588
579
581
587

**Figure S11**

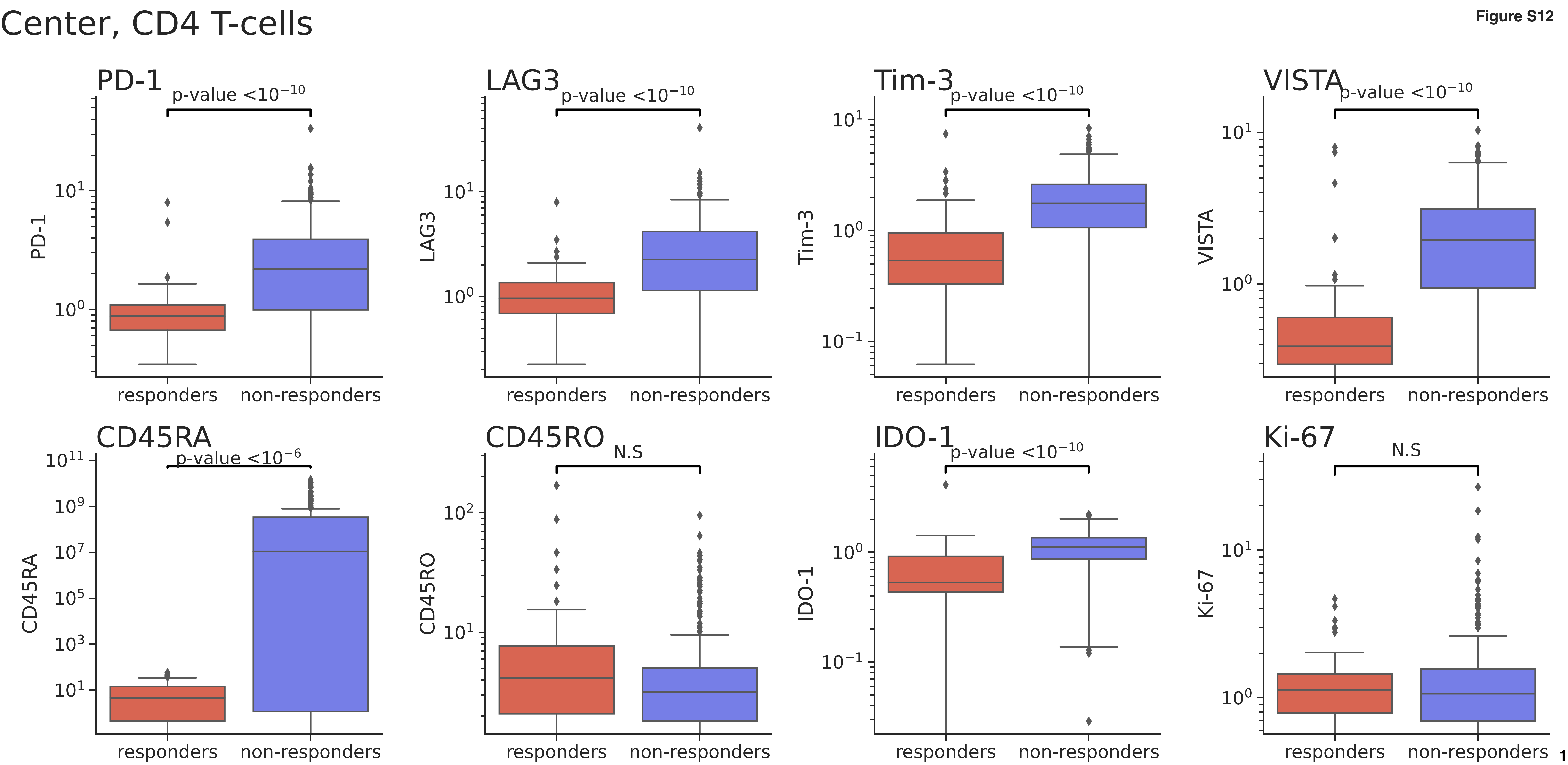
Center, CD4 T-cells
PD-1
p-value <$10^{-10}$
LAG3
p-value <$10^{-10}$
Tim-3
p-value <$10^{-10}$
VISTA
p-value <$10^{-10}$
CD45RA
p-value <$10^{-6}$
CD45RO
N.S
IDO-1
p-value <$10^{-10}$
Ki-67
N.S
responders
non-responders

Figure S12

Periphery, CD4 T-cells

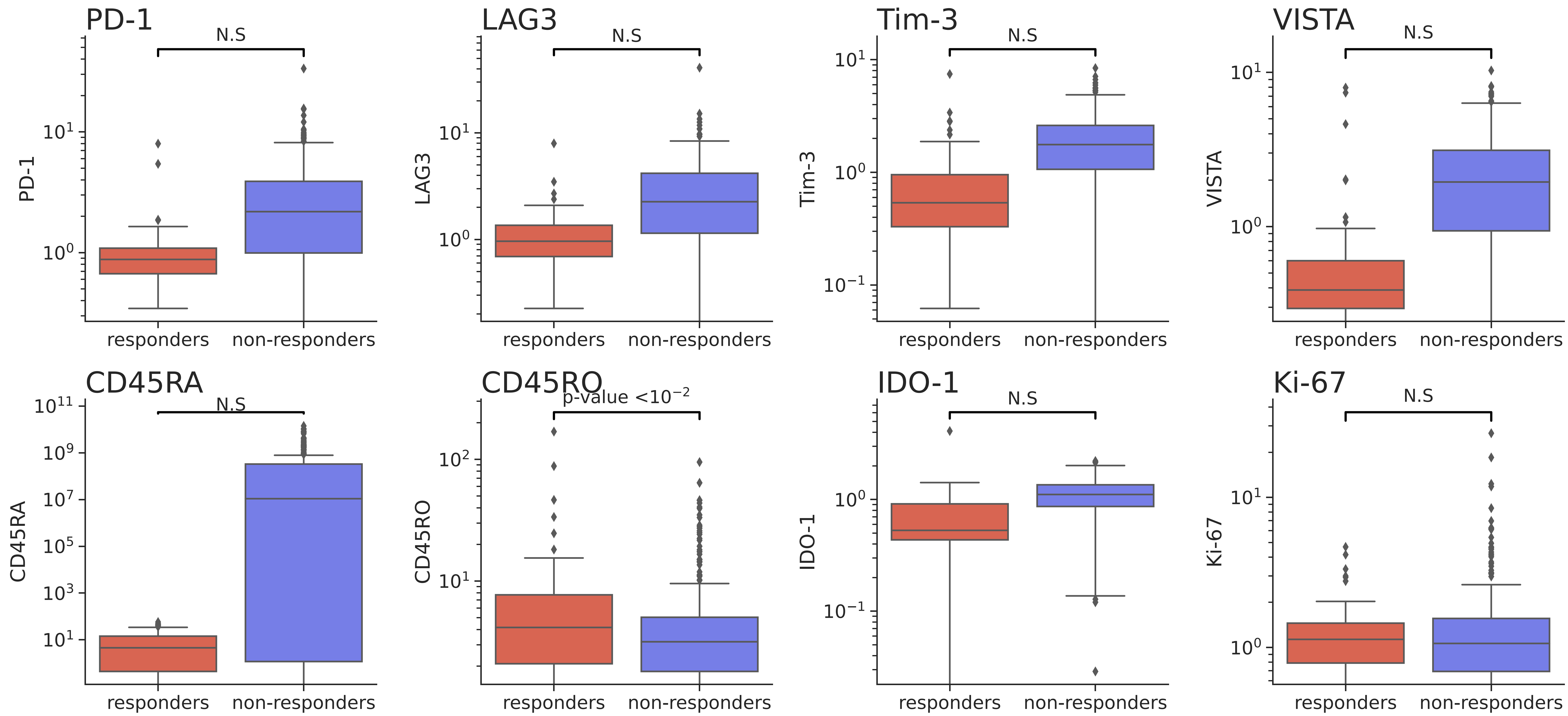

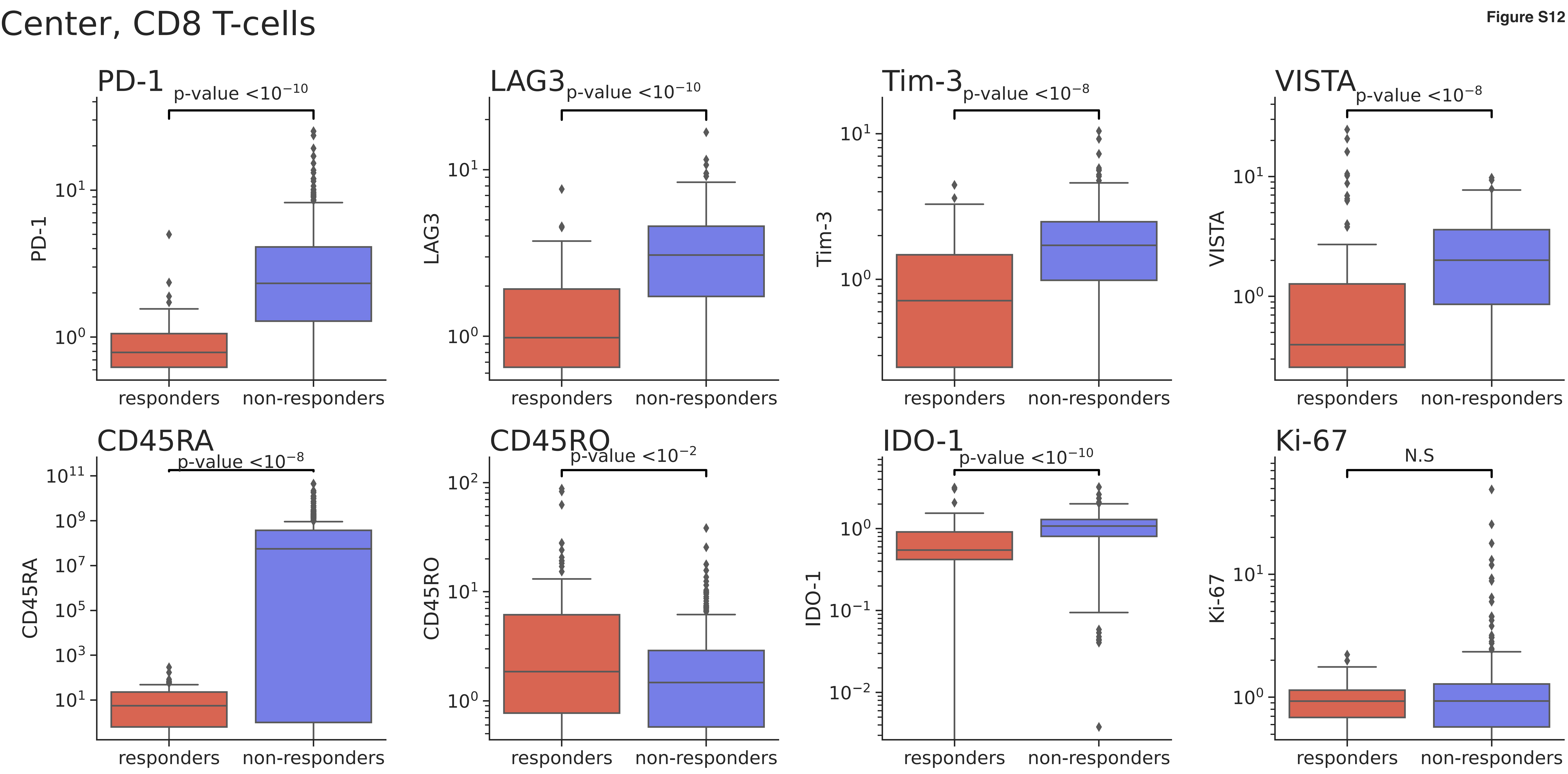

Center, CD8 T-cells
PD-1
p-value <$10^{-10}$
LAG3
p-value <$10^{-10}$
Tim-3
p-value <$10^{-8}$
VISTA
p-value <$10^{-8}$
CD45RA
p-value <$10^{-8}$
CD45RO
p-value <$10^{-2}$
IDO-1
p-value <$10^{-10}$
Ki-67
N.S
responders
non-responders

### Periphery, CD8 T-cells

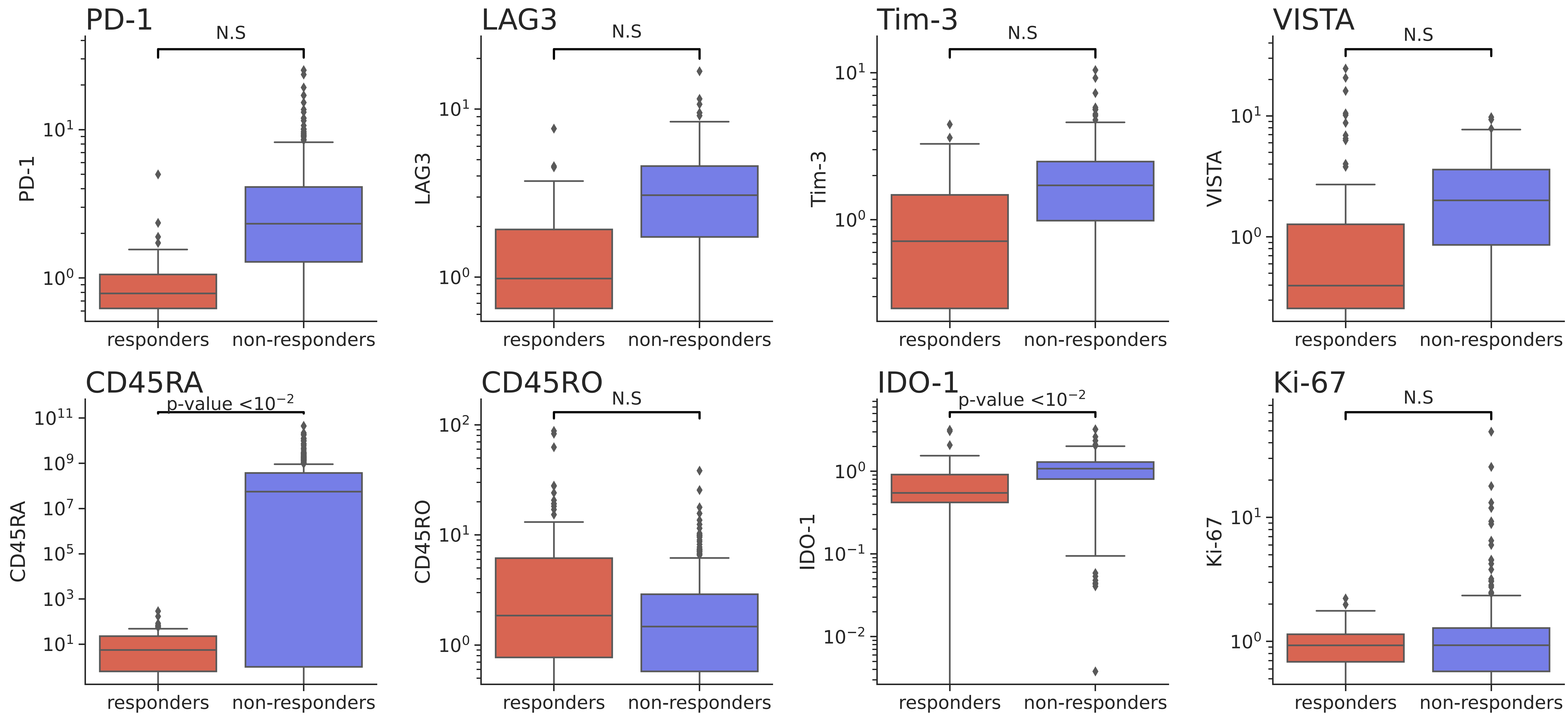

### Center, Lymphocytes

Figure S12

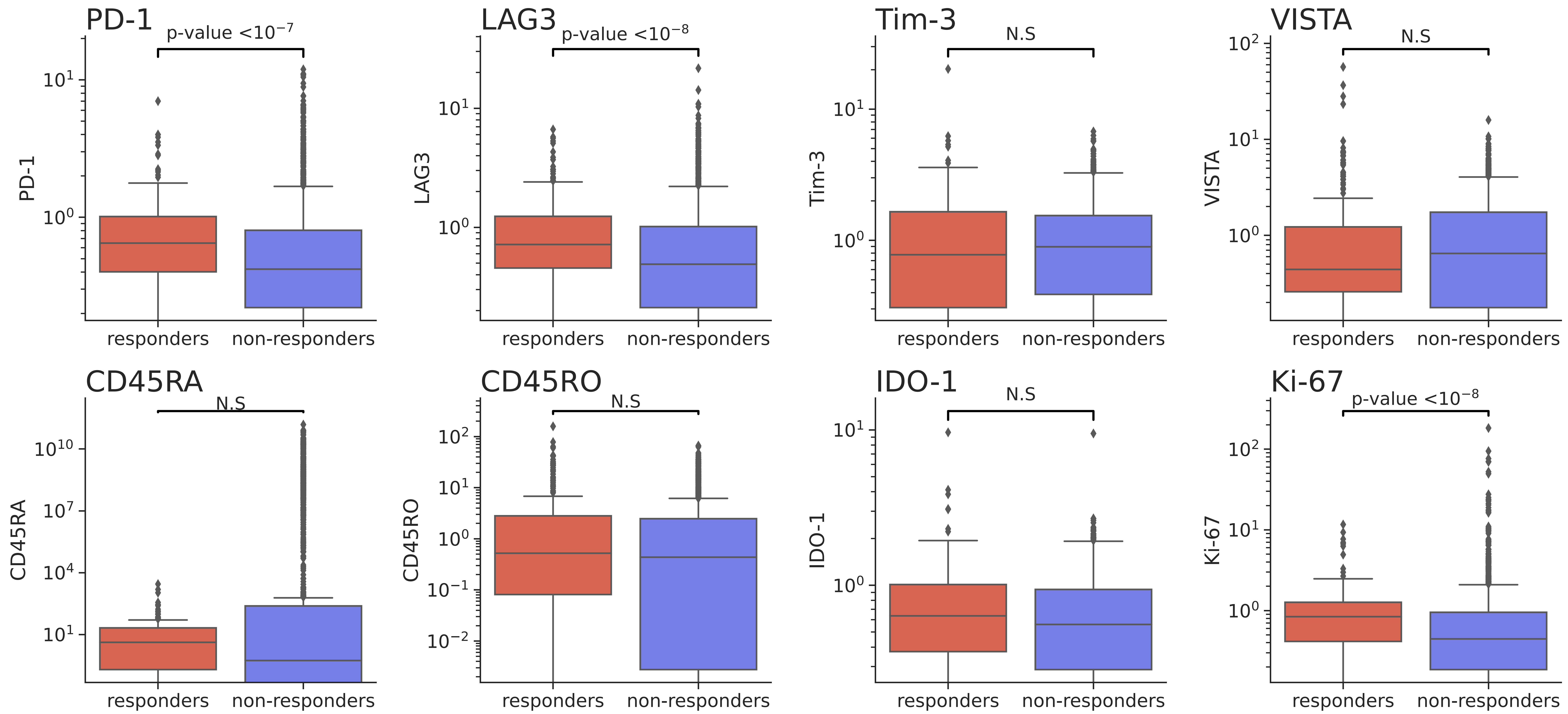

Figure S12

Periphery, Lymphocytes

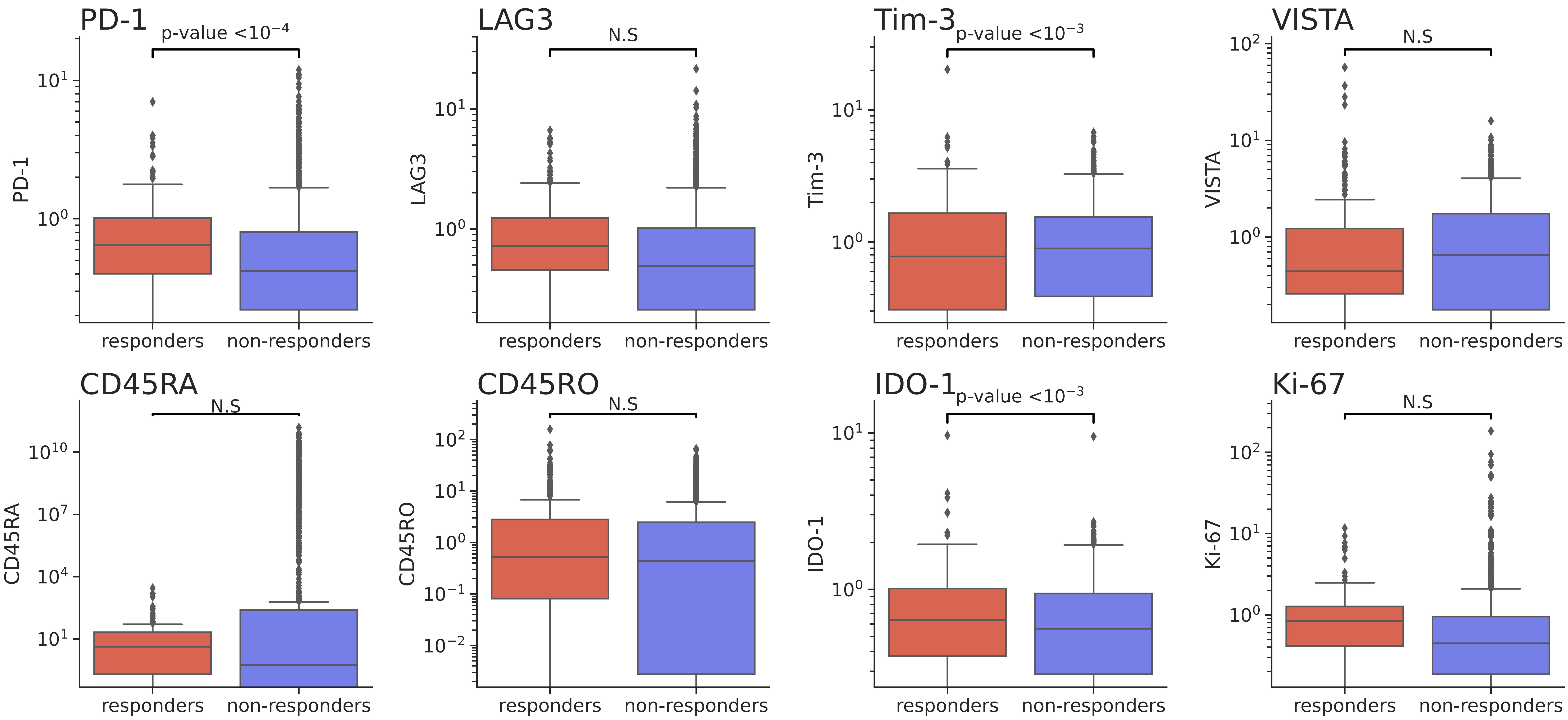

Figure S12

### Center, M1 macrophages

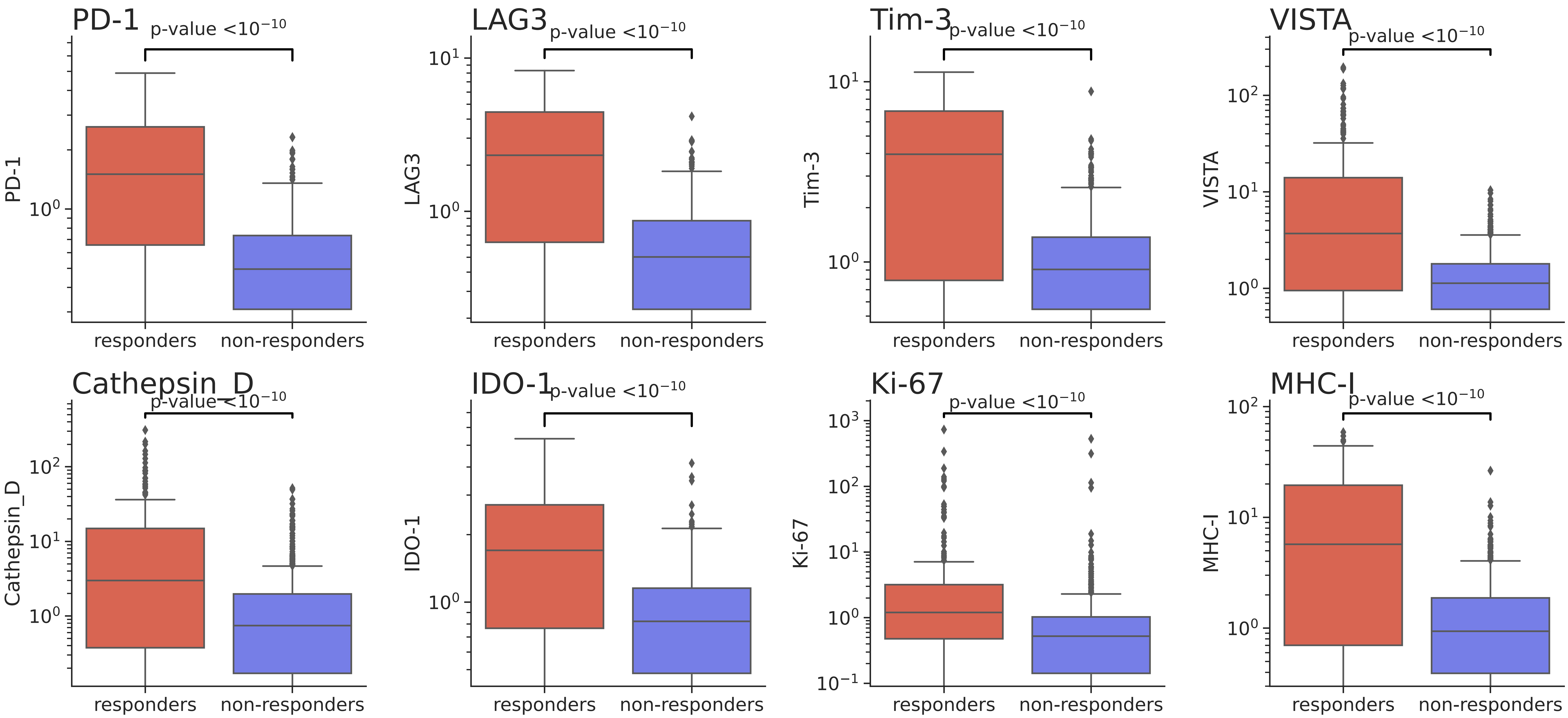

Figure S12

Periphery, M1 macrophages

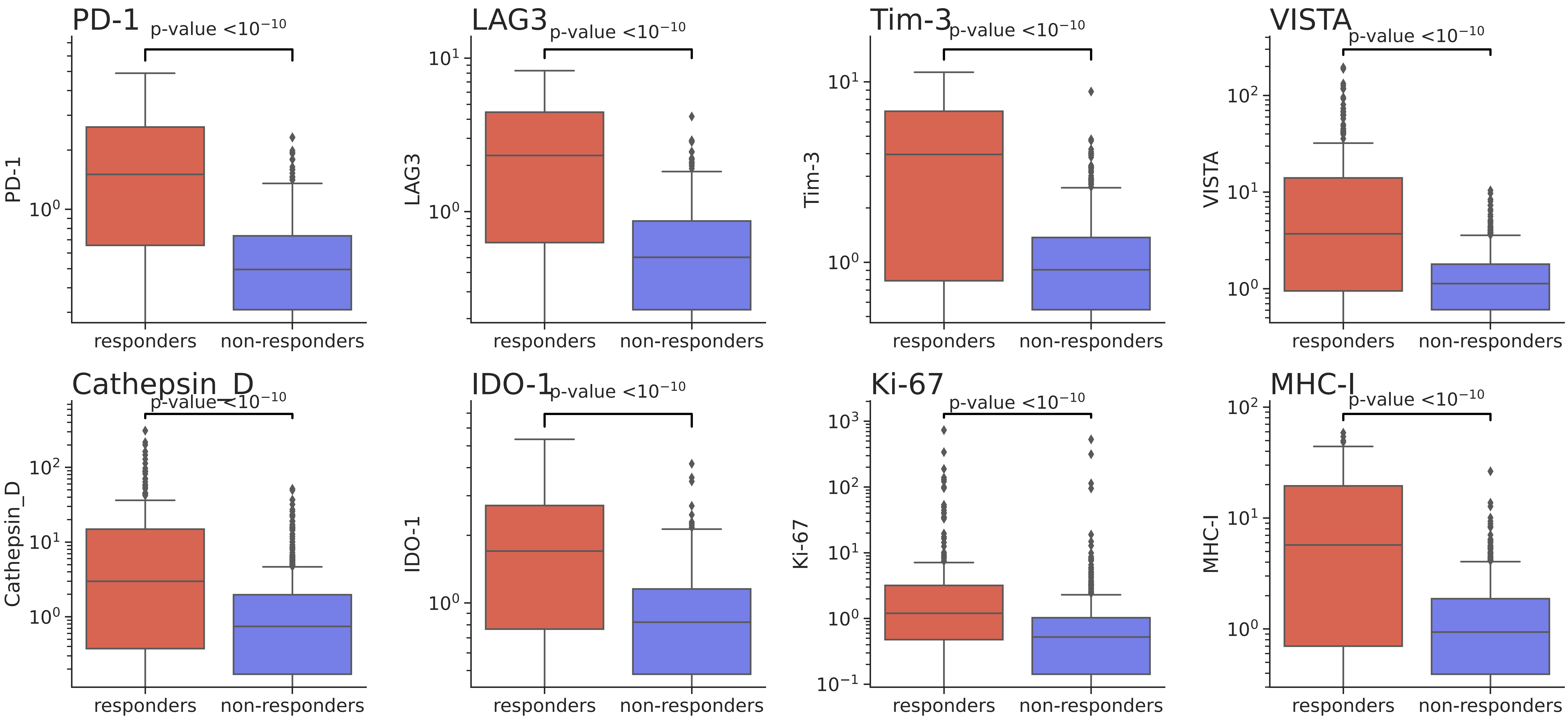

### Center, M2-like microglia

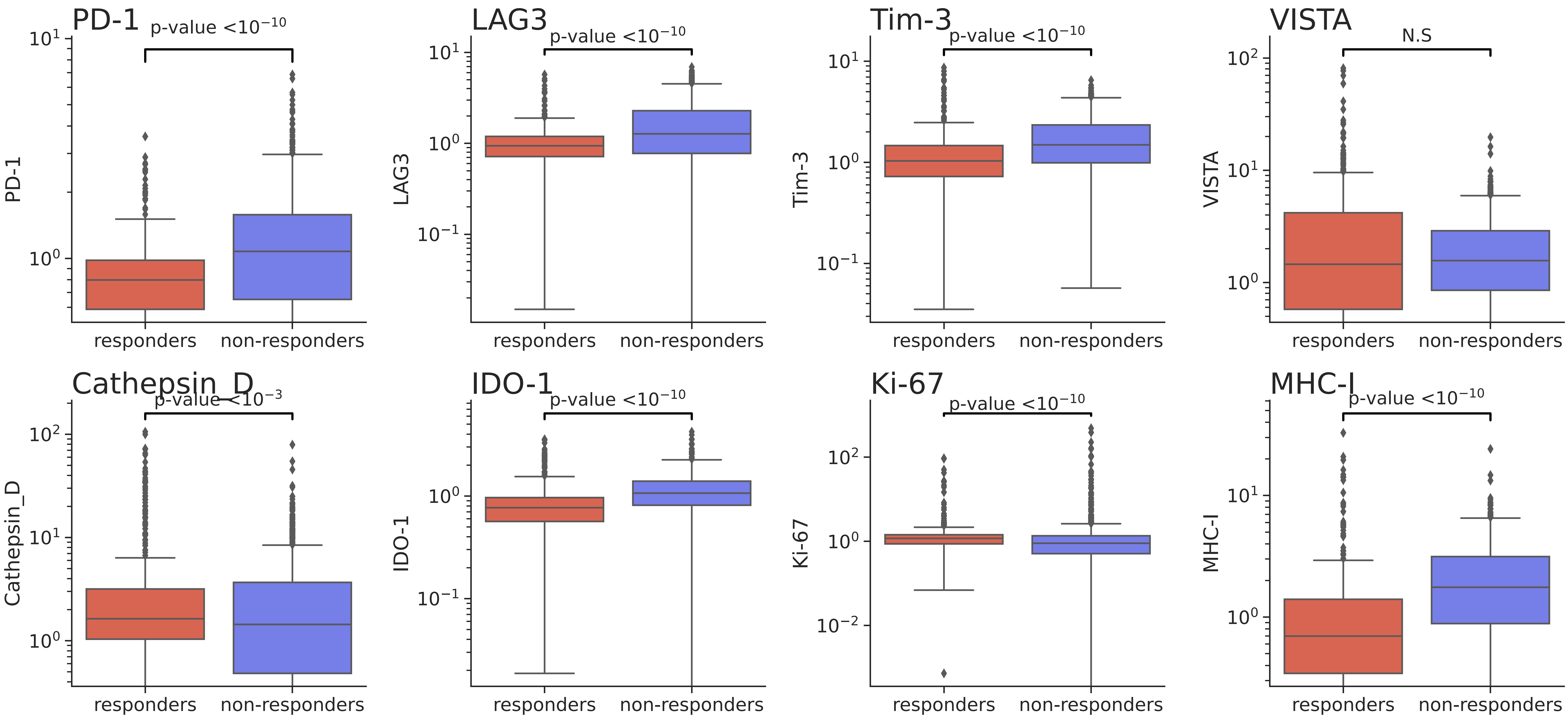

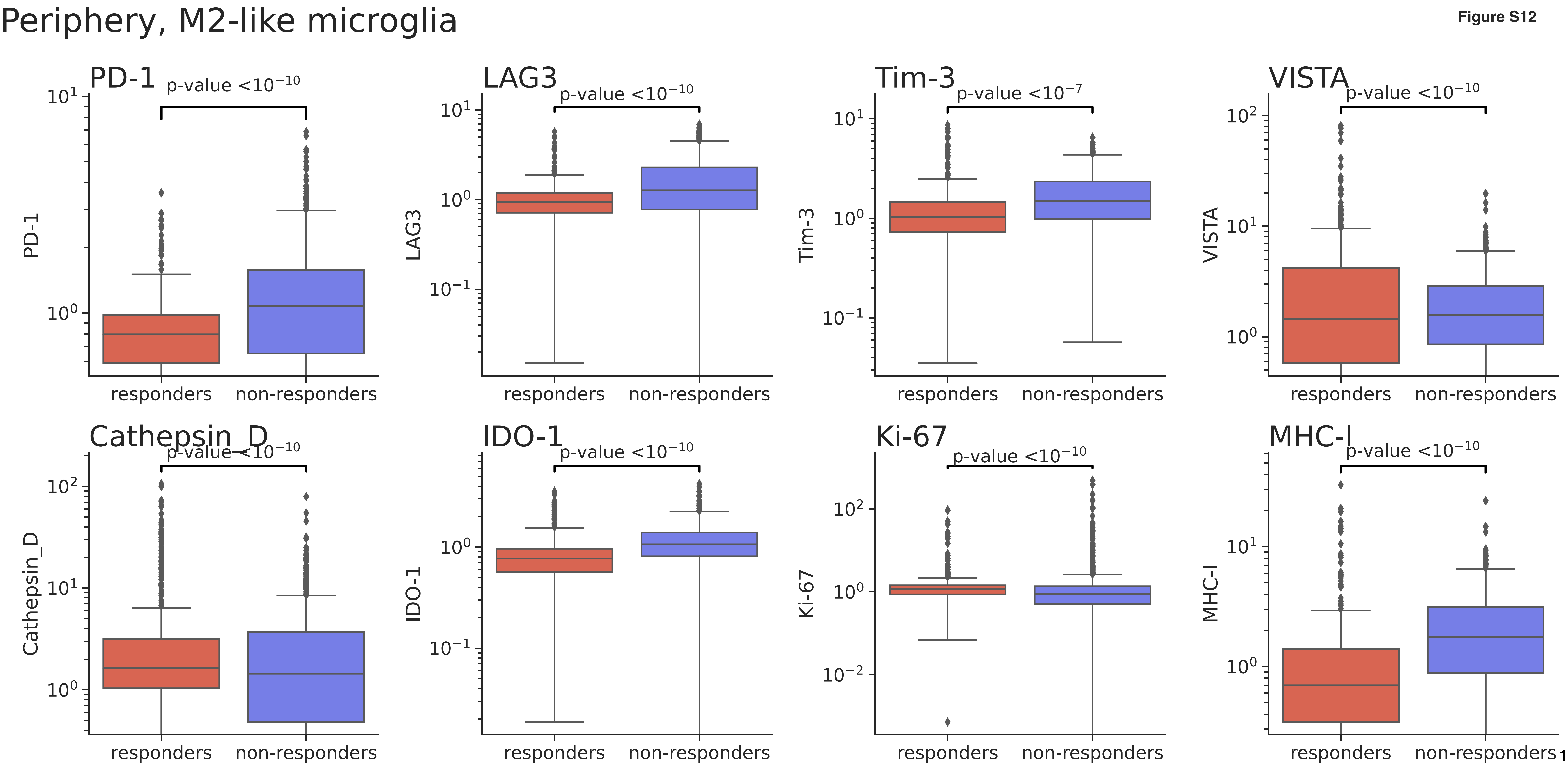
Periphery, M2-like microglia
PD-1
p-value <10−10
LAG3
p-value <10−10
Tim-3
p-value <10−7
VISTA
p-value <10−10
Cathepsin_D
p-value <10−10
IDO-1
p-value <10−10
Ki-67
p-value <10−10
MHC-I
p-value <10−10
responders
non-responders

Center, M2 macrophages

Figure S12

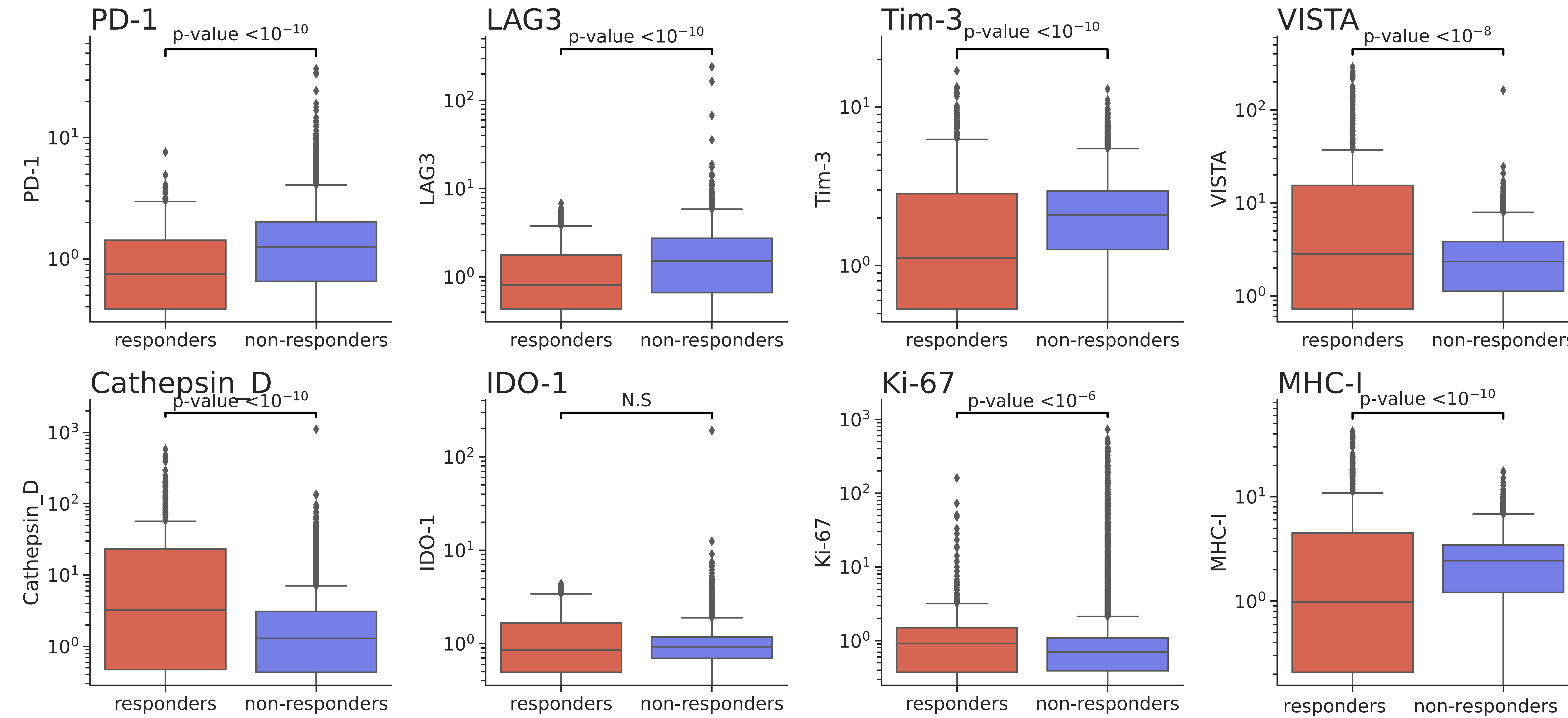

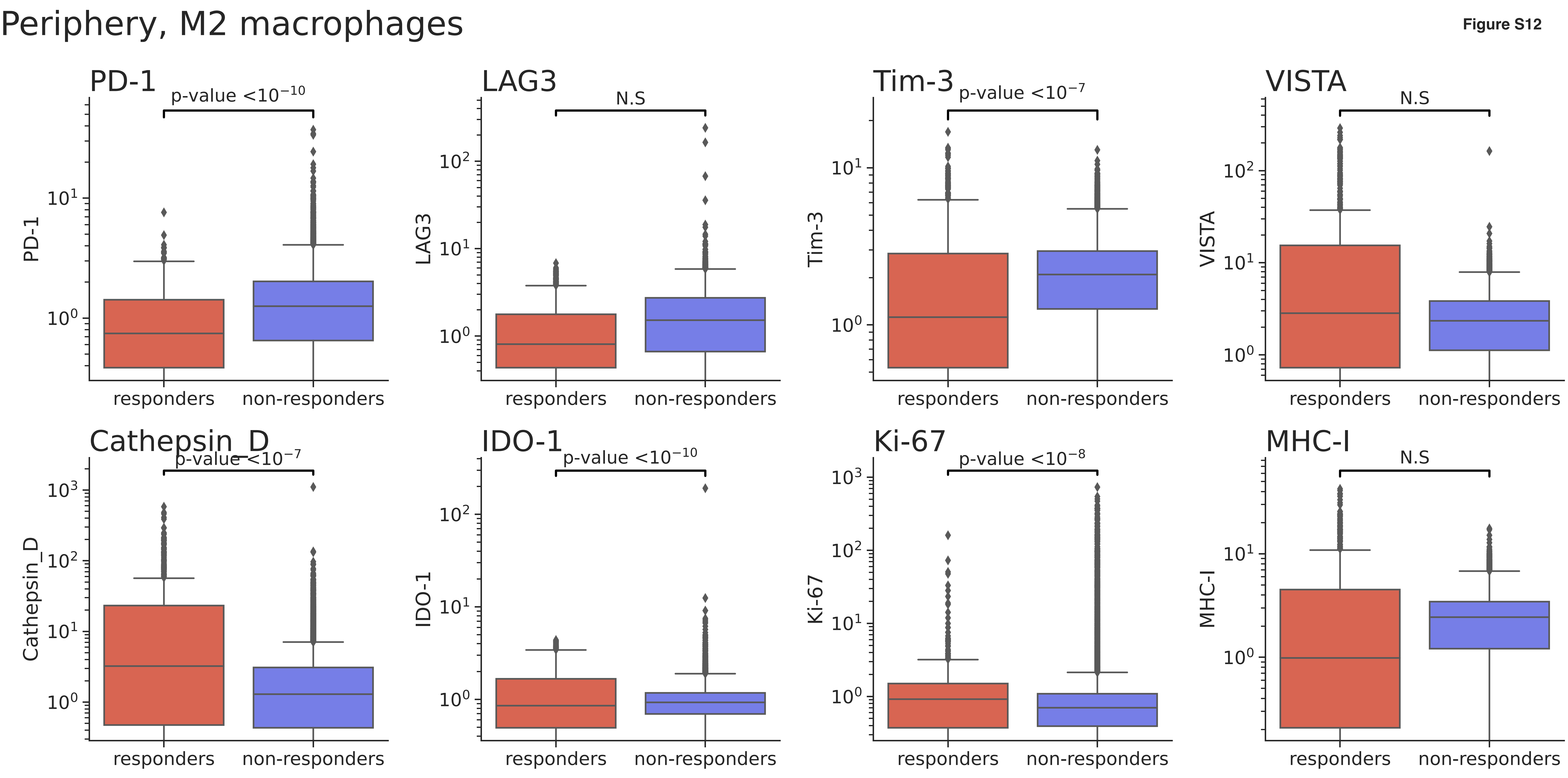

Periphery, M2 macrophages
PD-1
p-value <10^−10
LAG3
N.S
Tim-3
p-value <10^−7
VISTA
N.S
Cathepsin_D
p-value <10^−7
IDO-1
p-value <10^−10
Ki-67
p-value <10^−8
MHC-I
N.S
responders
non-responders

Figure S12

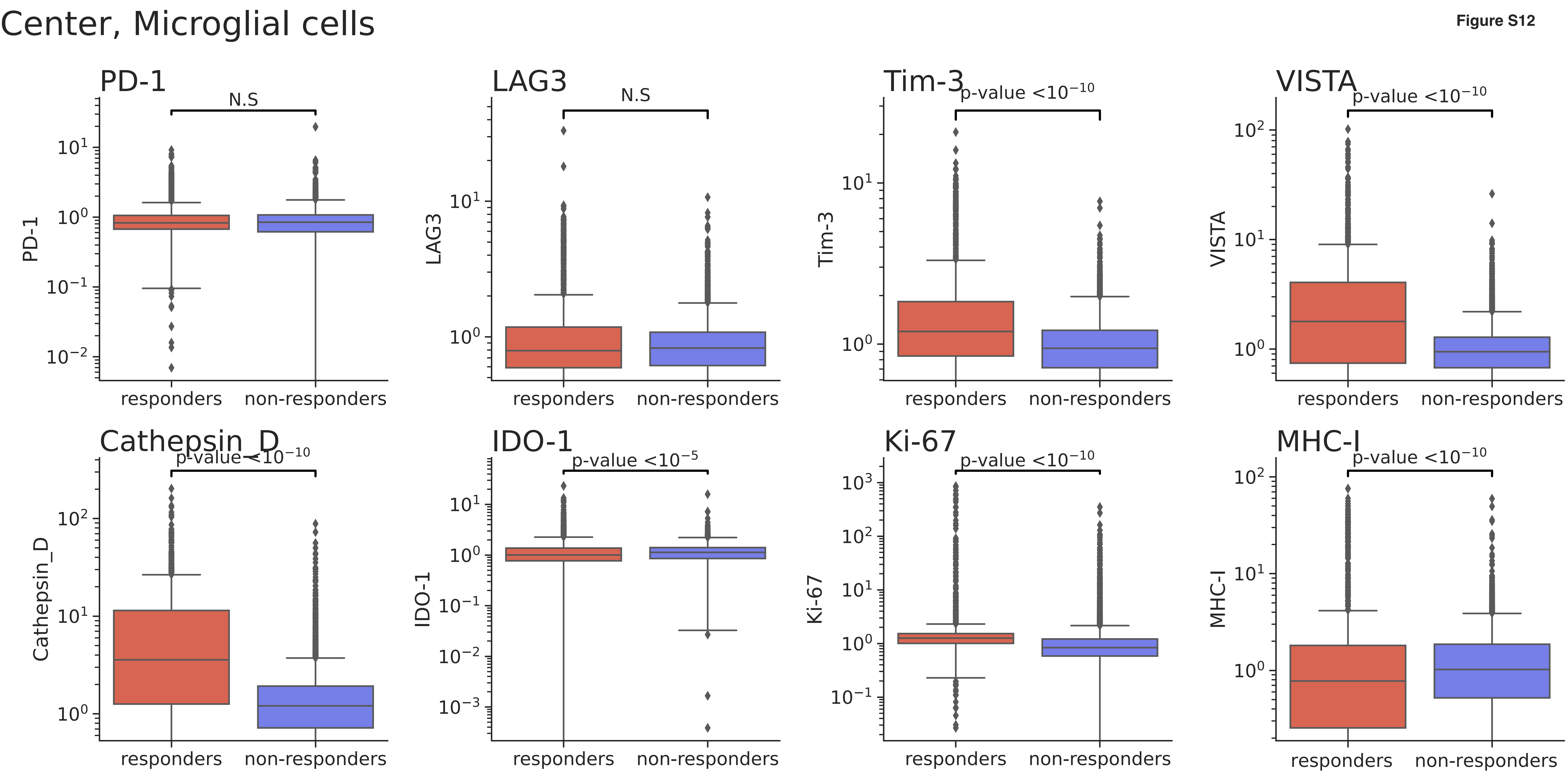

Center, Microglial cells
PD-1
N.S
LAG3
N.S
Tim-3
p-value <10^−10
VISTA
p-value <10^−10
Cathepsin_D
p-value <10^−10
IDO-1
p-value <10^−5
Ki-67
p-value <10^−10
MHC-I
p-value <10^−10
responders
non-responders

Periphery, Microglial cells

Figure S12

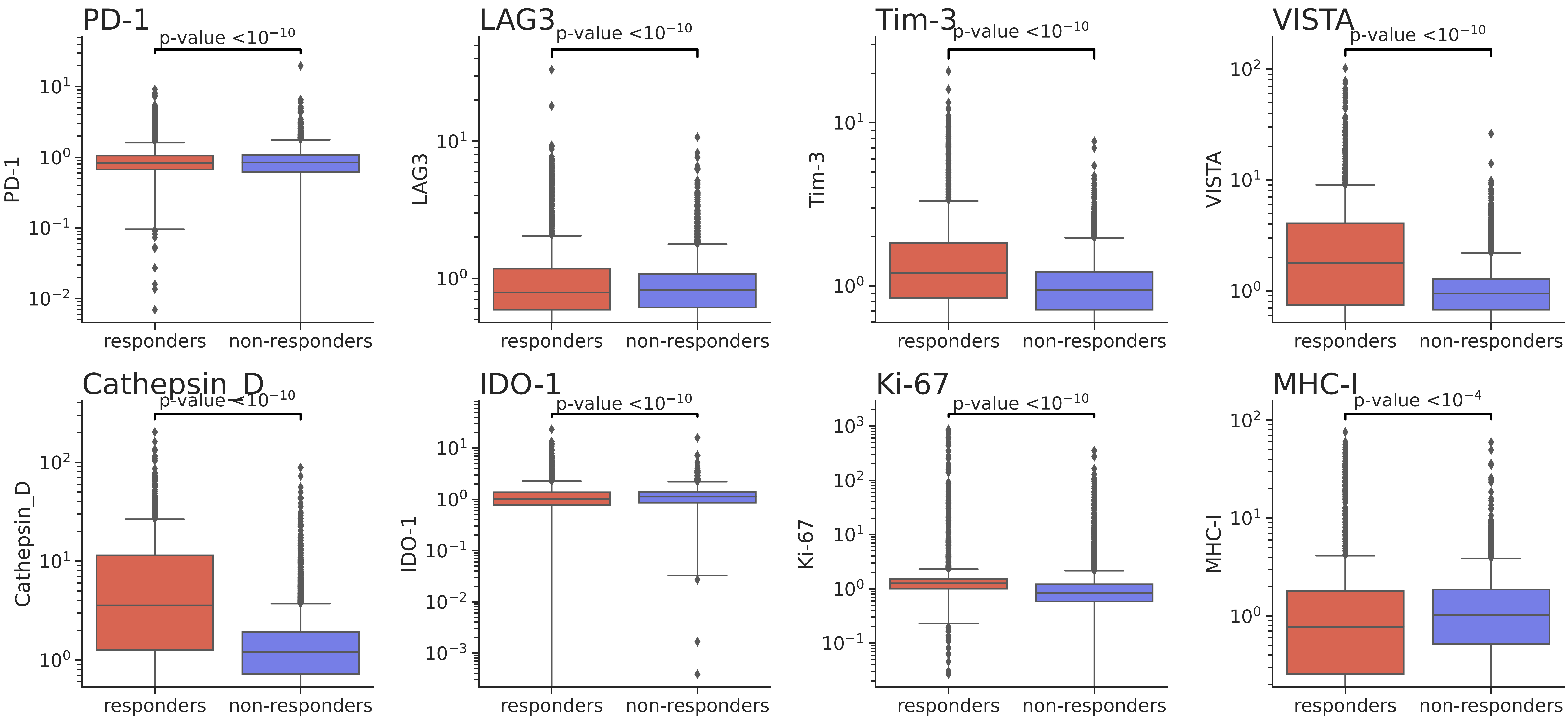

### Center, tumor cells/astrocytes

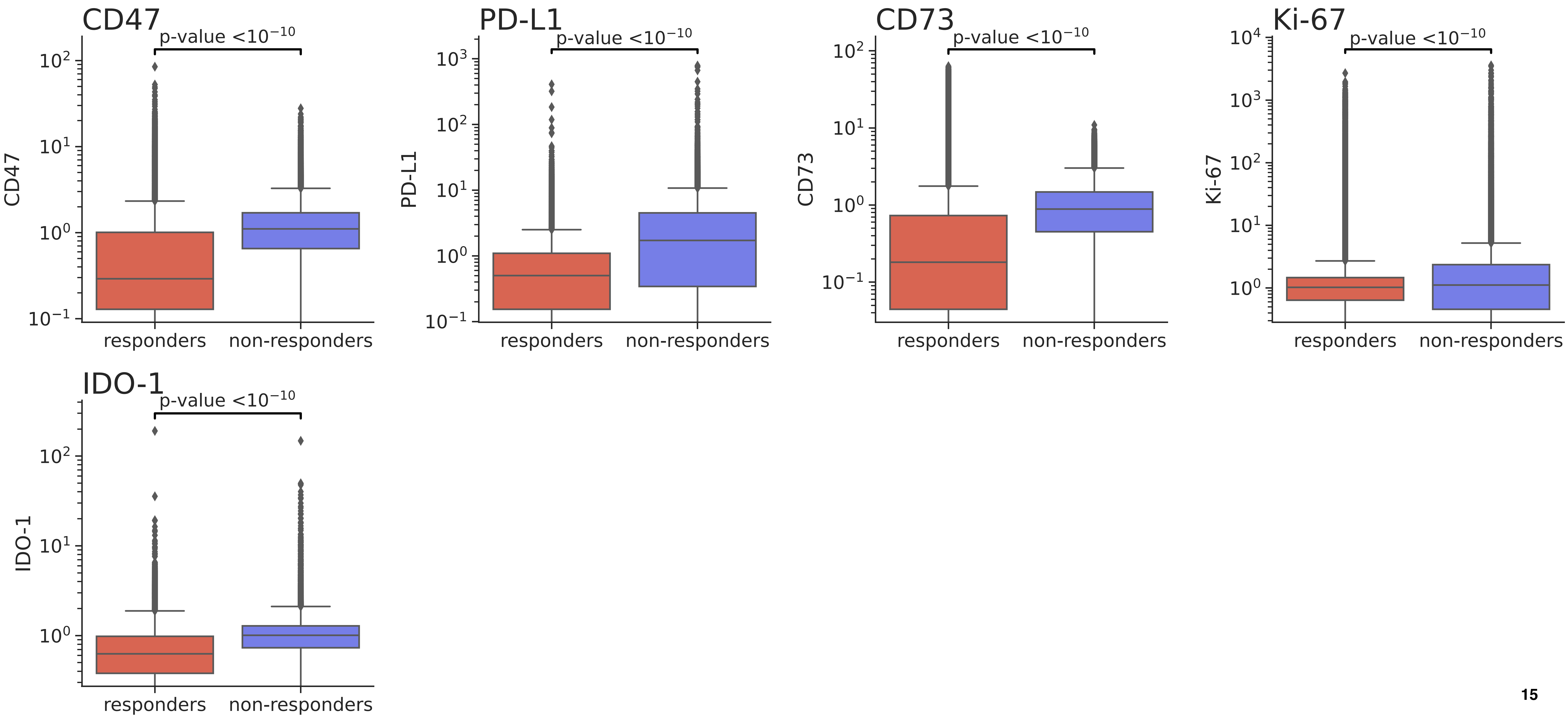

### Periphery, tumor cells/astrocytes

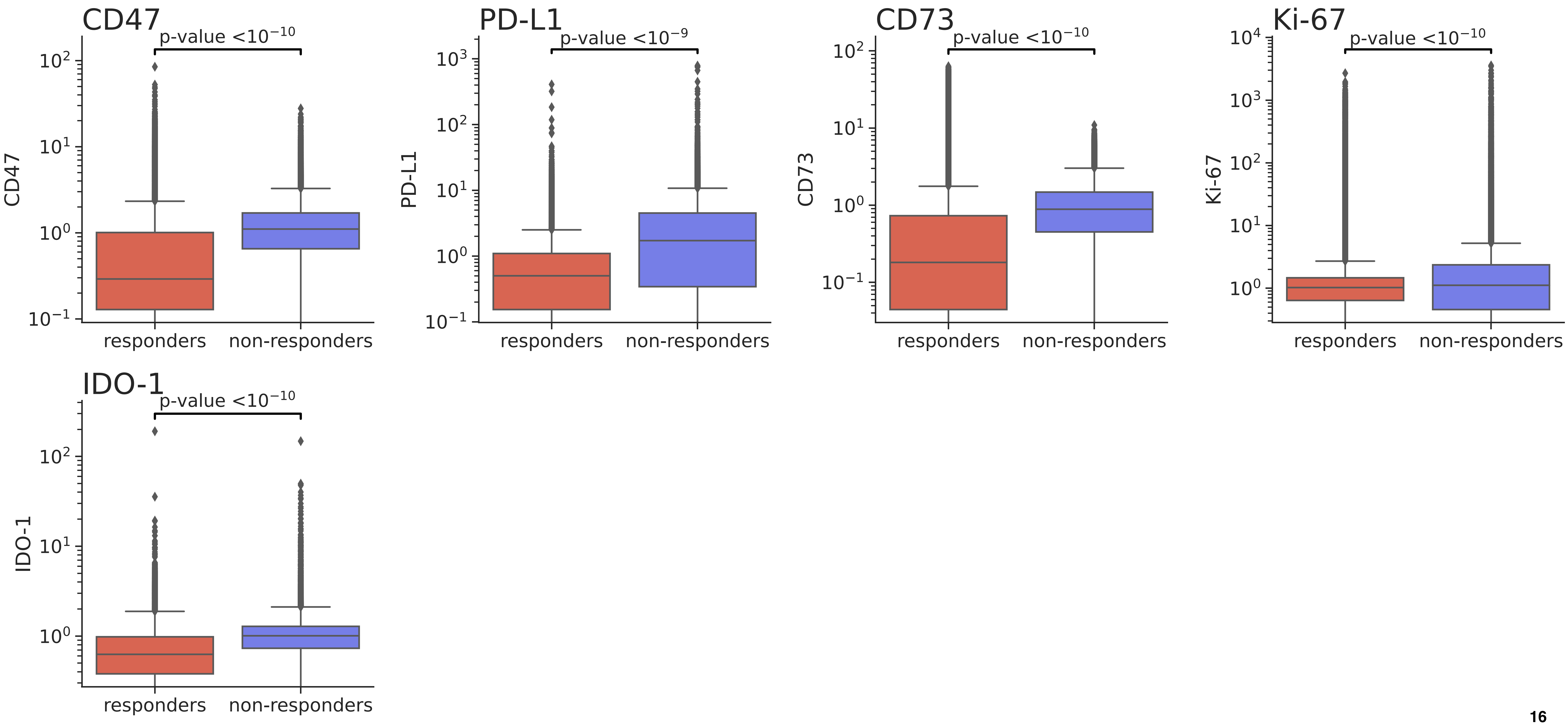
