## Supplementary material for "Immunotherapy of glioblastoma explants induces interferon-γ responses and spatial immune cell rearrangements in tumor center, but not periphery": Suppl Table 2

### Table S1

#### Immunotherapy of glioblastoma explants induces interferon-γ responses and spatial immune cell rearrangements in tumor center but not periphery

#### Acta Neuropathologica

Tala Shekarian, Carl P. Zinner, Ewelina M. Bartoszek, Wandrille Duchemin, Anna T. Wachnowicz, Sabrina Hogan, Manina M. Etter, Julia Flammer, Chiara Paganetti, Tomás A. Martins, Philip Schmassmann, Steven Zanganeh, Francois Le Goff, Manuele G. Muraro, Marie-Françoise Ritz, Darci Phillips, Salil S. Bhate, Graham L. Barlow, Garry P. Nolan,
Christian M. Schürch[1, 2, 3,*] and Gregor Hutter[4,5,*]

[1]Department of Pathology and Neuropathology, University Hospital and Comprehensive Cancer Center Tübingen, Tübingen, Germany

[2]Department of Microbiology & Immunology, Stanford University School of Medicine, Stanford, CA, USA

[3]Department of Pathology, Stanford University School of Medicine, Stanford, CA, USA

[4]Brain Tumor Immunotherapy Lab, Department of Biomedicine, University of Basel, Basel, Switzerland

[5]Department of Neurosurgery, University Hospital Basel, Basel, Switzerland

#### Baseline Parameters

| ID | Sex | Age | DG | WHO | Survival |
|---|---|---|---|---|---|
| 577 | M | 73 | GBM | IV | 500 |
| 579 | F | 58 | GBM | IV | 505 |
| 580 | M | 39 | GBM | IV | 651 |
| 581 | M | 59 | GBM | IV | >700* |
| 583 | M | 71 | GBM | IV | 227 |
| 587 | F | 69 | GBM | IV | 470 |
| 588** | F | 78 | GBM | IV | 204 |

***Pretreated with radio-chemotherapy*

** still alive*

#### Genetical characterization

| ID | Subclass | *IDH* | MGMT | *TERT* | *EGFR* |
|---|---|---|---|---|---|
| 577 | MES | WT | UNM | WT | 2n |
| 579 | MES | WT | UNM | WT | 2n |
| 580 | RTK II CL | WT | MET | G228A | 2n |
| 581 | RTK II CL | WT | MET | G228A | >2 |
| 583 | RTK II CL | WT | UNM | G228A | 2n |
| 587 | MES | WT | UNM | WT | 2n |
| 588 | MES | WT | MET | G228A | 2n |

#### Tissue availability

| ID | *Explant (D7) tissue* | | *Cytokines* |
|---|---|---|---|
| | Center | Periphery | |
| 577 | y | y | y |
| 579 | y | y | y |
| 580 | y | y | y |
| 581 | y | NA | y |
| 583 | y | y | y |
| 587 | y | NA | y |
| 588 | y | y | y |
